# Analysis of SARS-CoV-2 spike glycosylation reveals shedding of a vaccine candidate

**DOI:** 10.1101/2020.11.16.384594

**Authors:** Juliane Brun, Snežana Vasiljevic, Bevin Gangadharan, Mario Hensen, Anu V. Chandran, Michelle L. Hill, J.L. Kiappes, Raymond A. Dwek, Dominic S. Alonzi, Weston B. Struwe, Nicole Zitzmann

## Abstract

Severe acute respiratory syndrome coronavirus 2 is the causative pathogen of the COVID-19 pandemic which as of Nov 15, 2020 has claimed 1,319,946 lives worldwide. Vaccine development focuses on the viral trimeric spike glycoprotein as the main target of the humoral immune response. Viral spikes carry glycans that facilitate immune evasion by shielding specific protein epitopes from antibody neutralisation. Immunogen integrity is therefore important for glycoprotein-based vaccine candidates. Here we show how site-specific glycosylation differs between virus-derived spikes and spike proteins derived from a viral vectored SARS-CoV-2 vaccine candidate. We show that their distinctive cellular secretion pathways result in different protein glycosylation and secretion patterns, which may have implications for the resulting immune response and future vaccine design.

---

Severe acute respiratory syndrome coronavirus 2 (SARS-CoV-2), the causative agent of coronavirus disease 2019 (COVID-19) can induce fever, severe respiratory illness, and various multi-organ disease manifestations [1]. The virus enters host cells by binding to angiotensin-converting enzyme 2 (ACE2) using its extensively glycosylated spike (S) protein [2], [3]. The S glycoprotein is a class I fusion protein, comprising two functional subunits. The S1 subunit is responsible for ACE2 receptor binding and the S2 subunit initiates membrane fusion between the virus particle and host cell. S protein synthesis in the endoplasmic reticulum (ER) of an infected cell is accompanied by co-translational addition of pre-assembled N-glycans to its 22 N-glycosylation sites [4].

After trimerisation and initial N-glycan processing in the ER by ER resident sugar modifying enzymes, S proteins travel as membrane anchored trimers to the ER-Golgi intermediate compartment (ERGIC) where they are incorporated into the membranes of viruses budding into the ERGIC lumen [5], [6]. S trimers protrude from the viral surface while individual viruses move along inside the lumina of cis-, medial- and trans-Golgi, where their N-glycans are extensively processed and further modified further by Golgi resident glycosylation enzymes. O-glycans are also added in the Golgi, starting with the addition of GalNAc residues via GalNAc transferase, which can be elongated similarly to N-glycans across the Golgi stack. In the trans-Golgi, S trimers encounter the host protease furin which cleaves between S1 and S2 [7], [8], leaving the subunits on S trimers non-covalently associated before the virus is secreted via lysosomes into the extracellular surrounding [9].

Host-derived glycosylation plays many important roles in viral pathobiology, including mediating viral protein folding and stability, as well as influencing viral tropism and immune evasion [10].The trimeric spikes protruding from viruses are key targets of the natural immune response [11]. Neutralising antibodies binding to these spikes, especially to S1, prevent cellular uptake of viruses by the host. Consequently, most vaccine design efforts focus on the S protein. The surface of each trimeric spike displays up to 66 N-linked glycans and an undefined number of O-linked glycans [12]. Understanding how SARS-CoV-2 exploits glycosylation on native S proteins will help guide rational vaccine design, as glycans enable immune evasion by shielding underlying immunogenic protein epitopes from antibody neutralisation, as also observed for other coronaviruses [13], [14]. In other instances, glycans constitute functional epitopes in immune recognition [15], further highlighting the need for molecular mimicry between viruses and vaccines that are designed to prime the immune system by eliciting neutralising antibodies.

Importantly, several COVID-19 vaccine candidates are based on viral vectors encoding SARS-CoV-2 S protein, including ChAdOx1 nCoV-19 (AZD1222; Folegatti et al., 2020; van Doremalen et al., 2020). To compare virus-derived S protein glycosylation with that of a viral vector vaccine candidate, we grew SARS-CoV-2 (England/02/2020 strain) in Calu-3 lung epithelial cells, harvested the virus containing supernatant, and immunopurified detergent-solubilised spike using a CR3022 antibody column (**Figure 1A**). Immunopurified material was analysed by SDS-PAGE (**Figure 1B**) and the protein bands corresponding to S1 (S1_virus_) and S2 were excised from the gel and confirmed by mass spectrometry (**Supplementary Figure 1A**). S2 levels were in insufficient amounts for additional glycan/glycoproteomics analysis. Quantitative analysis of S1_virus_ released N-glycans by UPLC (**Figure 1C**) showed a predominant population of complex-type N-glycans (79%) with 21% oligomannose and/or hybrid structures. Comparing these values to a soluble recombinant trimeric form of S (S_recombinant trimer_), which has been engineered to abolish furin cleavage and therefore also contains S2 N-glycans [18], revealed S_recombinant trimer_ to carry only 11% oligomannose/hybrid and 89% complex N-glycans (**Figure 1D**). This observation is significant as it indicates large-scale differences in glycan processing, a complex pathway that is influenced by high glycan density and local protein architecture, both of which can sterically impair glycan maturation (**Figure 1E**). Changes in glycan maturation, resulting in the presence of oligomannose-type glycans, can be a sensitive reporter of native-like protein architecture [19], [20], and is also an important indicator for quality control and efficacy of different immunogens [21].

**Figure 1:**
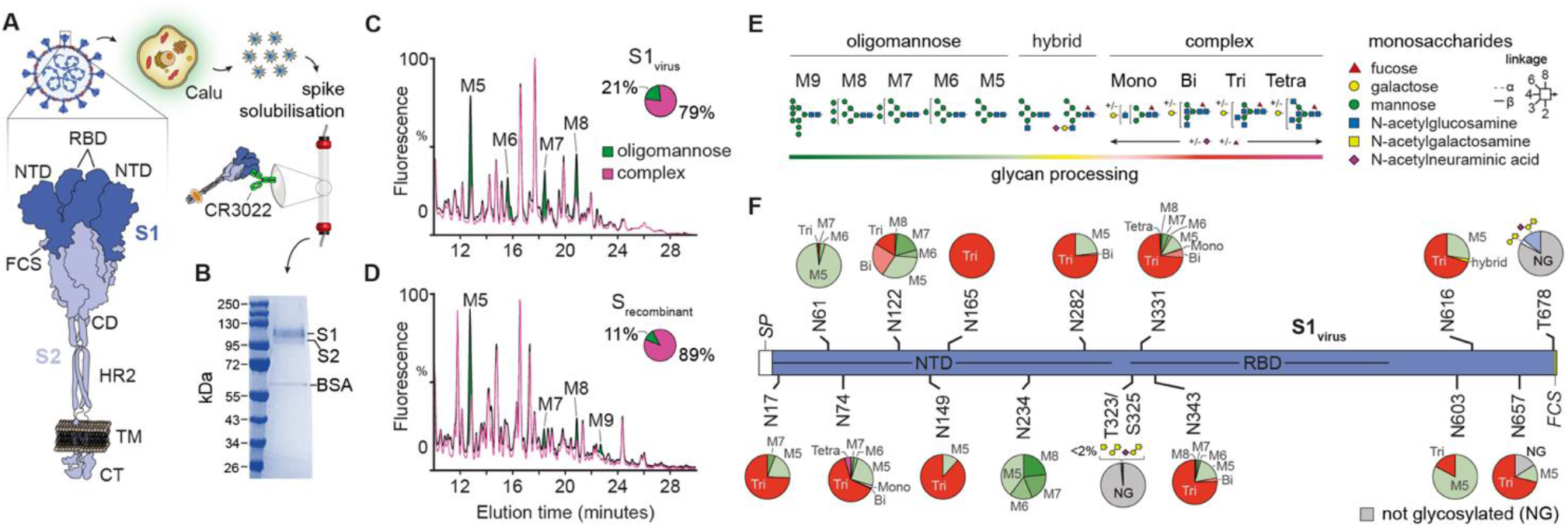
Purification and glycan analysis of the SARS-CoV-2 spike glycoprotein. (**A**) Schematic representation of spike purification from SARS-CoV-2 infected Calu-3 cells by immunoaffinity purification using the S1 targeting CR3022 antibody; (**B**) SDS-PAGE showing the presence of S1 and S2 subunits of virus-derived spike; (**C** and **D**) Quantitative UPLC N-glycan analysis showing the distribution of oligomannose and complex-type glycans on S1_virus_ (**C**) and S_recombinant trimer_ (**D**); (**E**) N-glycan maturation showing colour coding for degree of glycan processing from oligomannose (green) to hybrid (yellow) to complex (purple); (**F**) Quantitative site-specific N- and O-glycosylation by bottom-up glycoproteomics of S1_virus_. Pie charts depict the degree of N-glycan processing depicted in (**E**).

To pinpoint where, and the extent to which, differences in glycan processing occur, we performed a quantitative site-specific N- and O-glycosylation analysis of S1_virus_ (**Figure 1F**) and S_recombinant trimer_ (**Supplementary Figure 2**) by mass spectrometry. We detected glycopeptides for all 13 potential N-glycosylation sites in S1 and importantly, we found that S1 N-glycan processing is comparable between virus and recombinant material, excluding the possibility that differences in glycan processing observed by UPLC are outweighed by the presence of the S2 subunit on S_recombinant trimer._ Looking closer at S1_virus_, we observed three N-glycan sites, N61, N234 and N603, that are predominantly occupied by underprocessed oligomannose structures and are likely shielded by the quaternary protein structure. This is in contrast to previously reported N-glycan analysis on virus derived S, where N61 carried mostly complex-type (with some oligomannose) glycans; N234 was a mixture of oligomannose, hybrid and complex structures and N603 was mostly complex [22]. We found that the remaining sites on S1_virus_ were either occupied almost entirely by tri-antennary N-glycans (N149 and N165), or by a mixture of tri-antennary complex plus oligomannose (namely Man_5_GlcNAc_2_, i.e. M5) structures. We did not detect any O-linked glycosylation at T232/S325 on S_recombinant trimer_, an observation that is variably reported among recombinant S or S1 material [22]–[25]. However, we identified O-glycosylation at T678 on S1_virus,_ which was absent on S_recombinant trimer_. This is particularly informative, indicating this domain on S1 _virus_ is more accessible to GalNAc-transferases in the Golgi and that the viral spike is possibly configured in a more open or flexible trimeric state than the recombinant, stabilised spike. We also observed the presence of SARS-CoV-2 nucleoprotein and SARS-CoV-2 membrane protein at lower levels in the immunopurified material (**Supplementary Table 1**).

**Figure 2:**
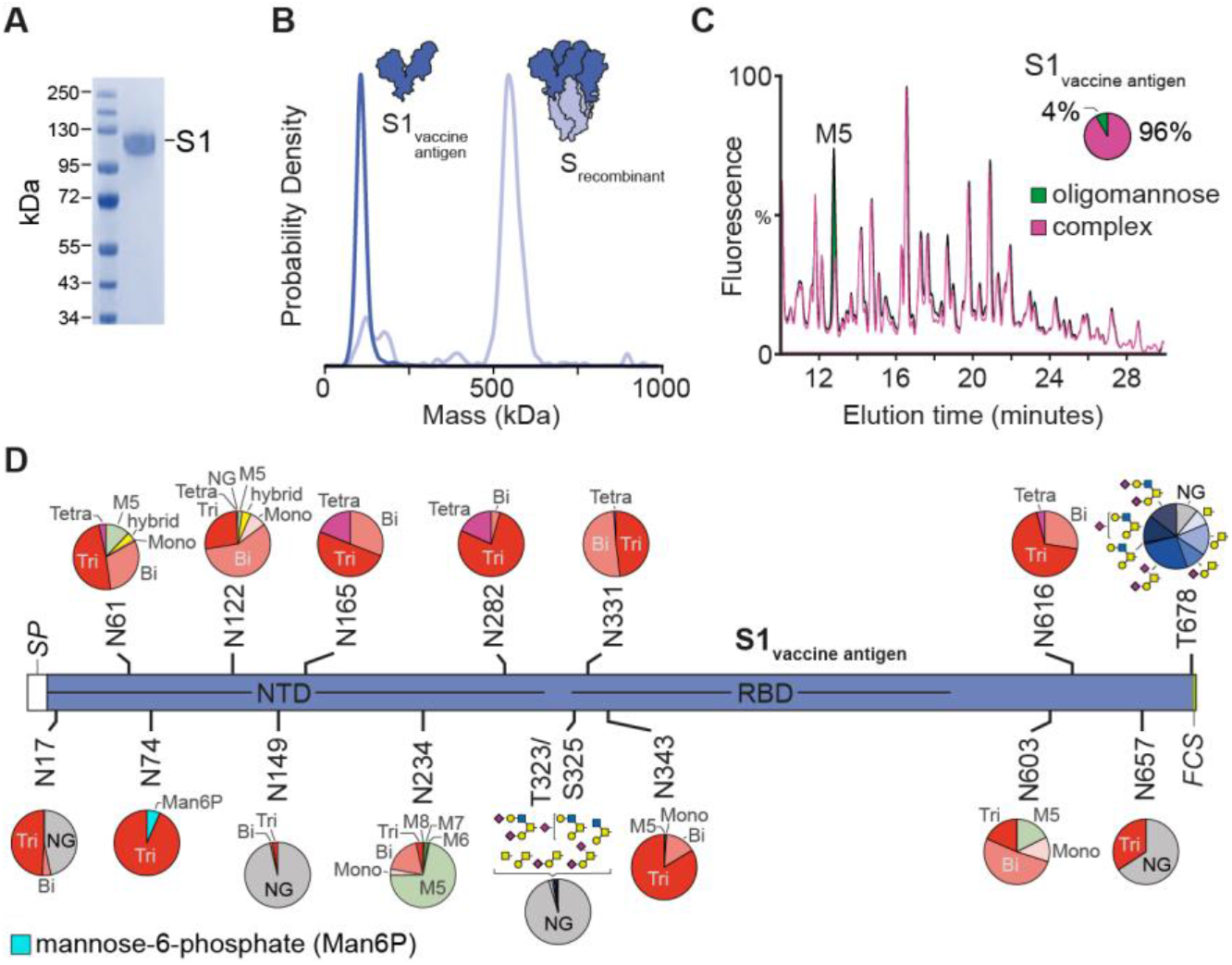
Glycosylation and assembly of vaccine-derived spike protein. (**A**) SDS-PAGE of CR3022 purified S1_vaccine antigen_; (**B**) Mass photometry of monomeric S1_vaccine antigen_ (~ 120kDa) and S_recombinant trimer_ (~ 550 kDa); (**C**) Quantitative UPLC N-glycan analysis of S1_vaccine antigen_ showing the degree of glycan processing; (**D**) Site-specific N- and O-glycosylation of S1_vaccine antigen_ (see Figure 1E for pie chart legend).

With the aim of comparing site-specific S glycosylation in the context of vaccine design and antigen structure, we produced S in mammalian cells using an expression construct identical to the one used in creating ChAdOx1 nCoV-19 [16]. This contains SARS-CoV-2 amino acids 2-1273 preceded by an N-terminal leader peptide consisting of tissue plasminogen activator (tPA) and a modified human cytomegalovirus major immediate early promoter. Using the same purification strategy as above, we observed that the majority of over-expressed protein was secreted into the supernatant as soluble S1 (herein referred to as S1_vaccine antigen_), as detected by SDS-PAGE (**Figure 2A**) and confirmed by mass spectrometry (**Supplementary Figure 1B**). S2 remained cell associated, embedded in the lipid bilayer, as shown by western blot probed with an antibody against S2 (**Supplementary Figure 3**). We analysed the secreted S1_vaccine antigen_ by mass photometry, and compared it to the stabilised S_recombinant trimer_, which revealed the shed vaccine antigen to be solely monomeric (**Figure 2B, Supplementary Movies 1 and 2**).

**Figure 3:**
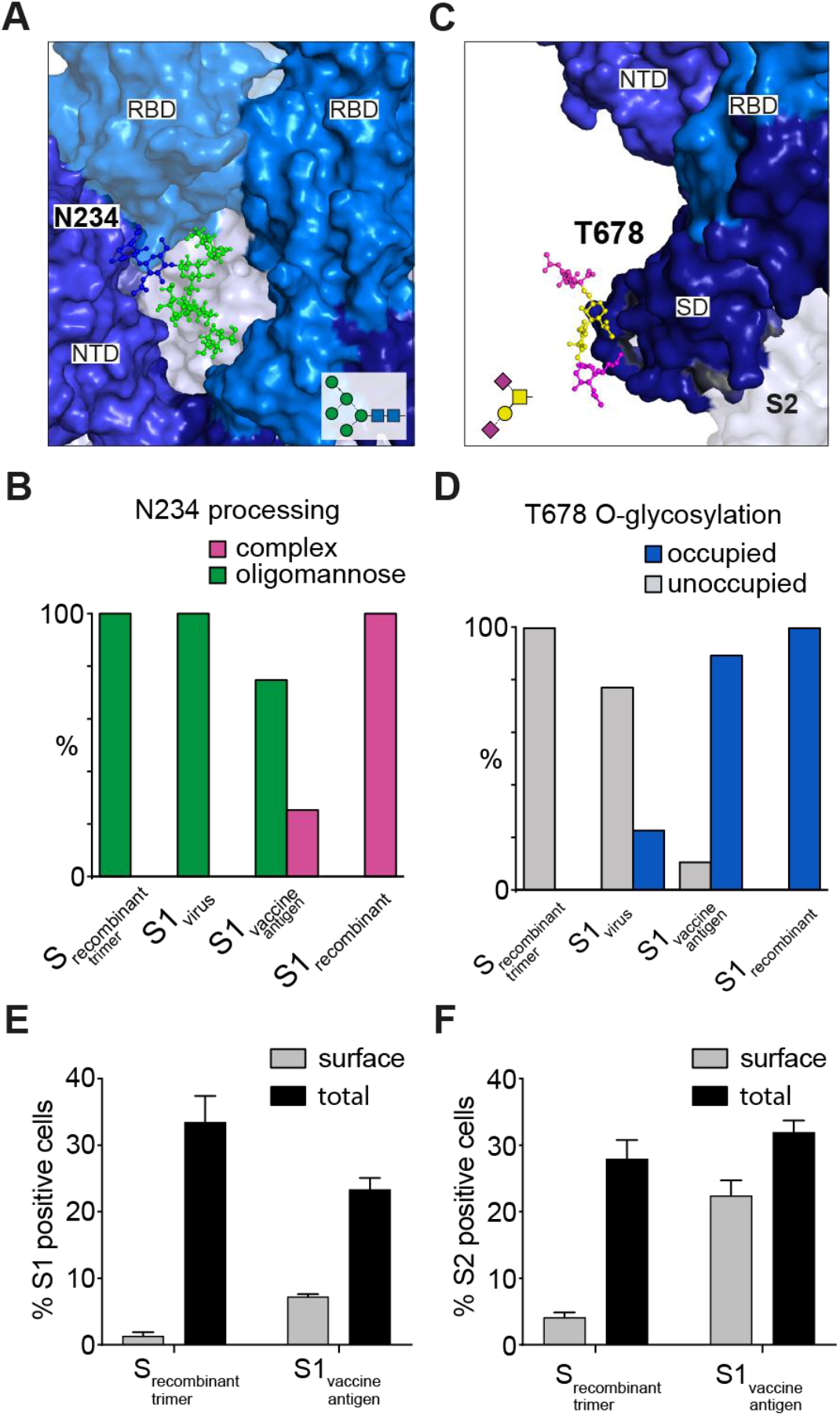
Correlation of spike cellular location and macromolecular assembly with N234 and T678 glycan processing. (**A**) Structural position and orientation of the S1 N-glycan N234 (shown as Man_5_GlcNAc_2_) in a pocket formed by the receptor binding domain (RBD, top right corner) and N-terminal domain (NTD) of the same protomer, and the neighbouring RBD (top left corner). GLYCAM web server (http://glycam.org) was used to model the glycan on to the PDB 6VXX; (**B**) Percentage change in oligomannose content of the N234 N-glycan of S_recombinant trimer_, S1_virus_, S1_vaccine candidate_ and S1_recombinant_; (**C**) Location of the S1 O-glycan T678 (shown as di-sialylated core-1 structure) located in the subdomain (SD) near the furin cleavage site between S1 and S2 (modelled on PDB 6VXX using GLYCAM webserver); (**D**) Changes in T678 O-glycan occupancy across samples tested; (**E**) Percentage of HEK293F S_recombinant trimer_- or S_vaccine antigen_-transfected cells stained positive for S1 or (F) S2 solely on the cell surface or in permeabilised cells. Data are shown as mean ± SEM (n = 3).

Glycan content analysis of S1_vaccine antigen_ demonstrated an extraordinary 96% of complex N-glycans and only 4% of oligomannose-type N-glycans (**Figure 2C**), indicating an increase in accessibility of glycan processing enzymes in the Golgi to S1_vaccine antigen_ glycan sites compared to S1_virus_. Quantitative site-specific N- and O-glycosylation analysis of S1_vaccine antigen_ confirmed that although overall N-glycan site occupancy was comparable to S1_virus_, except for N17, which was 47% non-glycosylated; the large majority of N-glycans attached to S1_vaccine antigen_ underwent considerably more processing, likely after furin cleavage in the Golgi, as evidenced by the presence of increased complex glycosylation. The N61 and N603 sites, which were 98% and 83% oligomannose on S1_virus_, became 12% and 18% on S1_vaccine antigen_, respectively. We also detected an increase in T323/S325 and T678 O-glycan extensions (i.e. presence of core-2 structures) as well as a 50% increase in sialylation at T678. Finally, N-glycan sites that had mixed oligomannose and complex glycan populations on S1_virus_ (N74, N122, N343 and N616) become heavily processed on S1_vaccine antigen_ (**Figure 1F** and **Figure 2D** for S1_virus_ and S1_vaccine antigen_, respectively).

However, a single N-glycan site is maintained in an underprocessed state. For S1_virus_, 60% of the N-glycans at position N234 were Man_6-8_GlcNAc_2_ (M6, M7 and M8) structures. In contrast, although the S1_vaccine antigen_ carried the slightly more processed M5 N-glycan, which is not accessible to GlcNAc-transferase I in the cis-Golgi, the remaining structures did not progress to more complex type glycosylation like the rest. This prevention of more extensive glycan processing of the N-glycan at N234 is due to the spatial and temporal assembly of S proteins in the ER and Golgi. On a fully assembled S trimer, N234 glycans are located in a pocket formed partly by the receptor binding domain (RBD) and the N-terminal domain (NTD) on the same protomer, and partly by a neighbouring RBD, which gives rise to the largely underprocessed oligomannose structures when early N-glycan trimming at this site is prevented by trimerisation of S in the ER (**Figure 3A**). In both recombinant trimer-derived and viral S1, N234 was 100% oligomannose, but dropped to 74.8% oligomannose on vaccine-derived S1 (**Figure 3B**). The fact that this site was also underprocessed on S1_vaccine antigen_ indicates that this protein is derived from a spike that initially trimerised in the ER, but that this trimer is apparently less “closed” and more accessible to mannose-trimming ER enzymes compared to its counterpart expressed by the SARS-CoV-2 virus.

This slightly less closed vaccine trimer travels in a membrane-bound form from the ER to the ERGIC, but as there are no viruses present to incorporate these trimers into their envelopes when budding into the ERGIC lumen, the overexpression system causes the S_vaccine antigen_ trimers to be pushed further along the membranes of the cis-, medial- and trans-Golgi. We detected mannose-6-phosphate (M-6-P) on S1_vaccine antigen_ (**Figure 2D, Supplementary Figure 4**), with some evidence for this modification also on S1_virus_ (**Supplementary Figure 5**). This sugar tag is initially added in the cis-Golgi in the form of GlcNAc-M-6-P, then decapped in the trans-Golgi and recognised by the M-6-P receptor responsible for directing tagged proteins and possibly whole viruses from the trans-Golgi to late endosomes/lysosomes; such lysosomal egress has recently been described for SARS-CoV-2 virus [9]. With the furin cleavage site intact, S_vaccine antigen_ is cleaved by furin in the trans-Golgi. However, unlike endogenous viral spikes, where we postulate that additional stabilising viral factors are present, S1_vaccine antigen_ dissociates from S2_vaccine antigen_ upon furin cleavage and becomes secreted. This shedding occurs in the trans-Golgi rather than at the plasma membrane of the cell, as evidenced by the increased N-glycan processing by late-stage Golgi glycosylation enzymes, resulting in the high complex-type N-glycan content of S1_vaccine antigen_, and also by the substantially increased O-glycosylation occupancy levels on T678 (**Figure 3D, Supplementary Figure 6**). Plausibly, the modest amount of S1_virus_ T678 O-glycosylation is related to furin cleavage, making S1_virus_ more accessible to O-GalNAc-transferases; however, differences in virus assembly and the continuous association with, and shielding by, S2_virus_ prevents this from reaching similar O-glycan occupancy levels as that of cleaved soluble S1_vaccine antigen_ (**Figure 3D**). Although S_recombinant trimer_ transits the trans-Golgi in soluble form, it is not O-glycosylated at this position as it lacks the furin site (R682-R685, **Figure 3C**), cleavage of which appears to favour this processing step.

**Figure 4:**
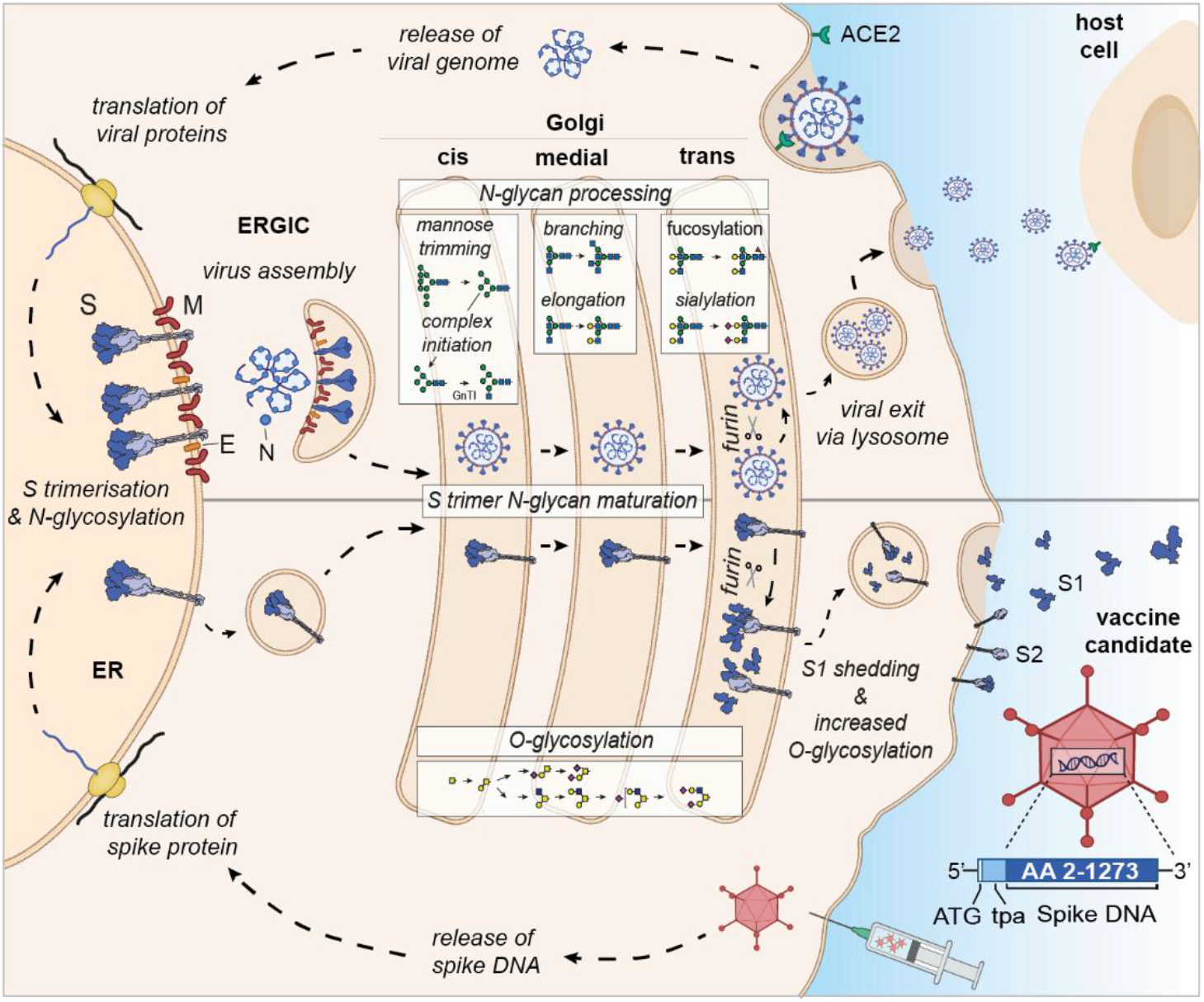
Differential expression and glycan processing of virions and vaccine derived spike glycoproteins. SARS-CoV-2 binds to its receptor ACE-2 and infects cells, leading to the release of the viral genome and translation of viral proteins. Spike protein is co-translationally N-glycosylated and forms trimers in the ER that traffic to the ERGIC where they are incorporated into budding virions. Individual virions continue through the secretory pathway to the trans-Golgi prior to following a lysosomal egress route. For the vaccine candidate, spike DNA is administered via an adenovirus vector system, and spike protein is synthesized in the ER, where it is N-glycosylated and trimerises as before, but as it is not incorporated into a budding virion in the ERGIC, it continues through the secretory pathway and, via lysosomes, to the plasma membrane. In both cases the spike glycoproteins have access to both the N- and O- linked host glycosylation machinery. Upon furin cleavage in the trans-Golgi, S1 and S2 of the virus stay non-covalently associated, whereas furin cleavage of the vaccine antigen results in shedding of monomeric S1_vaccine antigen_. Glycomic signature analysis of these two proteins show that the N-linked glycosylation occupancy levels, which are determined in the ER, are comparable for S1_virus_ and S1_vaccine antigen_ whereas the attached glycoforms vary reflecting their different accessibility to glycan processing enzymes. S1_vaccine antigen_ carries not only higher levels of complex N-glycans but is also extensively O-glycosylated after furin cleavage in the trans-Golgi, when most S1_vaccine antigen_ is shed and secreted in a soluble monomeric form. Some S1 and S2_vaccine antigen_ is displayed on the cell surface, presumably as trimers.

To test our hypothesis that S1_vaccine antigen_ comes from an assembled S trimer and is shed in the trans-Golgi following furin cleavage, we expressed the individual S1_recombinant_ subunit (**Supplementary Figures 7**), that cannot trimerise, and quantified the extent of N-glycan processing at N234 and O-glycosylation at T678. S1_recombinant_ had 100% complex-type glycans at N234, shifting from 25.3% complex for S1_vaccine antigen_ and 0% for S_recombinant trimer_ and S1_virus_ (**Figure 3B**). Similarly, T678 O-glycan occupancy was reversed from 0% (S_recombinant trimer_), 22.7% (S1_virus_), 89.5% (S1_vaccine antigen_) to 100% (S1_recombinant_) (**Figure 3D**). The site-specific changes across all S1 samples are illustrated in **Supplementary Figure 8**). However, cleavage of S_vaccine antigen_ by furin is not complete; around 10% was not O-glycosylated and appeared on the cell surface, which would not happen during natural virus infection. This was shown by fluorescence activated cell sorting (FACS) analysis using staining with the RDB-specific CR3022 antibody of either unpermeabilised or detergent-permeabilised cells (**Figures 3E & 3F**). It may be this potentially trimerised and still S1-containing cell surface accessible spike that gives rise to the promising antibody responses reported in an early phase ChAdOx1 nCoV-19 clinical trial [16], [17].

These results are encouraging, showing that it may be possible to improve on immunogen design. Shedding of monomeric and non-physiologically glycosylated S1_vaccine antigen_ from immunogen producing cells is reminiscent of HIV vaccine development, where early immunogens were hampered by the inability of monomeric gp120 to elicit a broadly neutralising antibody response, which is needed for virus neutralisation [26]. Indeed, immunogens that do not mimic trimeric spike glycoproteins as they are presented on infectious virions may effectively act as a decoy, eliciting more of the unwanted sub-optimal or non-neutralising antibodies that are incapable of binding to and neutralising trimeric spikes on the virus [15], [26]–[29]. For SARS-CoV-2, most neutralising antibodies bind to the trimer apex (**Supplementary Table 2**), and those will not be elicited by shed S1_vaccine antigen_ which lacks both the correct protein architecture and exposes non-neutralising epitopes that would otherwise be buried on assembled spikes. For example, S2M11 [30] (**Supplementary Figure 9A**) and C144 [31] bind on a quaternary epitope formed by two neighbouring RBDs on the trimer apex. The binding of neutralising antibodies that incorporate N-glycans as part of their binding epitopes, e.g. S309 [32] (**Supplementary Figure 9B**) and BD-23 [33] , will also be adversely affected by a vaccine antigen that differs from circulating viruses in a natural infection. Furthermore, specific glycans, including N234, affect the up/down orientation of the RBD domain [12] pointing to the critical need for physiological glycosylation on effective vaccines based on S1. Yet other neutralising antibodies could bind to peptide epitopes of S1 that may still be available on shed S1_vaccine antigen_ but that may be less accessible because of the non-physiologically high amount of complex N-glycans shielding those epitopes.

A strong B-cell response is based on the immunogen mimicking parts of an invading pathogen. Therefore, for SARS-CoV-2 we suggest that a stabilised trimeric pre-fusion spike protein, with the furin cleavage site abolished, and with non-physiological areas shielded to prevent unwanted non-neutralising immune response, may be able to elicit neutralising antibodies with the desirable significant breadth and potency. Viral vector-based, such as ChAdOx1 nCoV-19, as well as nucleic acid-based vaccine strategies, such as the Pfizer BNT162b2 and Moderna mRNA-1273 vaccines, rely on the supplied antigen-encoding DNA or RNA sequence, once inside a cell, to faithfully produce the spike protein in its fully folded, glycosylated and assembled state, resembling a natural infection and trigger a robust innate immune response, as well as provoking T and B cells. However, the cellular secretion pathway followed by such vaccine delivered antigens may differ in fundamental ways from antigens in the context of viral infection, where factors other than a single protein coding sequence may play decisive roles in immunogen presentation (**Figure 4**). These include the (intra)cellular location of viral morphogenesis (i.e. from which organelle a virus buds), as well as the overall shape in which an immunogen encounters the host cellular glycosylation machinery during a natural infection. The Pfizer BNT162b2 vaccine antigen aims to overcome some of these important differences by following a strategy first employed for MERS, as well as SARS-CoV spike glycoprotein stabilisation for vaccine design [34], [35], where two proline mutations are introduced in close proximity to the first heptad repeat of each protomer, which stabilises the prefusion conformation [36].

Abolishing the furin cleavage site and introducing mutations that lock the spike protein in the prefusion conformation and prevent shedding of S1 are likely to elicit more potent antibody responses. Some vaccine candidates already combine both approaches [37], [38] and it would be interesting to compare their glycan signatures to that of wild-type virus. Characterising and understanding the correct glycosylation of the virus will be crucial in the development of a high-quality immune response, and glycan processing may inform vaccine design strategies, aimed at achieving the correct immunogen presentation, for this and future pandemics.

## Acknowledgements

We thank the Krammer Laboratory for kindly providing the spike (soluble domain) and CR3022 expression vectors, Corinne Lutomski and Tarick El-Baba (University of Oxford, Department of Chemistry) for the Krogan Laboratory full-length spike expression vector (sourced from Addgene), Anderson Ryan (University of Oxford, Department of Oncology) for Calu-3 cells and Simon Draper (University of Oxford, Nuffield Department of Medicine) for the pENTR4-LPTOS vector. We thank Protein Metrics for software support. Figures 1 and 4 were created the aid of BioRender.com. We thank Sarah Karin Wideman, Felix Clemens Richter and Johannes Pettmann for technical assistance for Flow Cytometry. W.B.S. acknowledges the support of the philanthropic donors to the University of Oxford’s COVID-19 Research Response Fund and BBSRC/UKRI BB/V011456/1 grant (Molecular Mapping of SARS-CoV-2 and the Host Response with Multiomics Mass Spectrometry to Stratify Disease Outcomes). JB is supported by a Wellcome Trust PhD Programme (203853/Z/16/Z). JLK is supported by a Lerner-Fink Fellowship in Medicinal Chemistry. This work was supported by the Oxford Glycobiology Endowment.

