## Supplementary Material for "Analysis of SARS-CoV-2 spike glycosylation reveals shedding of a vaccine candidate"

### Material and Methods

*Cell culture*

African green monkey (*Chlorocebus sabaeus*) Vero E6 cells (ATCC CRL-1586) were grown in Dulbecco's Modified Eagle Medium (DMEM, Gibco) containing 4.5 g/L glucose and 2 mM L-glutamine supplemented with 10% heat inactivated fetal bovine serum (HI FBS, Gibco), 100 U/mL penicillin and 0.1 mg/mL streptomycin (Sigma). The human lung cancer cell line Calu-3 (kindly provided by Dr Anderson Ryan, Department of Oncology, University of Oxford, as part of an ongoing collaboration) was cultured in a 1:1 mixture of DMEM and Ham's F-12 medium (Gibco) supplemented with 10% HI FBS, 1% non-essential amino acids (Gibco), 1% sodium pyruvate (Gibco), 100 U/mL penicillin and 0.1 mg/mL streptomycin (Sigma). Incubations were carried out at 37 °C and 5% $CO_2$. The FreeStyle™ 293-F (HEK 293F, ThermoFisher Scientific) cell line was cultured in Freestyle 293 expression media (ThermoFisher Scientific) and incubated at 37 °C, 5% $CO_2$ and 130 rpm.

*Virus propagation*

The SARS-CoV-2 England/02/2020 strain (GISAID: EPI_ISL_407073) was provided at passage one from Public Health England, Collindale. Viral stocks were obtained by infecting Vero E6 cells at a multiplicity of infection (MOI) of 0.01 in virus propagation medium (DMEM with 2% HI FBS) and harvested 72 h post infection (hpi). Culture media was centrifuged at 3200 x g for 5 min, aliquoted and stored at –80 ˚C.
To obtain SARS-CoV-2 virions for endogenous spike glycoprotein purification, Calu-3 cells were infected with SARS-CoV-2 in Calu-3 media containing 2% HI FBS at a MOI of 0.1. Cell culture supernatant was harvested 72 hpi and centrifuged at 3200 x *g* for 5 min. Virus-containing supernatant was concentrated one-log - using a 100 kDa cut-off centrifugal filter (Amicon, Merck) - with subsequent inactivation of virions with a final concentration of 0.5% Triton X-100 (Sigma) by incubation for 30 min at 4 ˚C.

*Expression and purification of the CR3022 antibody*

The recombinant monoclonal antibody CR3022 was transiently expressed in HEK293F cells. The cells were co-transfected with plasmids encoding the heavy- and light chain of CR3022 (a kind gift from the Krammer Laboratory, Department of Microbiology, Icahn School of Medicine at Mount Sinai, New York) cloned into pFUSEss and pFUSEss2, respectively (1:2 molar transfection ratio) utilising FreeStyle™ MAX reagent (Invitrogen) and OptiPRO™ SFM (Gibco) following the manufacturer's protocol. Seven days post transfection, cell culture supernatant was harvested by centrifugation at 3000 x *g* for 45 min. Sodium phosphate was added to a final concentration of 20 mM, the pH adjusted to 7.2 and the sample sterile filtered. The solution was applied to a Protein A-Sepharose Fast Flow column (GE Healthcare) equilibrated in 20 mM sodium phosphate, pH 7.2. Antibody elution took place using 100 mM citric acid (pH 3.5) and the eluate pH adjusted to 7.2 using 1M Tris-NaOH. The sample was further purified by size exclusion chromatography using a Superdex™ 200 pg 16/600 column (GE Healthcare) equilibrated in 10 mM phosphate buffer, 2.7 mM potassium chloride 137 mM sodium chloride, pH 7.4 (PBS).

*SARS-CoV-2 S expression constructs*

*Vaccine candidate ($S_{vaccine\ antigen}$)*

The sequence encoding SARS-CoV-2 (Addgene # 141382) was cloned from amino acids 2-1273 into a pENTR4-LPTOS vector (kindly provided by Dr Simon Draper, Jenner Institute, University of Oxford) in frame to an N-terminal secretion leader peptide tissue plasminogen activator (tPA), using InFusion cloning (Clontech). The plasmid also encodes a modified human cytomegalovirus major immediate early enhancer/promoter (IE CMV).

*Recombinant, soluble spike glycoprotein ($S_{recombinant\ trimer}$)*

The vector pCAGGS encoding for the SARS-CoV-2 spike glycoprotein was a kind gift from the Krammer Laboratory, Department of Microbiology, Icahn School of Medicine at Mount Sinai, New York. In short, the SARS-CoV-2 spike glycoprotein (GenBank isolate: MN908947) residues 1-1213 were codon-optimised for mammalian cell expression; the polybasic cleavage site (RRAR, residues 682-685) were mutated to alanine and the prefusion conformation stabilising mutations K986P and V987P (numbering according to wild type) were introduced. In addition, the construct has a C-terminal thrombin cleavage site, T4 fibritin trimerisation domain and a hexahistidine tag.

*Recombinant S1 ($S1_{recombinant}$)*

The nucleotides encoding for residues 1-682 of the SARS-CoV-2 spike glycoprotein (GenBank isolate: MN908947) were optimised for mammalian cell expression and subcloned into the mammalian expression vector pHLsec (AgeI/-KpnI linearised) using Gibson assembly reaction mix (NEB) following the manufacturer’s protocol. All recombinant construct products were used to heat-shock transform *E. coli* DH5$\alpha$ competent cells (Clontech). Cells were incubated at 37 °C for 16 h under antibiotic selection (25 µg/mL kanamycin or carbenicillin for $S_{vaccine\ antigen}$, $S_{recombinant\ trimer}$ or $S1_{recombinant\ monomer}$, respectively) and single colonies were subsequently picked for DNA sequencing (Source Biosciences).

*Recombinant SARS-CoV-2 S glycoprotein expression*

Proteins were transiently expressed in HEK293F cells. The cells were transfected utilising FreeStyle™ MAX reagent (Invitrogen) and OptiPRO™ SFM (Gibco) following the manufacturer’s protocol. Five days post transfection, cell culture supernatant was harvested by centrifugation at 3000 x g for 45 min. The sample was applied to an immunoaffinity chromatography column with immobilised CR3022 antibody as an antagonist (see below).

*Affinity purification of $S_{virus}$, $S_{recombinant\ trimer}$, $S1_{recombinant\ monomer}$ and $S_{vaccine\ antigen}$*

CR3022 antibodies were immobilised on cyanogen bromide activated Sepharose 4 resin (GE Healthcare) according to the manufacturer’s protocol (Instructions 71-7086-00 AF) for subsequent use in immunoaffinity purification. The culture supernatant of SARS-CoV-2 infected Calu-3 cells (in total: $56 \times 10^6$ plaque forming units) was collected and the virus inactivated as previously described. Triton-X100 (0.5% w/v), carried over from the inactivation step, was diluted below its CMC (~ 0.02% w/v) using 20 mM Tris-HCl, 500 mM NaCl and a final concentration of 0.03% (w/v) *n*-dodecyl-b-D-maltoside (DDM) added. Subsequently, the solution was loaded onto a CR3022-

affinity column, washed with 10 column volumes (CV) of high-salt wash buffer (20 mM Tris-HCl, 500 mM NaCl, 0.03% DDM) and eluted with 3 M MgCl2, 0.03% DDM, pH 7.2 (2CV). The eluted protein was immediately buffer exchanged into 10 mM Tris-HCl,75 mM NaCl, 0.03% DDM, pH 7.2 and further concentrated using a 30 kDa cut-off centrifugal filter (GE Healthcare). Recombinant spike glycoproteins ($S_{vaccine\ antigen}$, $S_{recombinant\ trimer}$ and $S1_{recombinant}$) were purified in the same manner by omitting the detergents (Triton-X100 and DDM) throughout the purification process.

*Flow cytometry*

HEK293F cells transfected with constructs for $S_{recombinant\ trimer}$ or $S_{vaccine\ antigen}$ were harvested 48 h post transfection by centrifugation at 400 x *g* for 5 min, and stained using LIVE/DEAD fixable near-IR dead cell stain kit (Life Technologies) prior to fixation in 2% paraformaldehyde (Electron Microscopy Sciences) for 20 min to exclude dead cells. Cells were blocked in fluorescence activated cell sorting (FACS) block (PBS with 0.5% w/v BSA, 5 mM EDTA, 5% HI goat serum and 5% HI normal human plasma) for 20 min, followed by permeabilisation with 0.5% saponin (Sigma) in FACS block for 20 min on ice. Cells were first surface stained with an antibody recognising S1 (CR3022; 20 µg/mL) or with an antibody recognising S2 (Genetex #GTX632604; 10 µg/mL) for 40 min on ice. Primary antibodies were detected using 10 µg/mL of the appropriate Alexafluor 488-conjugated antibody (Invitrogen #A-11013 or #A-11008, respectively). Flow cytometry acquisition was performed using an Attune NxT Flow Cytometer (Thermofischer) and gated for live cells. A minimum of 10,000 gated events were identified and results were analysed using the software tool FlowJo V10.

*SDS-PAGE and western blot*

Recombinant and endogenous proteins were analysed using 4-12% NuPAGE Bis-Tris SDS-PAGE gel (Invitrogen) to determine the protein expression in either the supernatant or the cell pellet. To determine the expression of $S_{vaccine\ antigen}$ in HEK293F cells, cells were harvested 4 days post transfection as described above. The resulting cell pellet was resuspended in lysis buffer (1% DDM in PBS); the suspension was sonicated on ice three times for 5 min using pulses of 0.7s on and 0.3s off with an applied 65% intensity. After protein separation using SDS-PAGE, gels were either stained with SimplyBlue SafeStain (Invitrogen) for 1 h and de-stained in distilled water overnight, or used for western blot analysis (iBlot™, Thermo Fischer Scientific). Blots were incubated with the primary antibodies listed in the Flow Cytometry section and additionally against S2 (Genetex #GTX635693), and respective HRP secondary antibodies (GE Healthcare #NA933, Promega #W4011, Sigma #71045).

*In-gel protease digestion*

The SDS-PAGE bands corresponding to spike glycoprotein were excised from the gel and cut into pieces. The gel pieces were washed twice with water, twice with 50%v/v acetonitrile followed by 100%v/v acetonitrile. The spike glycoprotein sequence was separately digested *in silico* with three sequencing grade proteases (trypsin, chymotrypsin and alpha lytic protease) to confirm peptides of suitable length (between 7 to 25 amino acids) covering the 22 potential N-glycosylation sites. Gel pieces were rehydrated with 100 mM ammonium bicarbonate, 100 mM Tris-HCl 10 mM calcium chloride pH 8.0 or 100 mM ammonium bicarbonate for digests with trypsin, chymotrypsin and alpha lytic protease, respectively. Gels were dehydrated with 100%v/v acetonitrile and dried down in a centrifugal vacuum concentrator before rehydration for 45 min at 56°C with 10 mM dithiothreitol in 100 mM ammonium bicarbonate, 100 mM Tris-HCl 10 mM calcium chloride pH 8.0 or 100 mM ammonium bicarbonate for digests with trypsin, chymotrypsin and alpha lytic protease, respectively. Proteins were alkylated with 55 mM iodoacetamide in the respective buffer for 30 min in the dark. The gel pieces were washed twice with the respective buffer followed by the buffers in 50% v/v acetonitrile before drying in a centrifugal vacuum concentrator. Gel pieces were rehydrated on ice for 45 min using 12.5 ng/µl of trypsin, chymotrypsin (Promega) and alpha lytic protease (Sigma) in 100 mM ammonium bicarbonate, 100 mM Tris-HCl 10 mM calcium chloride pH 8.0 or 100 mM ammonium bicarbonate, respectively. Proteins were digested at 37 °C for 16 h. Gels were washed in the respective buffer in 50% v/v acetonitrile before extracting the peptides twice with 5%v/v formic acid and acetonitrile. Peptide extracts were dried, resuspended in 0.05%v/v TFA using a water bath sonicator and then run by LC-MS.

*In-solution digestion*

Endogenous spike (8 µg) purified by CR3022 was dried down and denatured with 8M urea containing 200 mM ammonium bicarbonate. Proteins were reduced with tris(2-carboxyethyl)phosphine (1 mM final concentration) in 100 mM ammonium bicarbonate for 1 h at 56 °C followed by alkylation with iodoacetamide (4.4 mM final concentration) in 100 mM ammonium bicarbonate for 1 h in the dark. Urea was diluted to 1 M with 100 mM ammonium bicarbonate and then the proteins were digested with 1:100 w:w alpha lytic protease:protein for 16 h at 37 °C. An additional 1:100 w:w alpha lytic protease:protein was added and the digestion was continued for 5 h. The peptide containing sample was dried, resuspended in 0.05%v/v TFA using a water bath sonicator and then run by LC-MS with online desalting.

*Liquid chromatography*

Peptides were separated on a Dionex Ultimate 3000 nano UHPLC system (Thermo Scientific). A nano analytical C18 reversed phase column (PepMap) was used with dimensions 75 µm x 15 cm, 2 µm particle size (Thermo Scientific) with a flow rate of 300 nL/min at 35 °C. The mobile phase used was solvent A: 0.1% v/v formic acid in LC-MS grade water, and solvent B: 0.1% v/v formic acid in 80% v/v acetonitrile. The gradient used to separate peptides on the analytical column was: 2% B (0-4.5 min), 2-30% B (4.5-40 min), 30-95% B (40-55 min), 95% B (55-59 min), 95-2% B (59-60 min) and 2% B (60-70 min) for column equilibration.

*Mass spectrometry*

Peptides from the nano LC were analysed on a benchtop Q Exactive hybrid quadrupole-Orbitrap mass spectrometer using the Nanospray Flex ion source. Prior to data acquisition, the mass spectrometer was calibrated for mass accuracy according to the manufacturer's recommendations using a positive ion calibration solution injected at 5 µl/min into a heated electrospray ionisation (HESI) probe (Thermo Scientific). The conditions for data dependent acquisition (DDA) were as follows: chromatographic peak width was set at 12 s and the Full MS conditions were with a resolution of 70,000, AGC target of 3e6, maximum IT (injection time) of 60 ms, scan range of 375 to 1500 m/z. The dd-MS2 conditions were with a resolution of

17,500, AGC target of 1e5, maximum IT of 60 ms, loop count of 10 (i.e. Top 10), isolation window of 2.0 m/z, fixed first mass of 100.0 m/z and normalised collision energy (NCE) was stepped at 27, 30 and 33 in a high-energy collision dissociation (HCD) cell. The data-dependent (dd) settings were with minimum intensity threshold of 3.3e4 ions, charge exclusion: unassigned, 1, >8, peptide match: preferred, dynamic exclusion: 20 s.

*Protein identification and sequence coverage*

The acquired *.raw files were converted to Mascot generic files (*.mgf) using MSConvertGUI 64-bit (ProteoWizard version: 3.0.19172-57d620127). A peak-picking filter was set with MS levels 1-2. The Mascot server software (Matrix Science, London) was used to search the *.mgf files against the SwissProt database (containing SPIKE_SARS2 spike glycoprotein P0DTC2 in Version 2020_03 with 562,755 sequences released on 17$^{th}$ June 2020). The search parameters in Mascot were set as follows: MS/MS Ion Search with either trypsin, chymotrypsin or 'none' for alpha lytic protease (with up to 3 missed cleavages). Carbamidomethyl (C) was set as a fixed modification and Oxidation (M) as a variable modification. Taxonomy was set to 'All'. Peptide mass tolerance and fragment mass tolerance were both set as ± 10 ppm and the peptide charge was set as +2, +3 and +4. Data format and instrument were set as Mascot generic and Q Exactive, respectively. A decoy database was searched in each case with the false discovery rate (FDR) adjusted to 1%. Sequence coverage was determined using only peptides with the statistical strongest assignment in bold red (i.e. highest-scoring match for a given MS/MS spectra and highest scoring protein in which the peptide sequence appears).

Glycopeptide data was analysed via Byonic$^{TM}$ v3.10.10 and Byologic$^{TM}$ v3.10-52x64 (Protein Metrics Inc.). Byonic search settings were as follows: Precursor mass tolerance = 5, Fragment mass tolerance = 20, FDR = 1% with a glycan library consisting of 132 N-glycan and 9 O-glycan structures. Additional permissible modifications were oxidation (Met/Trp), deamidation (Asn/Gln), N-terminal Gln to PyroGln and Cys carbamidomethyl (fixed modification). Glycopeptides search criteria was set for a minimum score ≥ 200 and <0.001 Pep2D value and data was inspected manually in Byonic to ensure quality of MS/MS spectra and precursor monoisotopic peak assignments. Byologic was used to obtain the area of each extracted ion chromatogram, which were exported and used to quantify site-specific glycosylation at each site.

*Mass photometry*

Mass photometry measurements were done as described in detail [1]. Briefly, borosilicate microscope coverslips were cleaned via sonication in Milli-Q $H_2O$, isopropanol, Milli-Q $H_2O$ and dried under a nitrogen gas flow. Sample chambers were assembled using silicone gaskets (CultureWell™ reusable gasket, 3mm diameter x 1 mm depth, Grace Bio-Labs). Coverslips were placed on the sample stage and a single gasket was filled with 10 µl phosphate buffered saline (PBS) to find focus and ensure a low signal-to-noise background. Immediately prior to each sample measurement, $S1_{vaccine\ candidate}$ and $S_{recombinant\ trimer}$ were diluted from a 1 µM stock (object concentration) to 100 nM, of which 2 µl was added to the 10 µl PBS in a single unused gasket, and data were recorded for 90 seconds. MP settings were as follows: frame averaging = 50, frame rate = 955 Hz and 4x4 pixel binning. Data was analysed using Discovery MP v2.3dev7 (Refeyn Ltd).

*Fluorescent labelling of N-linked glycans*

Glycans of the peptides from the in-gel trypsin digests were enzymatically released by PNGase F (NEB) for 16 h at 37 °C. Released glycans were labelled with 2-aminoanthranilic acid (2-AA) as previously described [2]. Briefly, glycans were resuspended in 30 μl of water followed by addition of 80 μl of labelling mixture (30 mg/ml 2-AA and 45 mg/ml sodium cyanoborohydride in a solution of sodium acetate trihydrate [4% w/v] and boric acid [2% w/v] in methanol). Samples were then incubated at 80 °C for 1 h. Excess label was removed using Spe-ed Amide-2 cartridges (Applied Separation), as previously described [2].

*Hydrophilic interaction liquid chromatography-ultra performance liquid chromatography*

Fluorescently labelled glycans were resolved by hydrophilic interaction liquid chromatography-ultra performance liquid chromatography (HILIC-UPLC) using a 2.1 mm × 10 mm Acquity BEH Amide Column (1.7 m particle size) (Waters, Elstree, UK). The mobile phase used was solvent A: 50 mM ammonium formate, pH 4.4, and solvent B: acetonitrile. The gradient used to separate fluorescently labelled glycans on the analytical column was: time = 0 min ($t$ = 0): 22.0% A, 78.0% B (flow rate of 0.5 ml/min); $t$ = 38.5: 44.1% A, 55.9% B (0.5 ml/min); $t$ = 39.5: 100% A, 0% B (0.25 ml/min); $t$ = 44.5: 100% A, 0% B (0.25 ml/min); $t$ = 46.5: 22.0% A, 78.0% B (0.5 ml/min), $t$ = 48: 22.0% A, 78.0% B (0.5 ml/min). Fluorescence was measured using an excitation wavelength of 360 nm and a detection wavelength of 425 nm.

*Sequencing of N-linked glycans*

Endoglycosidase H (Endo H, NEB) was used for quantitation of oligomannose structures. Digestions were performed at 37 °C for 16 hours, according to manufacturer's instructions. The digested glycans were purified using a PVDF protein-binding membrane plate (Millipore) prior to HILIC-UPLC analysis. Data processing was performed using Empower 3 software. The percentage abundance of oligomannose-type glycans was calculated by integration of the relevant peak areas before and after Endo H digestion, following normalisation.

### Supplementary Figures

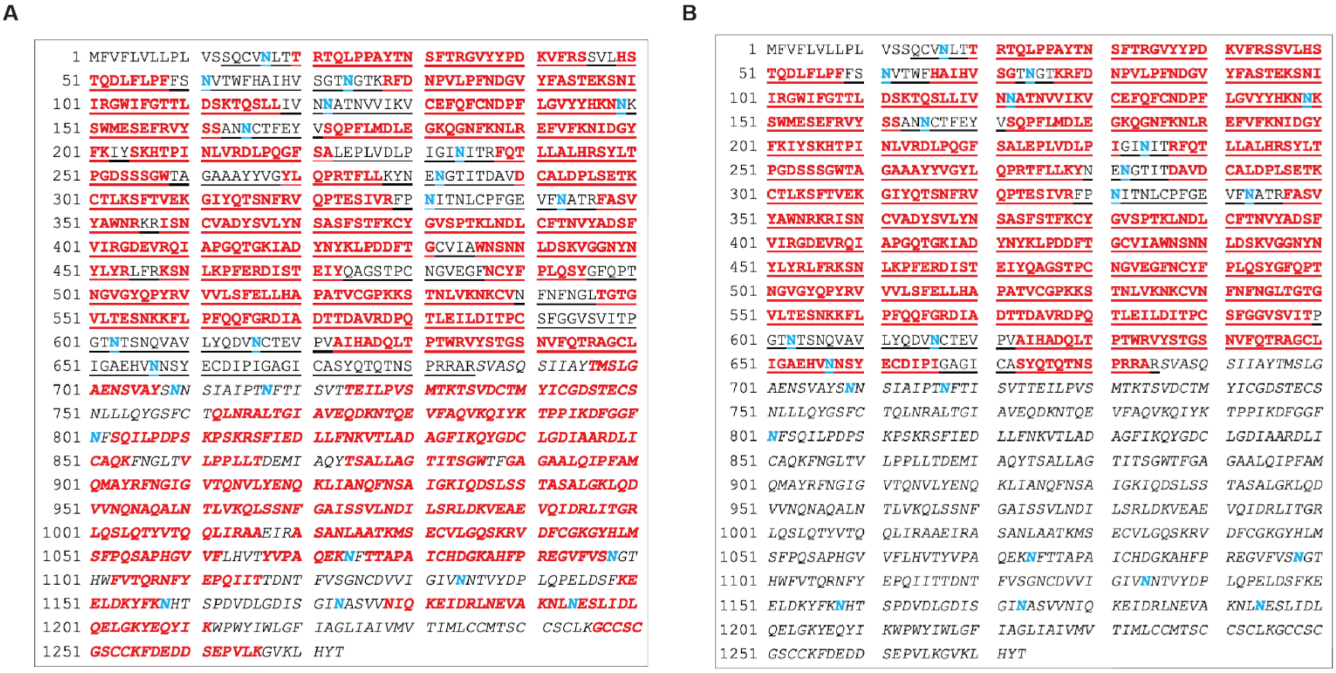

**Supplementary Figure 1:** Mascot sequence coverage for $S_{virus}$ and $S_{vaccine\ antigen}$. The overall sequence coverage for (A) $S_{virus}$ and (B) $S_{vaccine\ antigen}$ is shown for the three combined proteases (trypsin, chymotrypsin and alpha lytic protease) used for bottom-up MS. Over the sequence of S1, peptides covering 588 amino acids for $S_{virus}$ (88% coverage) and 586 amino acids for $S_{vaccine\ antigen}$ (87% coverage) were observed. Red bold: Peptides identified in Mascot. Blue bold: Potential N-glycosylation sites. Underlined: S1. Italics: S2. Not italics/not underlined: Signal peptide.

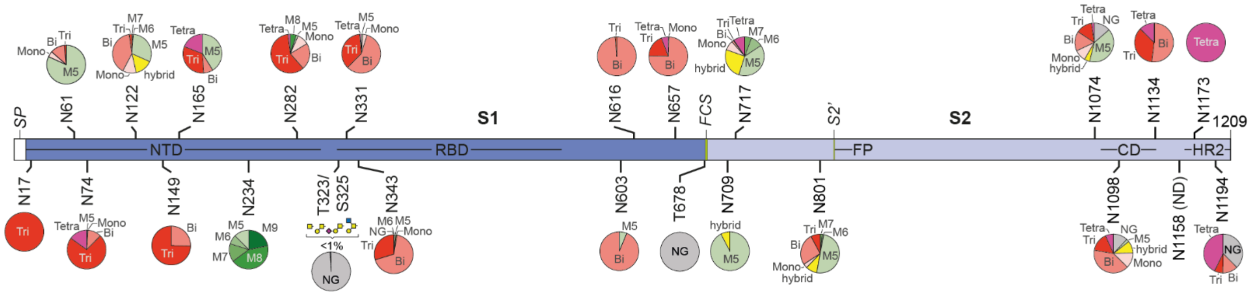

**Supplementary Figure 2**: Site-specific glycosylation of S<sub></sub>recombinant trimer.

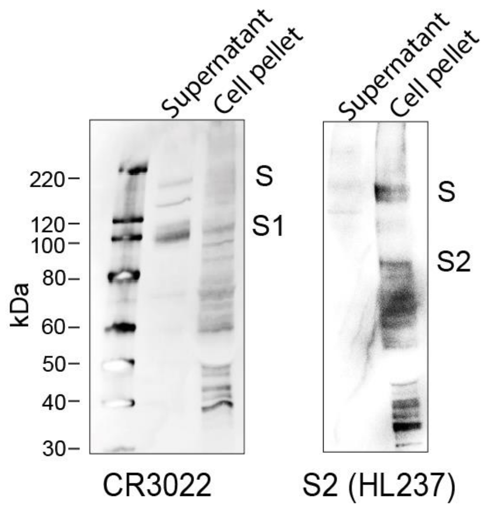

**Supplementary Figure 3**: Western blot of unpurified $S_{vaccine\ antigen}$ probed with anti-S1 (CR3022) and anti-S2 (HL237) antibody.

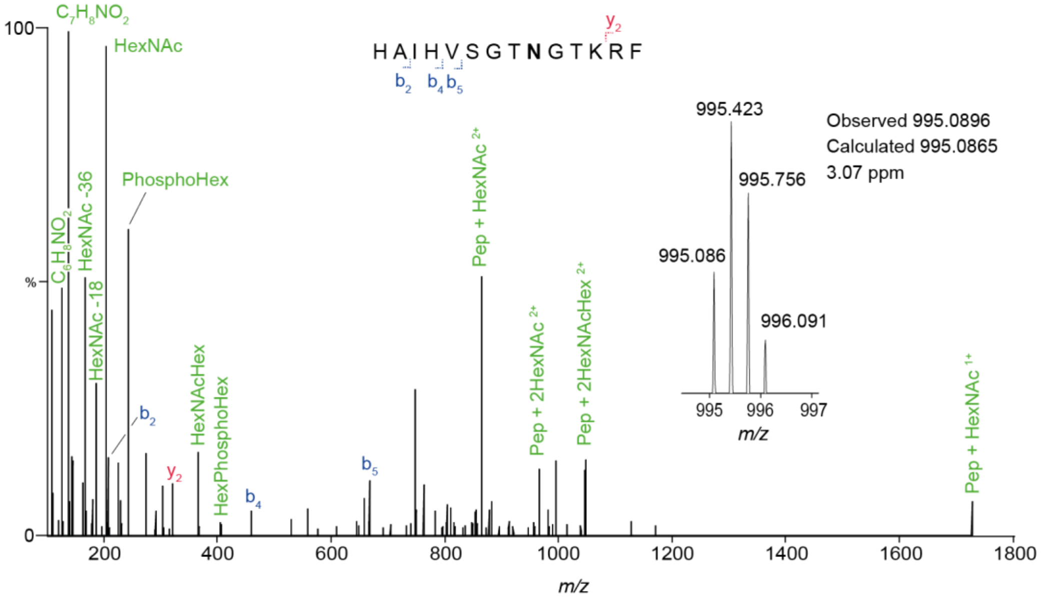

**Supplementary Figure 4**: MS/MS spectrum of the N74 glycopeptide from $S1_{vaccine}$ antigen

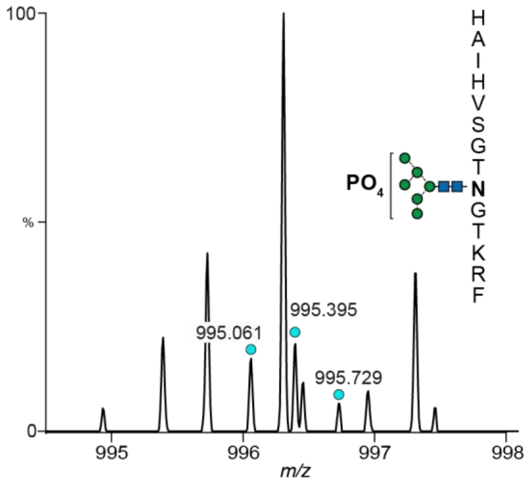

**Supplementary Figure 5**: MS spectrum showing the presence of the N74 N-glycopeptide (chymotrypsin digestion) carrying the mannose-6-phosphate glycan on $S1_{virus}$. The isotopic distribution is labelled (green circles).

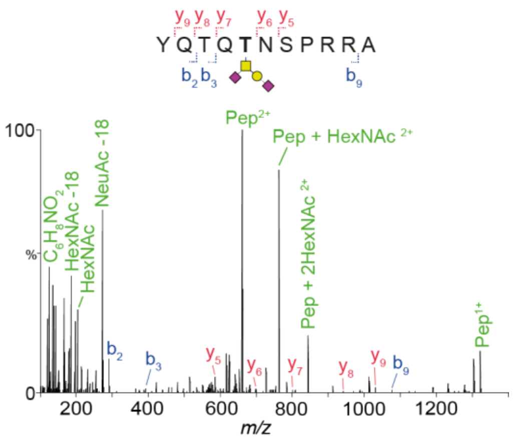

**Supplementary Figure 6**: MS/MS spectrum of the T678 glycopeptide from S1$_{vaccine}$ antigen.

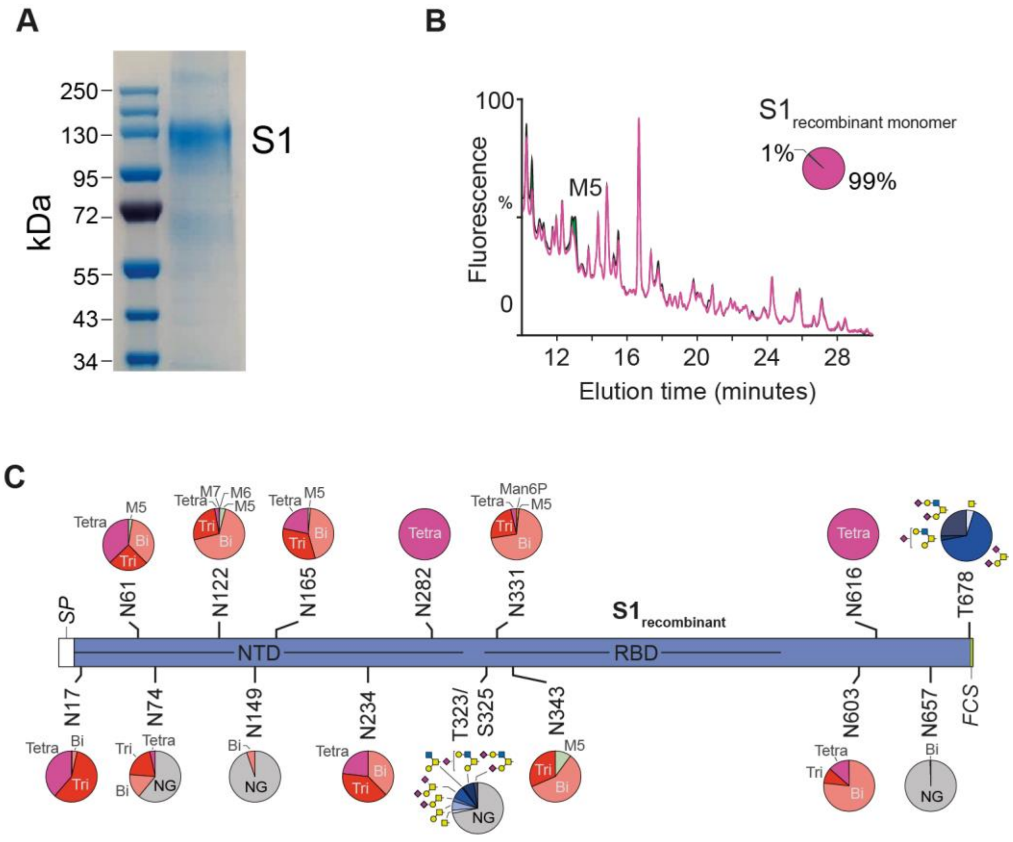

**Supplementary Figure 7**: Purification and glycan analysis of S1recombinant monomer. (A) SDS-gel of CR3022-purified S1reocmbinant monomer. (B) Quantitative HPLC N-glycan analysis of S1recombinant monomer showing distribution of complex (purple) and oligomannose (green) with the $GlcNAc_2Man_5$ (M5) structure labelled. (C) Site-specific glycosylation of Srecombinant monomer.

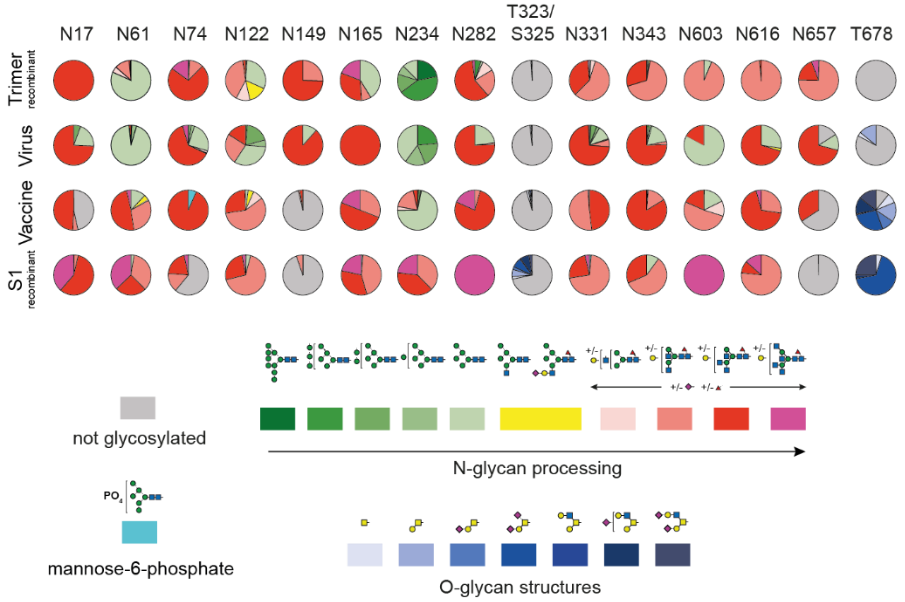

**Supplementary Figure 8**: Illustration of cumulative site-specific glycosylation changes across S1 subunits from recombinant trimer, virus, vaccine antigen and recombinant monomer samples.

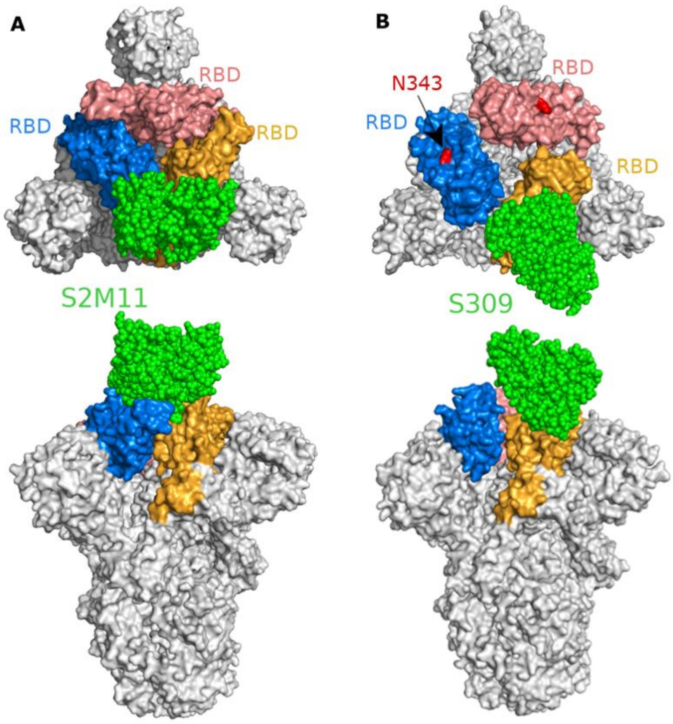

**Supplementary Figure 9: Structure and binding orientation of S2M11 and S309 neutralising antibodies to the spike glycoprotein**(A) S2M11 binds to a quaternary epitope formed by two neighbouring RBDs on the trimer apex (PDB ID: 7K43). (B) S309.binds to an epitope in RBD involving an N-linked glycan at position 343 (PDB ID: 6WPS). The three RBDs forming a trimer are shown in blue, yellow and peach colour. The neutralising antibodies are shown in green colour. N343 is shown in red color.

### Supplementary Tables

**Supplementary Table 1**: Identification of SARS-CoV-2 nucleoprotein and membrane protein in $S_{virus}$ sample following CR3022 purification.

| prot_acc | prot_desc | prot_score | pep_exp_mz | pep_exp_mr | pep_calc_mr | pep_exp_z | pep_delta | pep_score | pep_seq |
|---|---|---|---|---|---|---|---|---|---|
| P0DTC2 | Spike glycoprotein | 1206 | 376.7499 | 751.4852 | 751.4844 | 2 | 0.0008 | 10.52 | VLPPLLT |
| P0DTC2 | Spike glycoprotein | 1206 | 376.75 | 751.4855 | 751.4844 | 2 | 0.0011 | 6.28 | VLPPLLT |
| P0DTC2 | Spike glycoprotein | 1206 | 376.7502 | 751.4859 | 751.4844 | 2 | 0.0015 | 5.79 | VLPPLLT |
| P0DTC2 | Spike glycoprotein | 1206 | 376.7505 | 751.4864 | 751.4844 | 2 | 0.002 | 18.37 | VLPPLLT |
| P0DTC2 | Spike glycoprotein | 1206 | 376.7505 | 751.4864 | 751.4844 | 2 | 0.0021 | 15.77 | VLPPLLT |
| P0DTC2 | Spike glycoprotein | 1206 | 376.7505 | 751.4865 | 751.4844 | 2 | 0.0021 | 12.37 | VLPPLLT |
| P0DTC2 | Spike glycoprotein | 1206 | 376.7512 | 751.4878 | 751.4844 | 2 | 0.0035 | 1.68 | VLPPLLT |
| P0DTC2 | Spike glycoprotein | 1206 | 376.7519 | 751.4892 | 751.4844 | 2 | 0.0049 | 21.13 | VLPPLLT |
| P0DTC2 | Spike glycoprotein | 1206 | 377.1679 | 752.3213 | 752.3197 | 2 | 0.0016 | 6.2 | KMSECV |
| P0DTC2 | Spike glycoprotein | 1206 | 386.2046 | 770.3947 | 770.3923 | 2 | 0.0024 | 8.85 | SVLHSTQ |
| P0DTC2 | Spike glycoprotein | 1206 | 386.2218 | 770.429 | 770.4286 | 2 | 0.0004 | 18.09 | PGQTGKIA |
| P0DTC2 | Spike glycoprotein | 1206 | 387.6984 | 773.3823 | 773.3807 | 2 | 0.0016 | 14.31 | LDPLSET |
| P0DTC2 | Spike glycoprotein | 1206 | 387.6988 | 773.3831 | 773.3807 | 2 | 0.0024 | 5.08 | LDPLSET |
| P0DTC2 | Spike glycoprotein | 1206 | 389.1997 | 776.3848 | 776.3851 | 2 | -0.0003 | 8.21 | CGPKKST |
| P0DTC2 | Spike glycoprotein | 1206 | 389.1998 | 776.385 | 776.3851 | 2 | -0.0001 | 34.48 | CGPKKST |
| P0DTC2 | Spike glycoprotein | 1206 | 389.2176 | 776.4206 | 776.4181 | 2 | 0.0025 | 24.01 | YLQPRT |

| | | | | | | | | | |
|---|---|---|---|---|---|---|---|---|---|
| P0DTC2 | Spike glycoprotein | 1206 | 391.731 | 781.4475 | 781.4446 | 2 | 0.0028 | 14.31 | RTQLPPA |
| P0DTC2 | Spike glycoprotein | 1206 | 394.2436 | 786.4727 | 786.4712 | 2 | 0.0015 | 25.48 | LGQSKRV |
| P0DTC2 | Spike glycoprotein | 1206 | 395.218 | 788.4214 | 788.4215 | 2 | -0.0001 | 8.99 | TVCGPKK |
| P0DTC2 | Spike glycoprotein | 1206 | 396.199 | 790.3834 | 790.3821 | 2 | 0.0013 | 14.14 | RDIADTT |
| P0DTC2 | Spike glycoprotein | 1206 | 396.1995 | 790.3844 | 790.3821 | 2 | 0.0023 | 22.19 | RDIADTT |
| P0DTC2 | Spike glycoprotein | 1206 | 396.1995 | 790.3845 | 790.3821 | 2 | 0.0024 | 40.54 | RDIADTT |
| P0DTC2 | Spike glycoprotein | 1206 | 396.7087 | 791.4029 | 791.4025 | 2 | 0.0004 | 13.81 | TLDSKTQ |
| P0DTC2 | Spike glycoprotein | 1206 | 398.7138 | 795.4131 | 795.4127 | 2 | 0.0005 | 5.43 | KGIYQTS |
| P0DTC2 | Spike glycoprotein | 1206 | 400.6909 | 799.3673 | 799.3647 | 2 | 0.0026 | 3.95 | ICHDGKA |
| P0DTC2 | Spike glycoprotein | 1206 | 400.7 | 799.3855 | 799.3824 | 2 | 0.0031 | 33.65 | DAVRDPQ |
| P0DTC2 | Spike glycoprotein | 1206 | 403.7545 | 805.4944 | 805.4922 | 2 | 0.0022 | 6.85 | RLFRKS |
| P0DTC2 | Spike glycoprotein | 1206 | 405.2543 | 808.494 | 808.4919 | 2 | 0.0021 | 22.2 | LLALHRS |
| P0DTC2 | Spike glycoprotein | 1206 | 406.2091 | 810.4036 | 810.4024 | 2 | 0.0011 | 13.51 | WRVYST |
| P0DTC2 | Spike glycoprotein | 1206 | 406.2092 | 810.4038 | 810.4024 | 2 | 0.0014 | 9.26 | WRVYST |
| P0DTC2 | Spike glycoprotein | 1206 | 406.2095 | 810.4045 | 810.4024 | 2 | 0.002 | 3.89 | WRVYST |
| P0DTC2 | Spike glycoprotein | 1206 | 406.2098 | 810.405 | 810.4024 | 2 | 0.0026 | 10.54 | WRVYST |
| P0DTC2 | Spike glycoprotein | 1206 | 406.2098 | 810.4051 | 810.4024 | 2 | 0.0026 | 17.31 | WRVYST |
| P0DTC2 | Spike glycoprotein | 1206 | 412.2085 | 822.4024 | 822.4024 | 2 | 0 | 2.86 | VYYHKN |
| P0DTC2 | Spike glycoprotein | 1206 | 418.7257 | 835.4368 | 835.4341 | 2 | 0.0027 | 11.56 | RGWIFGT |
| P0DTC2 | Spike glycoprotein | 1206 | 419.7199 | 837.4252 | 837.4232 | 2 | 0.002 | 13.53 | EKGIYQT |
| P0DTC2 | Spike glycoprotein | 1206 | 430.756 | 859.4975 | 859.4949 | 2 | 0.0026 | 16.84 | LVKNKCV |

| | | | | | | | | | |
|---|---|---|---|---|---|---|---|---|---|
| P0DTC2 | Spike glycoprotein | 1206 | 431.2336 | 860.4526 | 860.4505 | 2 | 0.0022 | 14.07 | NFRVQPT |
| P0DTC2 | Spike glycoprotein | 1206 | 431.2338 | 860.4531 | 860.4505 | 2 | 0.0026 | 14.26 | NFRVQPT |
| P0DTC2 | Spike glycoprotein | 1206 | 431.2343 | 860.454 | 860.4505 | 2 | 0.0035 | 22.49 | NFRVQPT |
| P0DTC2 | Spike glycoprotein | 1206 | 431.2343 | 860.454 | 860.4505 | 2 | 0.0036 | 24.3 | NFRVQPT |
| P0DTC2 | Spike glycoprotein | 1206 | 434.7303 | 867.4461 | 867.445 | 2 | 0.0011 | 6.32 | AIHADQLT |
| P0DTC2 | Spike glycoprotein | 1206 | 434.7306 | 867.4467 | 867.445 | 2 | 0.0017 | 25.51 | AIHADQLT |
| P0DTC2 | Spike glycoprotein | 1206 | 434.7308 | 867.4471 | 867.445 | 2 | 0.0021 | 14.49 | AIHADQLT |
| P0DTC2 | Spike glycoprotein | 1206 | 434.7313 | 867.448 | 867.445 | 2 | 0.0029 | 26 | AIHADQLT |
| P0DTC2 | Spike glycoprotein | 1206 | 434.7313 | 867.448 | 867.445 | 2 | 0.003 | 7.26 | AIHADQLT |
| P0DTC2 | Spike glycoprotein | 1206 | 434.7313 | 867.4481 | 867.445 | 2 | 0.003 | 20.56 | AIHADQLT |
| P0DTC2 | Spike glycoprotein | 1206 | 434.7313 | 867.4481 | 867.445 | 2 | 0.0031 | 16.62 | AIHADQLT |
| P0DTC2 | Spike glycoprotein | 1206 | 434.7315 | 867.4485 | 867.445 | 2 | 0.0035 | 17.78 | AIHADQLT |
| P0DTC2 | Spike glycoprotein | 1206 | 434.7315 | 867.4485 | 867.445 | 2 | 0.0035 | 9.95 | AIHADQLT |
| P0DTC2 | Spike glycoprotein | 1206 | 434.7316 | 867.4486 | 867.445 | 2 | 0.0036 | 3.43 | AIHADQLT |
| P0DTC2 | Spike glycoprotein | 1206 | 435.7447 | 869.4748 | 869.4719 | 2 | 0.0029 | 45.19 | RQIAPGQT |
| P0DTC2 | Spike glycoprotein | 1206 | 435.745 | 869.4755 | 869.4719 | 2 | 0.0036 | 27.9 | RQIAPGQT |
| P0DTC2 | Spike glycoprotein | 1206 | 438.2431 | 874.4717 | 874.4695 | 2 | 0.0022 | 13.94 | NLVKNKC |
| P0DTC2 | Spike glycoprotein | 1206 | 438.7339 | 875.4533 | 875.4535 | 2 | -0.0002 | 27.2 | VCGPKKST |
| P0DTC2 | Spike glycoprotein | 1206 | 438.7349 | 875.4552 | 875.4535 | 2 | 0.0017 | 29.98 | TVCGPKKS |
| P0DTC2 | Spike glycoprotein | 1206 | 441.7207 | 881.4269 | 881.4243 | 2 | 0.0026 | 28.36 | HADQLTPT |
| P0DTC2 | Spike glycoprotein | 1206 | 444.2171 | 886.4197 | 886.4185 | 2 | 0.0012 | 4.28 | EFRVYSS |

| | | | | | | | | | |
|---|---|---|---|---|---|---|---|---|---|
| P0DTC2 | Spike glycoprotein | 1206 | 444.2172 | 886.4198 | 886.4185 | 2 | 0.0013 | 5.33 | EFRVYSS |
| P0DTC2 | Spike glycoprotein | 1206 | 444.2172 | 886.4199 | 886.4185 | 2 | 0.0014 | 5.12 | EFRVYSS |
| P0DTC2 | Spike glycoprotein | 1206 | 444.2175 | 886.4203 | 886.4185 | 2 | 0.0019 | 4.63 | EFRVYSS |
| P0DTC2 | Spike glycoprotein | 1206 | 444.2175 | 886.4205 | 886.4185 | 2 | 0.002 | 22.93 | EFRVYSS |
| P0DTC2 | Spike glycoprotein | 1206 | 444.2176 | 886.4206 | 886.4185 | 2 | 0.0022 | 13.67 | EFRVYSS |
| P0DTC2 | Spike glycoprotein | 1206 | 444.2177 | 886.4208 | 886.4185 | 2 | 0.0023 | 7.86 | EFRVYSS |
| P0DTC2 | Spike glycoprotein | 1206 | 444.2177 | 886.4209 | 886.4185 | 2 | 0.0025 | 9.47 | EFRVYSS |
| P0DTC2 | Spike glycoprotein | 1206 | 444.2179 | 886.4213 | 886.4185 | 2 | 0.0028 | 12.41 | EFRVYSS |
| P0DTC2 | Spike glycoprotein | 1206 | 444.218 | 886.4214 | 886.4185 | 2 | 0.003 | 11.31 | EFRVYSS |
| P0DTC2 | Spike glycoprotein | 1206 | 444.2182 | 886.4218 | 886.4185 | 2 | 0.0033 | 4.23 | EFRVYSS |
| P0DTC2 | Spike glycoprotein | 1206 | 444.2184 | 886.4222 | 886.4185 | 2 | 0.0038 | 7.45 | EFRVYSS |
| P0DTC2 | Spike glycoprotein | 1206 | 444.2185 | 886.4224 | 886.4185 | 2 | 0.004 | 4.51 | EFRVYSS |
| P0DTC2 | Spike glycoprotein | 1206 | 445.2491 | 888.4837 | 888.4817 | 2 | 0.002 | 23.32 | LHRSYLT |
| P0DTC2 | Spike glycoprotein | 1206 | 451.2225 | 900.4305 | 900.4301 | 2 | 0.0003 | 5.48 | DAVRDPQT |
| P0DTC2 | Spike glycoprotein | 1206 | 451.2238 | 900.4331 | 900.4301 | 2 | 0.003 | 21.97 | DAVRDPQT |
| P0DTC2 | Spike glycoprotein | 1206 | 451.224 | 900.4335 | 900.4301 | 2 | 0.0034 | 24.49 | DAVRDPQT |
| P0DTC2 | Spike glycoprotein | 1206 | 451.2241 | 900.4336 | 900.4301 | 2 | 0.0035 | 14.81 | DAVRDPQT |
| P0DTC2 | Spike glycoprotein | 1206 | 455.2407 | 908.4669 | 908.4657 | 2 | 0.0012 | 18.2 | WFHAIHV |
| P0DTC2 | Spike glycoprotein | 1206 | 455.2408 | 908.4671 | 908.4657 | 2 | 0.0013 | 21.18 | WFHAIHV |
| P0DTC2 | Spike glycoprotein | 1206 | 455.2411 | 908.4676 | 908.4657 | 2 | 0.0019 | 16.95 | WFHAIHV |
| P0DTC2 | Spike glycoprotein | 1206 | 456.2206 | 910.4267 | 910.4219 | 2 | 0.0049 | 27.94 | KCYGVSPT |

| | | | | | | | | | |
|---|---|---|---|---|---|---|---|---|---|
| P0DTC2 | Spike glycoprotein | 1206 | 456.7444 | 911.4742 | 911.4713 | 2 | 0.003 | 3.16 | VLHSTQDL |
| P0DTC2 | Spike glycoprotein | 1206 | 458.7659 | 915.5172 | 915.5137 | 2 | 0.0035 | 22.92 | EKSNIIRG |
| P0DTC2 | Spike glycoprotein | 1206 | 460.2367 | 918.4588 | 918.4559 | 2 | 0.0028 | 26.71 | RDLPQGFS |
| P0DTC2 | Spike glycoprotein | 1206 | 460.237 | 918.4594 | 918.4559 | 2 | 0.0035 | 37.11 | RDLPQGFS |
| P0DTC2 | Spike glycoprotein | 1206 | 460.2371 | 918.4596 | 918.4559 | 2 | 0.0036 | 33.46 | RDLPQGFS |
| P0DTC2 | Spike glycoprotein | 1206 | 462.7503 | 923.486 | 923.4865 | 2 | -0.0005 | 5.38 | YLQPRTF |
| P0DTC2 | Spike glycoprotein | 1206 | 462.7508 | 923.487 | 923.4865 | 2 | 0.0005 | 10.8 | YQPYRVV |
| P0DTC2 | Spike glycoprotein | 1206 | 462.7508 | 923.487 | 923.4865 | 2 | 0.0006 | 23.96 | YLQPRTF |
| P0DTC2 | Spike glycoprotein | 1206 | 462.751 | 923.4875 | 923.4865 | 2 | 0.001 | 7.36 | YLQPRTF |
| P0DTC2 | Spike glycoprotein | 1206 | 462.7511 | 923.4877 | 923.4865 | 2 | 0.0012 | 4.65 | YLQPRTF |
| P0DTC2 | Spike glycoprotein | 1206 | 462.7511 | 923.4877 | 923.4865 | 2 | 0.0012 | 15.57 | YQPYRVV |
| P0DTC2 | Spike glycoprotein | 1206 | 462.7512 | 923.4878 | 923.4865 | 2 | 0.0013 | 5.29 | YLQPRTF |
| P0DTC2 | Spike glycoprotein | 1206 | 462.7512 | 923.4879 | 923.4865 | 2 | 0.0014 | 4.51 | YLQPRTF |
| P0DTC2 | Spike glycoprotein | 1206 | 462.7512 | 923.4879 | 923.4865 | 2 | 0.0014 | 6.31 | YQPYRVV |
| P0DTC2 | Spike glycoprotein | 1206 | 462.7512 | 923.4879 | 923.4865 | 2 | 0.0014 | 4.99 | YLQPRTF |
| P0DTC2 | Spike glycoprotein | 1206 | 462.7512 | 923.4879 | 923.4865 | 2 | 0.0014 | 10.27 | YLQPRTF |
| P0DTC2 | Spike glycoprotein | 1206 | 462.7513 | 923.488 | 923.4865 | 2 | 0.0015 | 10.43 | YLQPRTF |
| P0DTC2 | Spike glycoprotein | 1206 | 462.7513 | 923.488 | 923.4865 | 2 | 0.0015 | 3.3 | YLQPRTF |
| P0DTC2 | Spike glycoprotein | 1206 | 462.7513 | 923.4881 | 923.4865 | 2 | 0.0016 | 20.05 | YLQPRTF |
| P0DTC2 | Spike glycoprotein | 1206 | 462.7514 | 923.4882 | 923.4865 | 2 | 0.0017 | 6.43 | YLQPRTF |
| P0DTC2 | Spike glycoprotein | 1206 | 462.7514 | 923.4882 | 923.4865 | 2 | 0.0017 | 5.84 | YQPYRVV |

| | | | | | | | | | |
|---|---|---|---|---|---|---|---|---|---|
| P0DTC2 | Spike glycoprotein | 1206 | 462.7514 | 923.4883 | 923.4865 | 2 | 0.0018 | 10.54 | YLQPRTF |
| P0DTC2 | Spike glycoprotein | 1206 | 462.7515 | 923.4884 | 923.4865 | 2 | 0.0019 | 7.67 | YQPYRVV |
| P0DTC2 | Spike glycoprotein | 1206 | 462.7515 | 923.4884 | 923.4865 | 2 | 0.0019 | 7.85 | YQPYRVV |
| P0DTC2 | Spike glycoprotein | 1206 | 462.7515 | 923.4884 | 923.4865 | 2 | 0.0019 | 8.3 | YLQPRTF |
| P0DTC2 | Spike glycoprotein | 1206 | 462.7515 | 923.4884 | 923.4865 | 2 | 0.0019 | 6.46 | YQPYRVV |
| P0DTC2 | Spike glycoprotein | 1206 | 462.7516 | 923.4887 | 923.4865 | 2 | 0.0022 | 10.59 | YLQPRTF |
| P0DTC2 | Spike glycoprotein | 1206 | 462.7517 | 923.4888 | 923.4865 | 2 | 0.0023 | 1.98 | YLQPRTF |
| P0DTC2 | Spike glycoprotein | 1206 | 462.7517 | 923.4888 | 923.4865 | 2 | 0.0023 | 10.32 | YLQPRTF |
| P0DTC2 | Spike glycoprotein | 1206 | 462.7517 | 923.4888 | 923.4865 | 2 | 0.0023 | 9.88 | YLQPRTF |
| P0DTC2 | Spike glycoprotein | 1206 | 462.7518 | 923.489 | 923.4865 | 2 | 0.0025 | 2.37 | YLQPRTF |
| P0DTC2 | Spike glycoprotein | 1206 | 462.7518 | 923.489 | 923.4865 | 2 | 0.0025 | 8.26 | YQPYRVV |
| P0DTC2 | Spike glycoprotein | 1206 | 462.7518 | 923.4891 | 923.4865 | 2 | 0.0026 | 10.14 | YLQPRTF |
| P0DTC2 | Spike glycoprotein | 1206 | 462.7519 | 923.4892 | 923.4865 | 2 | 0.0027 | 12.87 | YLQPRTF |
| P0DTC2 | Spike glycoprotein | 1206 | 462.752 | 923.4894 | 923.4865 | 2 | 0.0029 | 10.38 | YLQPRTF |
| P0DTC2 | Spike glycoprotein | 1206 | 462.752 | 923.4894 | 923.4865 | 2 | 0.0029 | 11.8 | YQPYRVV |
| P0DTC2 | Spike glycoprotein | 1206 | 462.752 | 923.4895 | 923.4865 | 2 | 0.003 | 34.8 | YLQPRTF |
| P0DTC2 | Spike glycoprotein | 1206 | 462.752 | 923.4895 | 923.4865 | 2 | 0.003 | 10.21 | YQPYRVV |
| P0DTC2 | Spike glycoprotein | 1206 | 462.7521 | 923.4897 | 923.4865 | 2 | 0.0032 | 10.4 | YLQPRTF |
| P0DTC2 | Spike glycoprotein | 1206 | 462.7521 | 923.4897 | 923.4865 | 2 | 0.0032 | 7.74 | YQPYRVV |
| P0DTC2 | Spike glycoprotein | 1206 | 462.7522 | 923.4898 | 923.4865 | 2 | 0.0033 | 8.8 | YLQPRTF |
| P0DTC2 | Spike glycoprotein | 1206 | 462.7523 | 923.4901 | 923.4865 | 2 | 0.0036 | 12.06 | YLQPRTF |

| | | | | | | | | | |
|---|---|---|---|---|---|---|---|---|---|
| P0DTC2 | Spike glycoprotein | 1206 | 462.7526 | 923.4907 | 923.4865 | 2 | 0.0042 | 5.77 | YQPYRVV |
| P0DTC2 | Spike glycoprotein | 1206 | 462.7528 | 923.491 | 923.4865 | 2 | 0.0045 | 5.28 | YLQPRTF |
| P0DTC2 | Spike glycoprotein | 1206 | 462.7541 | 923.4937 | 923.4865 | 2 | 0.0072 | 8.96 | YLQPRTF |
| P0DTC2 | Spike glycoprotein | 1206 | 463.2363 | 924.458 | 924.4552 | 2 | 0.0028 | 16.27 | EKGIYQTS |
| P0DTC2 | Spike glycoprotein | 1206 | 463.2367 | 924.4588 | 924.4552 | 2 | 0.0036 | 6.82 | EKGIYQTS |
| P0DTC2 | Spike glycoprotein | 1206 | 463.2367 | 924.4588 | 924.4552 | 2 | 0.0036 | 6.08 | EKGIYQTS |
| P0DTC2 | Spike glycoprotein | 1206 | 463.7229 | 925.4313 | 925.4294 | 2 | 0.0019 | 7.61 | NFNFNGLT |
| P0DTC2 | Spike glycoprotein | 1206 | 463.723 | 925.4314 | 925.4294 | 2 | 0.0021 | 9.09 | NFNFNGLT |
| P0DTC2 | Spike glycoprotein | 1206 | 464.2558 | 926.497 | 926.4934 | 2 | 0.0036 | 28.46 | RQIAPGQTG |
| P0DTC2 | Spike glycoprotein | 1206 | 467.7542 | 933.4939 | 933.492 | 2 | 0.0019 | 11.6 | FVIRGDEV |
| P0DTC2 | Spike glycoprotein | 1206 | 467.7547 | 933.4949 | 933.492 | 2 | 0.0029 | 16.36 | FVIRGDEV |
| P0DTC2 | Spike glycoprotein | 1206 | 467.7551 | 933.4956 | 933.492 | 2 | 0.0036 | 17.19 | FVIRGDEV |
| P0DTC2 | Spike glycoprotein | 1206 | 469.2492 | 936.4839 | 936.4818 | 2 | 0.0022 | 32.02 | RGWIFGTT |
| P0DTC2 | Spike glycoprotein | 1206 | 469.2493 | 936.4841 | 936.4818 | 2 | 0.0023 | 36.31 | RGWIFGTT |
| P0DTC2 | Spike glycoprotein | 1206 | 474.7503 | 947.4861 | 947.4825 | 2 | 0.0036 | 25.92 | SNFRVQPT |
| P0DTC2 | Spike glycoprotein | 1206 | 480.7682 | 959.5218 | 959.5188 | 2 | 0.003 | 15.9 | ALHRSYLT |
| P0DTC2 | Spike glycoprotein | 1206 | 485.2867 | 968.5588 | 968.5556 | 2 | 0.0033 | 1.08 | YRLFRKS |
| P0DTC2 | Spike glycoprotein | 1206 | 487.7774 | 973.5402 | 973.5379 | 2 | 0.0023 | 33.22 | NLVKNKCV |
| P0DTC2 | Spike glycoprotein | 1206 | 487.7774 | 973.5402 | 973.5379 | 2 | 0.0023 | 25.43 | NLVKNKCV |
| P0DTC2 | Spike glycoprotein | 1206 | 487.7774 | 973.5402 | 973.5379 | 2 | 0.0023 | 38.71 | NLVKNKCV |
| P0DTC2 | Spike glycoprotein | 1206 | 487.7774 | 973.5403 | 973.5379 | 2 | 0.0024 | 32.69 | NLVKNKCV |

| | | | | | | | | | |
|---|---|---|---|---|---|---|---|---|---|
| P0DTC2 | Spike glycoprotein | 1206 | 487.7776 | 973.5406 | 973.5379 | 2 | 0.0027 | 16.97 | NLVKNKCV |
| P0DTC2 | Spike glycoprotein | 1206 | 487.7776 | 973.5407 | 973.5379 | 2 | 0.0028 | 33.14 | NLVKNKCV |
| P0DTC2 | Spike glycoprotein | 1206 | 487.7777 | 973.5409 | 973.5379 | 2 | 0.003 | 16.42 | NLVKNKCV |
| P0DTC2 | Spike glycoprotein | 1206 | 487.7778 | 973.541 | 973.5379 | 2 | 0.0031 | 32.64 | NLVKNKCV |
| P0DTC2 | Spike glycoprotein | 1206 | 487.7779 | 973.5412 | 973.5379 | 2 | 0.0034 | 28.42 | NLVKNKCV |
| P0DTC2 | Spike glycoprotein | 1206 | 487.778 | 973.5414 | 973.5379 | 2 | 0.0035 | 28.24 | NLVKNKCV |
| P0DTC2 | Spike glycoprotein | 1206 | 487.7781 | 973.5416 | 973.5379 | 2 | 0.0037 | 32.55 | NLVKNKCV |
| P0DTC2 | Spike glycoprotein | 1206 | 487.7782 | 973.5418 | 973.5379 | 2 | 0.004 | 33.48 | NLVKNKCV |
| P0DTC2 | Spike glycoprotein | 1206 | 487.7784 | 973.5423 | 973.5379 | 2 | 0.0045 | 11.33 | NLVKNKCV |
| P0DTC2 | Spike glycoprotein | 1206 | 489.258 | 976.5014 | 976.5012 | 2 | 0.0002 | 32.52 | TVCGPKKST |
| P0DTC2 | Spike glycoprotein | 1206 | 489.2584 | 976.5023 | 976.5012 | 2 | 0.0012 | 11.8 | TVCGPKKST |
| P0DTC2 | Spike glycoprotein | 1206 | 489.2586 | 976.5026 | 976.5012 | 2 | 0.0014 | 34.8 | TVCGPKKST |
| P0DTC2 | Spike glycoprotein | 1206 | 489.2586 | 976.5027 | 976.5012 | 2 | 0.0015 | 35.16 | TVCGPKKST |
| P0DTC2 | Spike glycoprotein | 1206 | 489.2587 | 976.5028 | 976.5012 | 2 | 0.0016 | 29.99 | TVCGPKKST |
| P0DTC2 | Spike glycoprotein | 1206 | 489.259 | 976.5035 | 976.5012 | 2 | 0.0023 | 18.86 | TVCGPKKST |
| P0DTC2 | Spike glycoprotein | 1206 | 489.2591 | 976.5037 | 976.5012 | 2 | 0.0025 | 7.81 | TVCGPKKST |
| P0DTC2 | Spike glycoprotein | 1206 | 489.2593 | 976.504 | 976.5012 | 2 | 0.0029 | 3.64 | TVCGPKKST |
| P0DTC2 | Spike glycoprotein | 1206 | 489.2593 | 976.504 | 976.5012 | 2 | 0.0029 | 33.17 | TVCGPKKST |
| P0DTC2 | Spike glycoprotein | 1206 | 491.262 | 980.5094 | 980.508 | 2 | 0.0015 | 17.3 | GYLQPRTF |
| P0DTC2 | Spike glycoprotein | 1206 | 492.7606 | 983.5066 | 983.5036 | 2 | 0.003 | 30.13 | DEVRQIAPG |
| P0DTC2 | Spike glycoprotein | 1206 | 492.7614 | 983.5083 | 983.5036 | 2 | 0.0047 | 18.57 | DEVRQIAPG |

| | | | | | | | | | |
|---|---|---|---|---|---|---|---|---|---|
| P0DTC2 | Spike glycoprotein | 1206 | 494.3043 | 986.594 | 986.5913 | 2 | 0.0027 | 10.86 | LQPRTFLL |
| P0DTC2 | Spike glycoprotein | 1206 | 495.7555 | 989.4965 | 989.493 | 2 | 0.0035 | 20.2 | RDLPQGFSA |
| P0DTC2 | Spike glycoprotein | 1206 | 496.266 | 990.5175 | 990.5134 | 2 | 0.0041 | 12.09 | KPFERDIS |
| P0DTC2 | Spike glycoprotein | 1206 | 498.2634 | 994.5122 | 994.5084 | 2 | 0.0038 | 36.21 | IHADQLTPT |
| P0DTC2 | Spike glycoprotein | 1206 | 498.7567 | 995.4988 | 995.4977 | 2 | 0.001 | 16.57 | WFHAIHVS |
| P0DTC2 | Spike glycoprotein | 1206 | 498.7574 | 995.5002 | 995.4977 | 2 | 0.0024 | 15.58 | WFHAIHVS |
| P0DTC2 | Spike glycoprotein | 1206 | 498.7576 | 995.5007 | 995.4977 | 2 | 0.0029 | 5.96 | WFHAIHVS |
| P0DTC2 | Spike glycoprotein | 1206 | 498.7578 | 995.501 | 995.4977 | 2 | 0.0032 | 9.91 | WFHAIHVS |
| P0DTC2 | Spike glycoprotein | 1206 | 498.7579 | 995.5012 | 995.4977 | 2 | 0.0034 | 15.26 | WFHAIHVS |
| P0DTC2 | Spike glycoprotein | 1206 | 498.7581 | 995.5017 | 995.4977 | 2 | 0.0039 | 15.98 | WFHAIHVS |
| P0DTC2 | Spike glycoprotein | 1206 | 498.7581 | 995.5017 | 995.4977 | 2 | 0.004 | 15.85 | WFHAIHVS |
| P0DTC2 | Spike glycoprotein | 1206 | 498.7593 | 995.5041 | 995.4977 | 2 | 0.0064 | 14.91 | WFHAIHVS |
| P0DTC2 | Spike glycoprotein | 1206 | 504.8047 | 1007.5949 | 1007.5903 | 2 | 0.0046 | 3.02 | LEPLVDLPI |
| P0DTC2 | Spike glycoprotein | 1206 | 505.7529 | 1009.4913 | 1009.4903 | 2 | 0.001 | 23.2 | KLNDLCFT |
| P0DTC2 | Spike glycoprotein | 1206 | 505.7532 | 1009.4918 | 1009.4903 | 2 | 0.0015 | 14.95 | KLNDLCFT |
| P0DTC2 | Spike glycoprotein | 1206 | 505.7532 | 1009.4919 | 1009.4903 | 2 | 0.0016 | 20.53 | KLNDLCFT |
| P0DTC2 | Spike glycoprotein | 1206 | 505.7533 | 1009.492 | 1009.4903 | 2 | 0.0017 | 17.13 | KLNDLCFT |
| P0DTC2 | Spike glycoprotein | 1206 | 505.7533 | 1009.4921 | 1009.4903 | 2 | 0.0018 | 14.67 | KLNDLCFT |
| P0DTC2 | Spike glycoprotein | 1206 | 505.7534 | 1009.4922 | 1009.4903 | 2 | 0.0019 | 30.35 | KLNDLCFT |
| P0DTC2 | Spike glycoprotein | 1206 | 505.7534 | 1009.4922 | 1009.4903 | 2 | 0.0019 | 30.09 | KLNDLCFT |
| P0DTC2 | Spike glycoprotein | 1206 | 505.7534 | 1009.4923 | 1009.4903 | 2 | 0.0021 | 33.7 | KLNDLCFT |

| | | | | | | | | | |
|---|---|---|---|---|---|---|---|---|---|
| P0DTC2 | Spike glycoprotein | 1206 | 505.7535 | 1009.4924 | 1009.4903 | 2 | 0.0022 | 15.35 | KLNDLCFT |
| P0DTC2 | Spike glycoprotein | 1206 | 505.7535 | 1009.4925 | 1009.4903 | 2 | 0.0023 | 24.22 | KLNDLCFT |
| P0DTC2 | Spike glycoprotein | 1206 | 505.7536 | 1009.4927 | 1009.4903 | 2 | 0.0024 | 17.97 | KLNDLCFT |
| P0DTC2 | Spike glycoprotein | 1206 | 505.7537 | 1009.4928 | 1009.4903 | 2 | 0.0026 | 22.97 | KLNDLCFT |
| P0DTC2 | Spike glycoprotein | 1206 | 505.7537 | 1009.4929 | 1009.4903 | 2 | 0.0026 | 15.7 | KLNDLCFT |
| P0DTC2 | Spike glycoprotein | 1206 | 505.7538 | 1009.4931 | 1009.4903 | 2 | 0.0029 | 23.27 | KLNDLCFT |
| P0DTC2 | Spike glycoprotein | 1206 | 505.754 | 1009.4934 | 1009.4903 | 2 | 0.0032 | 27.47 | KLNDLCFT |
| P0DTC2 | Spike glycoprotein | 1206 | 505.754 | 1009.4934 | 1009.4903 | 2 | 0.0032 | 17.95 | KLNDLCFT |
| P0DTC2 | Spike glycoprotein | 1206 | 505.754 | 1009.4935 | 1009.4903 | 2 | 0.0032 | 16.69 | KLNDLCFT |
| P0DTC2 | Spike glycoprotein | 1206 | 505.754 | 1009.4935 | 1009.4903 | 2 | 0.0033 | 8.16 | KLNDLCFT |
| P0DTC2 | Spike glycoprotein | 1206 | 505.7541 | 1009.4936 | 1009.4903 | 2 | 0.0033 | 16.33 | KLNDLCFT |
| P0DTC2 | Spike glycoprotein | 1206 | 505.7541 | 1009.4936 | 1009.4903 | 2 | 0.0034 | 12.85 | KLNDLCFT |
| P0DTC2 | Spike glycoprotein | 1206 | 505.7541 | 1009.4936 | 1009.4903 | 2 | 0.0034 | 19.86 | KLNDLCFT |
| P0DTC2 | Spike glycoprotein | 1206 | 505.7541 | 1009.4936 | 1009.4903 | 2 | 0.0034 | 23.38 | KLNDLCFT |
| P0DTC2 | Spike glycoprotein | 1206 | 505.7541 | 1009.4937 | 1009.4903 | 2 | 0.0034 | 19.49 | KLNDLCFT |
| P0DTC2 | Spike glycoprotein | 1206 | 505.7541 | 1009.4937 | 1009.4903 | 2 | 0.0034 | 23.69 | KLNDLCFT |
| P0DTC2 | Spike glycoprotein | 1206 | 505.7541 | 1009.4937 | 1009.4903 | 2 | 0.0035 | 21.56 | KLNDLCFT |
| P0DTC2 | Spike glycoprotein | 1206 | 505.7541 | 1009.4937 | 1009.4903 | 2 | 0.0035 | 19.4 | KLNDLCFT |
| P0DTC2 | Spike glycoprotein | 1206 | 505.7542 | 1009.4939 | 1009.4903 | 2 | 0.0037 | 18.12 | KLNDLCFT |
| P0DTC2 | Spike glycoprotein | 1206 | 505.7543 | 1009.4941 | 1009.4903 | 2 | 0.0038 | 12.31 | KLNDLCFT |
| P0DTC2 | Spike glycoprotein | 1206 | 505.7548 | 1009.4951 | 1009.4903 | 2 | 0.0048 | 19.32 | KLNDLCFT |

| | | | | | | | | | |
|---|---|---|---|---|---|---|---|---|---|
| P0DTC2 | Spike glycoprotein | 1206 | 505.7549 | 1009.4952 | 1009.4903 | 2 | 0.0049 | 30.34 | KLNDLCFT |
| P0DTC2 | Spike glycoprotein | 1206 | 505.7555 | 1009.4964 | 1009.4903 | 2 | 0.0061 | 19.43 | KLNDLCFT |
| P0DTC2 | Spike glycoprotein | 1206 | 506.2824 | 1010.5502 | 1010.5509 | 2 | -0.0007 | 10.06 | NLVRDLPQG |
| P0DTC2 | Spike glycoprotein | 1206 | 506.283 | 1010.5514 | 1010.5509 | 2 | 0.0005 | 15.29 | NLVRDLPQG |
| P0DTC2 | Spike glycoprotein | 1206 | 507.7651 | 1013.5156 | 1013.5142 | 2 | 0.0014 | 16.36 | DAVRDPQTL |
| P0DTC2 | Spike glycoprotein | 1206 | 507.7651 | 1013.5157 | 1013.5142 | 2 | 0.0015 | 29.04 | DAVRDPQTL |
| P0DTC2 | Spike glycoprotein | 1206 | 507.7657 | 1013.5168 | 1013.5142 | 2 | 0.0027 | 27.04 | DAVRDPQTL |
| P0DTC2 | Spike glycoprotein | 1206 | 507.7659 | 1013.5172 | 1013.5142 | 2 | 0.0031 | 29.06 | DAVRDPQTL |
| P0DTC2 | Spike glycoprotein | 1206 | 507.7659 | 1013.5173 | 1013.5142 | 2 | 0.0031 | 32.01 | DAVRDPQTL |
| P0DTC2 | Spike glycoprotein | 1206 | 507.766 | 1013.5174 | 1013.5142 | 2 | 0.0032 | 26.87 | DAVRDPQTL |
| P0DTC2 | Spike glycoprotein | 1206 | 507.766 | 1013.5174 | 1013.5142 | 2 | 0.0032 | 33.84 | DAVRDPQTL |
| P0DTC2 | Spike glycoprotein | 1206 | 507.7661 | 1013.5176 | 1013.5142 | 2 | 0.0034 | 31.75 | DAVRDPQTL |
| P0DTC2 | Spike glycoprotein | 1206 | 507.7661 | 1013.5176 | 1013.5142 | 2 | 0.0034 | 13.43 | DAVRDPQTL |
| P0DTC2 | Spike glycoprotein | 1206 | 507.7661 | 1013.5176 | 1013.5142 | 2 | 0.0034 | 28.94 | DAVRDPQTL |
| P0DTC2 | Spike glycoprotein | 1206 | 507.7661 | 1013.5177 | 1013.5142 | 2 | 0.0035 | 29 | DAVRDPQTL |
| P0DTC2 | Spike glycoprotein | 1206 | 507.7662 | 1013.5178 | 1013.5142 | 2 | 0.0037 | 26.84 | DAVRDPQTL |
| P0DTC2 | Spike glycoprotein | 1206 | 507.7662 | 1013.5179 | 1013.5142 | 2 | 0.0037 | 35.41 | DAVRDPQTL |
| P0DTC2 | Spike glycoprotein | 1206 | 507.7663 | 1013.518 | 1013.5142 | 2 | 0.0038 | 28.89 | DAVRDPQTL |
| P0DTC2 | Spike glycoprotein | 1206 | 507.7663 | 1013.5181 | 1013.5142 | 2 | 0.0039 | 17.48 | DAVRDPQTL |
| P0DTC2 | Spike glycoprotein | 1206 | 507.7665 | 1013.5183 | 1013.5142 | 2 | 0.0042 | 31.29 | DAVRDPQTL |
| P0DTC2 | Spike glycoprotein | 1206 | 507.7666 | 1013.5186 | 1013.5142 | 2 | 0.0044 | 21.37 | DAVRDPQTL |

| | | | | | | | | | |
|---|---|---|---|---|---|---|---|---|---|
| P0DTC2 | Spike glycoprotein | 1206 | 507.7669 | 1013.5193 | 1013.5142 | 2 | 0.0051 | 12.62 | DAVRDPQTL |
| P0DTC2 | Spike glycoprotein | 1206 | 507.7671 | 1013.5196 | 1013.5142 | 2 | 0.0055 | 8 | DAVRDPQTL |
| P0DTC2 | Spike glycoprotein | 1206 | 509.2659 | 1016.5172 | 1016.5138 | 2 | 0.0033 | 8.53 | IAVEQDKNT |
| P0DTC2 | Spike glycoprotein | 1206 | 511.2706 | 1020.5266 | 1020.524 | 2 | 0.0026 | 2.85 | SFVIRGDEV |
| P0DTC2 | Spike glycoprotein | 1206 | 514.2595 | 1026.5045 | 1026.5022 | 2 | 0.0023 | 2.18 | NVYADSFVI |
| P0DTC2 | Spike glycoprotein | 1206 | 514.2597 | 1026.5048 | 1026.5022 | 2 | 0.0026 | 19.99 | NVYADSFVI |
| P0DTC2 | Spike glycoprotein | 1206 | 514.2597 | 1026.5049 | 1026.5022 | 2 | 0.0026 | 23.78 | NVYADSFVI |
| P0DTC2 | Spike glycoprotein | 1206 | 514.2598 | 1026.505 | 1026.5022 | 2 | 0.0028 | 2.61 | NVYADSFVI |
| P0DTC2 | Spike glycoprotein | 1206 | 514.2602 | 1026.5058 | 1026.5022 | 2 | 0.0036 | 9.47 | NVYADSFVI |
| P0DTC2 | Spike glycoprotein | 1206 | 514.2613 | 1026.508 | 1026.5022 | 2 | 0.0058 | 9.83 | NVYADSFVI |
| P0DTC2 | Spike glycoprotein | 1206 | 514.7941 | 1027.5737 | 1027.5662 | 2 | 0.0075 | 24.02 | VQPTESIVR |
| P0DTC2 | Spike glycoprotein | 1206 | 514.8042 | 1027.5938 | 1027.5913 | 2 | 0.0025 | 3.86 | LIDLQELGK |
| P0DTC2 | Spike glycoprotein | 1206 | 517.7683 | 1033.5221 | 1033.5226 | 2 | -0.0006 | 8.28 | TVCGPKKSTN |
| P0DTC2 | Spike glycoprotein | 1206 | 522.285 | 1042.5554 | 1042.5519 | 2 | 0.0034 | 19.76 | RGDEVRQIA |
| P0DTC2 | Spike glycoprotein | 1206 | 522.285 | 1042.5554 | 1042.5519 | 2 | 0.0035 | 10.41 | RGDEVRQIA |
| P0DTC2 | Spike glycoprotein | 1206 | 522.2852 | 1042.5558 | 1042.5519 | 2 | 0.0038 | 11.37 | RGDEVRQIA |
| P0DTC2 | Spike glycoprotein | 1206 | 522.2861 | 1042.5577 | 1042.5519 | 2 | 0.0058 | 14.27 | RGDEVRQIA |
| P0DTC2 | Spike glycoprotein | 1206 | 523.7866 | 1045.5587 | 1045.5556 | 2 | 0.0031 | 21.19 | RTQLPPAYT |
| P0DTC2 | Spike glycoprotein | 1206 | 523.7867 | 1045.5588 | 1045.5556 | 2 | 0.0031 | 10.82 | RTQLPPAYT |
| P0DTC2 | Spike glycoprotein | 1206 | 523.7868 | 1045.559 | 1045.5556 | 2 | 0.0034 | 32.9 | RTQLPPAYT |
| P0DTC2 | Spike glycoprotein | 1206 | 523.7871 | 1045.5597 | 1045.5556 | 2 | 0.0041 | 33.12 | RTQLPPAYT |

| | | | | | | | | | |
|---|---|---|---|---|---|---|---|---|---|
| P0DTC2 | Spike glycoprotein | 1206 | 523.7872 | 1045.5598 | 1045.5556 | 2 | 0.0041 | 29.61 | RTQLPPAYT |
| P0DTC2 | Spike glycoprotein | 1206 | 523.7873 | 1045.56 | 1045.5556 | 2 | 0.0043 | 33.12 | RTQLPPAYT |
| P0DTC2 | Spike glycoprotein | 1206 | 523.7873 | 1045.56 | 1045.5556 | 2 | 0.0043 | 16.43 | RTQLPPAYT |
| P0DTC2 | Spike glycoprotein | 1206 | 523.7874 | 1045.5602 | 1045.5556 | 2 | 0.0046 | 31.38 | RTQLPPAYT |
| P0DTC2 | Spike glycoprotein | 1206 | 523.7877 | 1045.5608 | 1045.5556 | 2 | 0.0051 | 29.36 | RTQLPPAYT |
| P0DTC2 | Spike glycoprotein | 1206 | 523.7879 | 1045.5612 | 1045.5556 | 2 | 0.0056 | 17.27 | RTQLPPAYT |
| P0DTC2 | Spike glycoprotein | 1206 | 529.7509 | 1057.4873 | 1057.4903 | 2 | -0.003 | 5.24 | FKCYGVSPT |
| P0DTC2 | Spike glycoprotein | 1206 | 529.7528 | 1057.491 | 1057.4903 | 2 | 0.0007 | 2.48 | FKCYGVSPT |
| P0DTC2 | Spike glycoprotein | 1206 | 529.753 | 1057.4914 | 1057.4903 | 2 | 0.0011 | 6.99 | FKCYGVSPT |
| P0DTC2 | Spike glycoprotein | 1206 | 529.7533 | 1057.492 | 1057.4903 | 2 | 0.0017 | 15.89 | FKCYGVSPT |
| P0DTC2 | Spike glycoprotein | 1206 | 529.7535 | 1057.4925 | 1057.4903 | 2 | 0.0022 | 8.84 | FKCYGVSPT |
| P0DTC2 | Spike glycoprotein | 1206 | 529.7537 | 1057.4929 | 1057.4903 | 2 | 0.0026 | 19.21 | FKCYGVSPT |
| P0DTC2 | Spike glycoprotein | 1206 | 529.754 | 1057.4934 | 1057.4903 | 2 | 0.0032 | 15.42 | FKCYGVSPT |
| P0DTC2 | Spike glycoprotein | 1206 | 529.7544 | 1057.4943 | 1057.4903 | 2 | 0.0041 | 17.37 | FKCYGVSPT |
| P0DTC2 | Spike glycoprotein | 1206 | 529.7545 | 1057.4943 | 1057.4903 | 2 | 0.0041 | 6.28 | FKCYGVSPT |
| P0DTC2 | Spike glycoprotein | 1206 | 529.7545 | 1057.4945 | 1057.4903 | 2 | 0.0042 | 12 | FKCYGVSPT |
| P0DTC2 | Spike glycoprotein | 1206 | 529.7545 | 1057.4945 | 1057.4903 | 2 | 0.0042 | 12.85 | FKCYGVSPT |
| P0DTC2 | Spike glycoprotein | 1206 | 529.7546 | 1057.4946 | 1057.4903 | 2 | 0.0043 | 20.35 | FKCYGVSPT |
| P0DTC2 | Spike glycoprotein | 1206 | 529.7546 | 1057.4946 | 1057.4903 | 2 | 0.0043 | 5.8 | FKCYGVSPT |
| P0DTC2 | Spike glycoprotein | 1206 | 529.7548 | 1057.495 | 1057.4903 | 2 | 0.0047 | 22.41 | FKCYGVSPT |
| P0DTC2 | Spike glycoprotein | 1206 | 529.7549 | 1057.4952 | 1057.4903 | 2 | 0.0049 | 22.38 | FKCYGVSPT |

| | | | | | | | | | |
|---|---|---|---|---|---|---|---|---|---|
| P0DTC2 | Spike glycoprotein | 1206 | 529.7549 | 1057.4953 | 1057.4903 | 2 | 0.005 | 23.59 | FKCYGVSPT |
| P0DTC2 | Spike glycoprotein | 1206 | 533.7789 | 1065.5432 | 1065.5455 | 2 | -0.0023 | 27.09 | AIHADQLTPT |
| P0DTC2 | Spike glycoprotein | 1206 | 533.7797 | 1065.5449 | 1065.5455 | 2 | -0.0005 | 30.02 | AIHADQLTPT |
| P0DTC2 | Spike glycoprotein | 1206 | 533.78 | 1065.5454 | 1065.5455 | 2 | 0 | 33.97 | AIHADQLTPT |
| P0DTC2 | Spike glycoprotein | 1206 | 533.78 | 1065.5455 | 1065.5455 | 2 | 0.0001 | 25.86 | AIHADQLTPT |
| P0DTC2 | Spike glycoprotein | 1206 | 533.7804 | 1065.5462 | 1065.5455 | 2 | 0.0008 | 8.68 | AIHADQLTPT |
| P0DTC2 | Spike glycoprotein | 1206 | 533.7806 | 1065.5466 | 1065.5455 | 2 | 0.0011 | 33.57 | AIHADQLTPT |
| P0DTC2 | Spike glycoprotein | 1206 | 533.7807 | 1065.5469 | 1065.5455 | 2 | 0.0014 | 26.04 | AIHADQLTPT |
| P0DTC2 | Spike glycoprotein | 1206 | 533.7807 | 1065.5469 | 1065.5455 | 2 | 0.0015 | 9.65 | AIHADQLTPT |
| P0DTC2 | Spike glycoprotein | 1206 | 533.7809 | 1065.5472 | 1065.5455 | 2 | 0.0017 | 30.46 | AIHADQLTPT |
| P0DTC2 | Spike glycoprotein | 1206 | 533.7809 | 1065.5472 | 1065.5455 | 2 | 0.0017 | 30.32 | AIHADQLTPT |
| P0DTC2 | Spike glycoprotein | 1206 | 533.7809 | 1065.5473 | 1065.5455 | 2 | 0.0018 | 26.16 | AIHADQLTPT |
| P0DTC2 | Spike glycoprotein | 1206 | 533.7809 | 1065.5473 | 1065.5455 | 2 | 0.0019 | 3.52 | AIHADQLTPT |
| P0DTC2 | Spike glycoprotein | 1206 | 533.7812 | 1065.5478 | 1065.5455 | 2 | 0.0024 | 29.46 | AIHADQLTPT |
| P0DTC2 | Spike glycoprotein | 1206 | 533.7812 | 1065.5479 | 1065.5455 | 2 | 0.0024 | 32.67 | AIHADQLTPT |
| P0DTC2 | Spike glycoprotein | 1206 | 533.7812 | 1065.5479 | 1065.5455 | 2 | 0.0025 | 27.08 | AIHADQLTPT |
| P0DTC2 | Spike glycoprotein | 1206 | 533.7813 | 1065.548 | 1065.5455 | 2 | 0.0025 | 33.94 | AIHADQLTPT |
| P0DTC2 | Spike glycoprotein | 1206 | 533.7813 | 1065.548 | 1065.5455 | 2 | 0.0026 | 35.9 | AIHADQLTPT |
| P0DTC2 | Spike glycoprotein | 1206 | 533.7813 | 1065.548 | 1065.5455 | 2 | 0.0026 | 33.77 | AIHADQLTPT |
| P0DTC2 | Spike glycoprotein | 1206 | 533.7814 | 1065.5482 | 1065.5455 | 2 | 0.0027 | 7 | AIHADQLTPT |
| P0DTC2 | Spike glycoprotein | 1206 | 533.7814 | 1065.5482 | 1065.5455 | 2 | 0.0027 | 30.74 | AIHADQLTPT |

| | | | | | | | | | |
|---|---|---|---|---|---|---|---|---|---|
| P0DTC2 | Spike glycoprotein | 1206 | 533.7814 | 1065.5482 | 1065.5455 | 2 | 0.0028 | 19.4 | AIHADQLTPT |
| P0DTC2 | Spike glycoprotein | 1206 | 533.7814 | 1065.5482 | 1065.5455 | 2 | 0.0028 | 29.55 | AIHADQLTPT |
| P0DTC2 | Spike glycoprotein | 1206 | 533.7814 | 1065.5483 | 1065.5455 | 2 | 0.0028 | 26.75 | AIHADQLTPT |
| P0DTC2 | Spike glycoprotein | 1206 | 533.7814 | 1065.5483 | 1065.5455 | 2 | 0.0029 | 30.52 | AIHADQLTPT |
| P0DTC2 | Spike glycoprotein | 1206 | 533.7815 | 1065.5484 | 1065.5455 | 2 | 0.0029 | 30.9 | AIHADQLTPT |
| P0DTC2 | Spike glycoprotein | 1206 | 533.7815 | 1065.5484 | 1065.5455 | 2 | 0.0029 | 30.3 | AIHADQLTPT |
| P0DTC2 | Spike glycoprotein | 1206 | 533.7815 | 1065.5484 | 1065.5455 | 2 | 0.003 | 30.32 | AIHADQLTPT |
| P0DTC2 | Spike glycoprotein | 1206 | 533.7815 | 1065.5485 | 1065.5455 | 2 | 0.003 | 27.36 | AIHADQLTPT |
| P0DTC2 | Spike glycoprotein | 1206 | 533.7816 | 1065.5486 | 1065.5455 | 2 | 0.0031 | 32.89 | AIHADQLTPT |
| P0DTC2 | Spike glycoprotein | 1206 | 533.7816 | 1065.5487 | 1065.5455 | 2 | 0.0032 | 33.99 | AIHADQLTPT |
| P0DTC2 | Spike glycoprotein | 1206 | 533.7817 | 1065.5488 | 1065.5455 | 2 | 0.0034 | 33.91 | AIHADQLTPT |
| P0DTC2 | Spike glycoprotein | 1206 | 533.7817 | 1065.5489 | 1065.5455 | 2 | 0.0034 | 28.7 | AIHADQLTPT |
| P0DTC2 | Spike glycoprotein | 1206 | 533.7818 | 1065.549 | 1065.5455 | 2 | 0.0035 | 30.26 | AIHADQLTPT |
| P0DTC2 | Spike glycoprotein | 1206 | 533.7818 | 1065.5491 | 1065.5455 | 2 | 0.0036 | 32.45 | AIHADQLTPT |
| P0DTC2 | Spike glycoprotein | 1206 | 533.7818 | 1065.5491 | 1065.5455 | 2 | 0.0036 | 34.03 | AIHADQLTPT |
| P0DTC2 | Spike glycoprotein | 1206 | 533.7819 | 1065.5492 | 1065.5455 | 2 | 0.0037 | 22.96 | AIHADQLTPT |
| P0DTC2 | Spike glycoprotein | 1206 | 533.7819 | 1065.5492 | 1065.5455 | 2 | 0.0038 | 31.8 | AIHADQLTPT |
| P0DTC2 | Spike glycoprotein | 1206 | 533.7819 | 1065.5493 | 1065.5455 | 2 | 0.0038 | 33.76 | AIHADQLTPT |
| P0DTC2 | Spike glycoprotein | 1206 | 533.782 | 1065.5494 | 1065.5455 | 2 | 0.0039 | 34.04 | AIHADQLTPT |
| P0DTC2 | Spike glycoprotein | 1206 | 533.782 | 1065.5495 | 1065.5455 | 2 | 0.0041 | 32.63 | AIHADQLTPT |
| P0DTC2 | Spike glycoprotein | 1206 | 533.7821 | 1065.5496 | 1065.5455 | 2 | 0.0041 | 33.98 | AIHADQLTPT |

| | | | | | | | | | |
|---|---|---|---|---|---|---|---|---|---|
| P0DTC2 | Spike glycoprotein | 1206 | 533.7822 | 1065.5499 | 1065.5455 | 2 | 0.0044 | 2.85 | AIHADQLTPT |
| P0DTC2 | Spike glycoprotein | 1206 | 533.7823 | 1065.55 | 1065.5455 | 2 | 0.0045 | 32.47 | AIHADQLTPT |
| P0DTC2 | Spike glycoprotein | 1206 | 533.7823 | 1065.5501 | 1065.5455 | 2 | 0.0047 | 6.76 | AIHADQLTPT |
| P0DTC2 | Spike glycoprotein | 1206 | 533.7828 | 1065.551 | 1065.5455 | 2 | 0.0055 | 12.34 | AIHADQLTPT |
| P0DTC2 | Spike glycoprotein | 1206 | 533.7829 | 1065.5513 | 1065.5455 | 2 | 0.0058 | 32.4 | AIHADQLTPT |
| P0DTC2 | Spike glycoprotein | 1206 | 533.7838 | 1065.5531 | 1065.5455 | 2 | 0.0077 | 25.12 | AIHADQLTPT |
| P0DTC2 | Spike glycoprotein | 1206 | 533.7839 | 1065.5532 | 1065.5455 | 2 | 0.0077 | 29.35 | AIHADQLTPT |
| P0DTC2 | Spike glycoprotein | 1206 | 537.7762 | 1073.5378 | 1073.5353 | 2 | 0.0025 | 24.58 | GIAVEQDKNT |
| P0DTC2 | Spike glycoprotein | 1206 | 538.3004 | 1074.5863 | 1074.5856 | 2 | 0.0007 | 1.68 | TNLVKNKCV |
| P0DTC2 | Spike glycoprotein | 1206 | 538.3092 | 1074.6038 | 1074.5974 | 2 | 0.0063 | 6.19 | NIIRGWIFG |
| P0DTC2 | Spike glycoprotein | 1206 | 540.323 | 1078.6315 | 1078.6274 | 2 | 0.0041 | 7.27 | ALEPLVDLPI |
| P0DTC2 | Spike glycoprotein | 1206 | 541.2998 | 1080.585 | 1080.5815 | 2 | 0.0035 | 4.63 | QILPDPSKPS |
| P0DTC2 | Spike glycoprotein | 1206 | 541.2999 | 1080.5852 | 1080.5815 | 2 | 0.0037 | 17.18 | QILPDPSKPS |
| P0DTC2 | Spike glycoprotein | 1206 | 542.2971 | 1082.5796 | 1082.5794 | 2 | 0.0002 | 29.75 | KCTLKSFTV |
| P0DTC2 | Spike glycoprotein | 1206 | 542.2982 | 1082.5818 | 1082.5794 | 2 | 0.0024 | 31.18 | KCTLKSFTV |
| P0DTC2 | Spike glycoprotein | 1206 | 542.2987 | 1082.5829 | 1082.5794 | 2 | 0.0035 | 26.39 | KCTLKSFTV |
| P0DTC2 | Spike glycoprotein | 1206 | 542.2988 | 1082.5831 | 1082.5794 | 2 | 0.0036 | 7.69 | KCTLKSFTV |
| P0DTC2 | Spike glycoprotein | 1206 | 542.8045 | 1083.5945 | 1083.5924 | 2 | 0.0021 | 43.23 | RDPQTLEIL |
| P0DTC2 | Spike glycoprotein | 1206 | 542.8047 | 1083.5949 | 1083.5924 | 2 | 0.0025 | 13.42 | RDPQTLEIL |
| P0DTC2 | Spike glycoprotein | 1206 | 542.8047 | 1083.5949 | 1083.5924 | 2 | 0.0025 | 15.64 | RDPQTLEIL |
| P0DTC2 | Spike glycoprotein | 1206 | 542.8048 | 1083.5951 | 1083.5924 | 2 | 0.0027 | 44.02 | RDPQTLEIL |

| | | | | | | | | | |
|---|---|---|---|---|---|---|---|---|---|
| P0DTC2 | Spike glycoprotein | 1206 | 542.8049 | 1083.5952 | 1083.5924 | 2 | 0.0027 | 27.21 | RDPQTLEIL |
| P0DTC2 | Spike glycoprotein | 1206 | 542.8049 | 1083.5952 | 1083.5924 | 2 | 0.0028 | 49.33 | RDPQTLEIL |
| P0DTC2 | Spike glycoprotein | 1206 | 542.805 | 1083.5954 | 1083.5924 | 2 | 0.003 | 62.19 | RDPQTLEIL |
| P0DTC2 | Spike glycoprotein | 1206 | 542.8052 | 1083.5959 | 1083.5924 | 2 | 0.0035 | 42.56 | RDPQTLEIL |
| P0DTC2 | Spike glycoprotein | 1206 | 542.8081 | 1083.6016 | 1083.5924 | 2 | 0.0092 | 1.6 | RDPQTLEIL |
| P0DTC2 | Spike glycoprotein | 1206 | 544.7997 | 1087.5849 | 1087.5808 | 2 | 0.0041 | 16.37 | NLVKNKCVN |
| P0DTC2 | Spike glycoprotein | 1206 | 546.2807 | 1090.5468 | 1090.5441 | 2 | 0.0027 | 20.21 | TVCGPKKSTN |
| P0DTC2 | Spike glycoprotein | 1206 | 548.7936 | 1095.5726 | 1095.5713 | 2 | 0.0013 | 31.92 | RGVYYPDKV |
| P0DTC2 | Spike glycoprotein | 1206 | 548.7938 | 1095.5731 | 1095.5713 | 2 | 0.0019 | 6.26 | RGVYYPDKV |
| P0DTC2 | Spike glycoprotein | 1206 | 548.794 | 1095.5735 | 1095.5713 | 2 | 0.0022 | 20.16 | RGVYYPDKV |
| P0DTC2 | Spike glycoprotein | 1206 | 548.7943 | 1095.5741 | 1095.5713 | 2 | 0.0028 | 20.05 | RGVYYPDKV |
| P0DTC2 | Spike glycoprotein | 1206 | 548.7946 | 1095.5747 | 1095.5713 | 2 | 0.0034 | 26.58 | RGVYYPDKV |
| P0DTC2 | Spike glycoprotein | 1206 | 548.7947 | 1095.5749 | 1095.5713 | 2 | 0.0036 | 29.65 | RGVYYPDKV |
| P0DTC2 | Spike glycoprotein | 1206 | 548.7948 | 1095.575 | 1095.5713 | 2 | 0.0037 | 25.22 | RGVYYPDKV |
| P0DTC2 | Spike glycoprotein | 1206 | 548.7948 | 1095.5751 | 1095.5713 | 2 | 0.0038 | 24.94 | RGVYYPDKV |
| P0DTC2 | Spike glycoprotein | 1206 | 552.8076 | 1103.6007 | 1103.5975 | 2 | 0.0032 | 0.04 | LKPFERDIS |
| P0DTC2 | Spike glycoprotein | 1206 | 560.2665 | 1118.5184 | 1118.5145 | 2 | 0.0039 | 30.11 | GGNYNYLYR |
| P0DTC2 | Spike glycoprotein | 1206 | 560.2666 | 1118.5186 | 1118.5145 | 2 | 0.0041 | 30.06 | GGNYNYLYR |
| P0DTC2 | Spike glycoprotein | 1206 | 560.2667 | 1118.5188 | 1118.5145 | 2 | 0.0043 | 46.07 | GGNYNYLYR |
| P0DTC2 | Spike glycoprotein | 1206 | 560.7457 | 1119.4768 | 1119.4754 | 2 | 0.0014 | 10.71 | DCALDPLSET |
| P0DTC2 | Spike glycoprotein | 1206 | 560.7459 | 1119.4773 | 1119.4754 | 2 | 0.0019 | 9.08 | DCALDPLSET |

| | | | | | | | | | |
|---|---|---|---|---|---|---|---|---|---|
| P0DTC2 | Spike glycoprotein | 1206 | 560.7464 | 1119.4783 | 1119.4754 | 2 | 0.0029 | 7.55 | DCALDPLSET |
| P0DTC2 | Spike glycoprotein | 1206 | 560.7465 | 1119.4785 | 1119.4754 | 2 | 0.0031 | 57.19 | DCALDPLSET |
| P0DTC2 | Spike glycoprotein | 1206 | 560.7466 | 1119.4786 | 1119.4754 | 2 | 0.0031 | 35.8 | DCALDPLSET |
| P0DTC2 | Spike glycoprotein | 1206 | 560.7468 | 1119.4791 | 1119.4754 | 2 | 0.0037 | 21.39 | DCALDPLSET |
| P0DTC2 | Spike glycoprotein | 1206 | 560.7469 | 1119.4792 | 1119.4754 | 2 | 0.0038 | 36.91 | DCALDPLSET |
| P0DTC2 | Spike glycoprotein | 1206 | 560.7473 | 1119.48 | 1119.4754 | 2 | 0.0046 | 11.83 | DCALDPLSET |
| P0DTC2 | Spike glycoprotein | 1206 | 560.7481 | 1119.4817 | 1119.4754 | 2 | 0.0063 | 32.45 | DCALDPLSET |
| P0DTC2 | Spike glycoprotein | 1206 | 562.775 | 1123.5355 | 1123.5332 | 2 | 0.0023 | 13.74 | KLNDLCFTN |
| P0DTC2 | Spike glycoprotein | 1206 | 563.2896 | 1124.5646 | 1124.5615 | 2 | 0.0031 | 9.2 | SFPQSAPHGVV |
| P0DTC2 | Spike glycoprotein | 1206 | 563.2905 | 1124.5664 | 1124.5615 | 2 | 0.005 | 9.46 | SFPQSAPHGVV |
| P0DTC2 | Spike glycoprotein | 1206 | 568.292 | 1134.5694 | 1134.5669 | 2 | 0.0025 | 13.57 | DLEGKQGNFK |
| P0DTC2 | Spike glycoprotein | 1206 | 379.1975 | 1134.5706 | 1134.5669 | 3 | 0.0036 | 32.51 | DLEGKQGNFK |
| P0DTC2 | Spike glycoprotein | 1206 | 569.3096 | 1136.6046 | 1136.5979 | 2 | 0.0068 | 10.24 | GVGYQPYRVV |
| P0DTC2 | Spike glycoprotein | 1206 | 570.3048 | 1138.595 | 1138.5924 | 2 | 0.0027 | 13.52 | FLPFQQFGR |
| P0DTC2 | Spike glycoprotein | 1206 | 570.3052 | 1138.5959 | 1138.5924 | 2 | 0.0035 | 1.51 | FLPFQQFGR |
| P0DTC2 | Spike glycoprotein | 1206 | 570.3053 | 1138.5961 | 1138.5924 | 2 | 0.0037 | 11.06 | FLPFQQFGR |
| P0DTC2 | Spike glycoprotein | 1206 | 570.3055 | 1138.5965 | 1138.5924 | 2 | 0.0041 | 16.33 | FLPFQQFGR |
| P0DTC2 | Spike glycoprotein | 1206 | 574.3064 | 1146.5982 | 1146.6033 | 2 | -0.0051 | 16.39 | TRTQLPPAYT |
| P0DTC2 | Spike glycoprotein | 1206 | 574.3068 | 1146.599 | 1146.6033 | 2 | -0.0043 | 16.04 | TRTQLPPAYT |
| P0DTC2 | Spike glycoprotein | 1206 | 574.3073 | 1146.6001 | 1146.6033 | 2 | -0.0032 | 26.07 | TRTQLPPAYT |
| P0DTC2 | Spike glycoprotein | 1206 | 574.3076 | 1146.6006 | 1146.6033 | 2 | -0.0027 | 17.37 | TRTQLPPAYT |

| | | | | | | | | | |
|---|---|---|---|---|---|---|---|---|---|
| P0DTC2 | Spike glycoprotein | 1206 | 574.3082 | 1146.6019 | 1146.6033 | 2 | -0.0014 | 25.76 | TRTQLPPAYT |
| P0DTC2 | Spike glycoprotein | 1206 | 574.3084 | 1146.6023 | 1146.6033 | 2 | -0.001 | 21.62 | TRTQLPPAYT |
| P0DTC2 | Spike glycoprotein | 1206 | 574.3088 | 1146.6031 | 1146.6033 | 2 | -0.0003 | 14.19 | TRTQLPPAYT |
| P0DTC2 | Spike glycoprotein | 1206 | 574.3093 | 1146.6041 | 1146.6033 | 2 | 0.0008 | 28.42 | TRTQLPPAYT |
| P0DTC2 | Spike glycoprotein | 1206 | 574.3094 | 1146.6042 | 1146.6033 | 2 | 0.0009 | 14.05 | TRTQLPPAYT |
| P0DTC2 | Spike glycoprotein | 1206 | 574.3094 | 1146.6042 | 1146.6033 | 2 | 0.0009 | 21.15 | TRTQLPPAYT |
| P0DTC2 | Spike glycoprotein | 1206 | 574.3094 | 1146.6042 | 1146.6033 | 2 | 0.0009 | 39.17 | TRTQLPPAYT |
| P0DTC2 | Spike glycoprotein | 1206 | 574.3094 | 1146.6043 | 1146.6033 | 2 | 0.001 | 16.53 | TRTQLPPAYT |
| P0DTC2 | Spike glycoprotein | 1206 | 574.3094 | 1146.6043 | 1146.6033 | 2 | 0.001 | 24.44 | TRTQLPPAYT |
| P0DTC2 | Spike glycoprotein | 1206 | 574.3094 | 1146.6043 | 1146.6033 | 2 | 0.001 | 25.42 | TRTQLPPAYT |
| P0DTC2 | Spike glycoprotein | 1206 | 574.3095 | 1146.6045 | 1146.6033 | 2 | 0.0011 | 13.51 | TRTQLPPAYT |
| P0DTC2 | Spike glycoprotein | 1206 | 574.3097 | 1146.6048 | 1146.6033 | 2 | 0.0015 | 27.77 | TRTQLPPAYT |
| P0DTC2 | Spike glycoprotein | 1206 | 574.3097 | 1146.6048 | 1146.6033 | 2 | 0.0015 | 37.72 | TRTQLPPAYT |
| P0DTC2 | Spike glycoprotein | 1206 | 574.3099 | 1146.6053 | 1146.6033 | 2 | 0.002 | 22.66 | TRTQLPPAYT |
| P0DTC2 | Spike glycoprotein | 1206 | 574.31 | 1146.6054 | 1146.6033 | 2 | 0.0021 | 38.72 | TRTQLPPAYT |
| P0DTC2 | Spike glycoprotein | 1206 | 574.31 | 1146.6055 | 1146.6033 | 2 | 0.0021 | 22.53 | TRTQLPPAYT |
| P0DTC2 | Spike glycoprotein | 1206 | 574.3101 | 1146.6056 | 1146.6033 | 2 | 0.0023 | 39.11 | TRTQLPPAYT |
| P0DTC2 | Spike glycoprotein | 1206 | 574.3102 | 1146.6058 | 1146.6033 | 2 | 0.0025 | 28.07 | TRTQLPPAYT |
| P0DTC2 | Spike glycoprotein | 1206 | 574.3102 | 1146.6059 | 1146.6033 | 2 | 0.0026 | 18.14 | TRTQLPPAYT |
| P0DTC2 | Spike glycoprotein | 1206 | 574.3102 | 1146.6059 | 1146.6033 | 2 | 0.0026 | 16.77 | TRTQLPPAYT |
| P0DTC2 | Spike glycoprotein | 1206 | 574.3104 | 1146.6062 | 1146.6033 | 2 | 0.0029 | 39.01 | TRTQLPPAYT |

| | | | | | | | | | |
|---|---|---|---|---|---|---|---|---|---|
| P0DTC2 | Spike glycoprotein | 1206 | 574.3104 | 1146.6062 | 1146.6033 | 2 | 0.0029 | 42.52 | TRTQLPPAYT |
| P0DTC2 | Spike glycoprotein | 1206 | 574.3105 | 1146.6065 | 1146.6033 | 2 | 0.0031 | 22.14 | TRTQLPPAYT |
| P0DTC2 | Spike glycoprotein | 1206 | 574.3106 | 1146.6067 | 1146.6033 | 2 | 0.0033 | 26.59 | TRTQLPPAYT |
| P0DTC2 | Spike glycoprotein | 1206 | 574.3107 | 1146.6069 | 1146.6033 | 2 | 0.0036 | 27.63 | TRTQLPPAYT |
| P0DTC2 | Spike glycoprotein | 1206 | 574.3113 | 1146.608 | 1146.6033 | 2 | 0.0047 | 14.73 | TRTQLPPAYT |
| P0DTC2 | Spike glycoprotein | 1206 | 574.3113 | 1146.6081 | 1146.6033 | 2 | 0.0048 | 31.81 | TRTQLPPAYT |
| P0DTC2 | Spike glycoprotein | 1206 | 575.8354 | 1149.6562 | 1149.6546 | 2 | 0.0016 | 15.87 | YLQPRTFLL |
| P0DTC2 | Spike glycoprotein | 1206 | 575.8354 | 1149.6563 | 1149.6546 | 2 | 0.0017 | 17.58 | YLQPRTFLL |
| P0DTC2 | Spike glycoprotein | 1206 | 575.8355 | 1149.6564 | 1149.6546 | 2 | 0.0018 | 23.45 | YLQPRTFLL |
| P0DTC2 | Spike glycoprotein | 1206 | 575.8355 | 1149.6565 | 1149.6546 | 2 | 0.0019 | 19.69 | YLQPRTFLL |
| P0DTC2 | Spike glycoprotein | 1206 | 575.836 | 1149.6574 | 1149.6546 | 2 | 0.0028 | 14.38 | YLQPRTFLL |
| P0DTC2 | Spike glycoprotein | 1206 | 575.836 | 1149.6575 | 1149.6546 | 2 | 0.0029 | 12.92 | YLQPRTFLL |
| P0DTC2 | Spike glycoprotein | 1206 | 575.8362 | 1149.6578 | 1149.6546 | 2 | 0.0031 | 23.53 | YLQPRTFLL |
| P0DTC2 | Spike glycoprotein | 1206 | 575.8363 | 1149.658 | 1149.6546 | 2 | 0.0034 | 16.22 | YLQPRTFLL |
| P0DTC2 | Spike glycoprotein | 1206 | 575.8363 | 1149.658 | 1149.6546 | 2 | 0.0034 | 22.13 | YLQPRTFLL |
| P0DTC2 | Spike glycoprotein | 1206 | 575.8363 | 1149.6581 | 1149.6546 | 2 | 0.0035 | 20.62 | YLQPRTFLL |
| P0DTC2 | Spike glycoprotein | 1206 | 575.8364 | 1149.6582 | 1149.6546 | 2 | 0.0036 | 17.93 | YLQPRTFLL |
| P0DTC2 | Spike glycoprotein | 1206 | 575.8364 | 1149.6583 | 1149.6546 | 2 | 0.0037 | 9.53 | YLQPRTFLL |
| P0DTC2 | Spike glycoprotein | 1206 | 575.8365 | 1149.6584 | 1149.6546 | 2 | 0.0037 | 6.46 | YLQPRTFLL |
| P0DTC2 | Spike glycoprotein | 1206 | 575.8365 | 1149.6584 | 1149.6546 | 2 | 0.0038 | 24.76 | YLQPRTFLL |
| P0DTC2 | Spike glycoprotein | 1206 | 575.8365 | 1149.6585 | 1149.6546 | 2 | 0.0039 | 15.97 | YLQPRTFLL |

| | | | | | | | | | |
|---|---|---|---|---|---|---|---|---|---|
| P0DTC2 | Spike glycoprotein | 1206 | 575.8365 | 1149.6585 | 1149.6546 | 2 | 0.0039 | 23.69 | YLQPRTFLL |
| P0DTC2 | Spike glycoprotein | 1206 | 575.8365 | 1149.6585 | 1149.6546 | 2 | 0.0039 | 19.85 | YLQPRTFLL |
| P0DTC2 | Spike glycoprotein | 1206 | 575.8368 | 1149.659 | 1149.6546 | 2 | 0.0044 | 19.72 | YLQPRTFLL |
| P0DTC2 | Spike glycoprotein | 1206 | 575.8371 | 1149.6596 | 1149.6546 | 2 | 0.005 | 10.85 | YLQPRTFLL |
| P0DTC2 | Spike glycoprotein | 1206 | 576.7944 | 1151.5743 | 1151.5723 | 2 | 0.0019 | 1.94 | VYYHKNNKS |
| P0DTC2 | Spike glycoprotein | 1206 | 580.2785 | 1158.5425 | 1158.538 | 2 | 0.0045 | 6.86 | TFKCYGVSPT |
| P0DTC2 | Spike glycoprotein | 1206 | 580.8082 | 1159.6019 | 1159.5986 | 2 | 0.0033 | 18.77 | RTQLPPAYTN |
| P0DTC2 | Spike glycoprotein | 1206 | 580.8086 | 1159.6027 | 1159.5986 | 2 | 0.0041 | 27.84 | RTQLPPAYTN |
| P0DTC2 | Spike glycoprotein | 1206 | 580.8108 | 1159.6071 | 1159.5986 | 2 | 0.0085 | 19.12 | RTQLPPAYTN |
| P0DTC2 | Spike glycoprotein | 1206 | 587.804 | 1173.5935 | 1173.5931 | 2 | 0.0004 | 20.21 | HFPREGVFVS |
| P0DTC2 | Spike glycoprotein | 1206 | 587.8042 | 1173.5938 | 1173.5931 | 2 | 0.0007 | 21.97 | HFPREGVFVS |
| P0DTC2 | Spike glycoprotein | 1206 | 392.2054 | 1173.5943 | 1173.5931 | 3 | 0.0012 | 14.8 | HFPREGVFVS |
| P0DTC2 | Spike glycoprotein | 1206 | 392.206 | 1173.5961 | 1173.5931 | 3 | 0.003 | 13.3 | HFPREGVFVS |
| P0DTC2 | Spike glycoprotein | 1206 | 588.7867 | 1175.5589 | 1175.5571 | 2 | 0.0019 | 8.54 | WNSNNLDSKV |
| P0DTC2 | Spike glycoprotein | 1206 | 588.7869 | 1175.5593 | 1175.5571 | 2 | 0.0022 | 11.06 | WNSNNLDSKV |
| P0DTC2 | Spike glycoprotein | 1206 | 588.7873 | 1175.56 | 1175.5571 | 2 | 0.0029 | 7.2 | WNSNNLDSKV |
| P0DTC2 | Spike glycoprotein | 1206 | 588.7874 | 1175.5602 | 1175.5571 | 2 | 0.0031 | 15.25 | WNSNNLDSKV |
| P0DTC2 | Spike glycoprotein | 1206 | 588.7875 | 1175.5605 | 1175.5571 | 2 | 0.0034 | 63.03 | WNSNNLDSKV |
| P0DTC2 | Spike glycoprotein | 1206 | 588.7877 | 1175.5608 | 1175.5571 | 2 | 0.0037 | 45.4 | WNSNNLDSKV |
| P0DTC2 | Spike glycoprotein | 1206 | 588.7877 | 1175.5608 | 1175.5571 | 2 | 0.0037 | 10.21 | WNSNNLDSKV |
| P0DTC2 | Spike glycoprotein | 1206 | 588.7877 | 1175.5608 | 1175.5571 | 2 | 0.0037 | 49.92 | WNSNNLDSKV |

| | | | | | | | | | |
|---|---|---|---|---|---|---|---|---|---|
| P0DTC2 | Spike glycoprotein | 1206 | 588.788 | 1175.5614 | 1175.5571 | 2 | 0.0043 | 5.71 | WNSNNLDSKV |
| P0DTC2 | Spike glycoprotein | 1206 | 588.7885 | 1175.5625 | 1175.5571 | 2 | 0.0054 | 10.94 | WNSNNLDSKV |
| P0DTC2 | Spike glycoprotein | 1206 | 588.7886 | 1175.5626 | 1175.5571 | 2 | 0.0056 | 22.08 | WNSNNLDSKV |
| P0DTC2 | Spike glycoprotein | 1206 | 588.7891 | 1175.5636 | 1175.5571 | 2 | 0.0066 | 23.2 | WNSNNLDSKV |
| P0DTC2 | Spike glycoprotein | 1206 | 588.7908 | 1175.567 | 1175.5571 | 2 | 0.0099 | 12.42 | WNSNNLDSKV |
| P0DTC2 | Spike glycoprotein | 1206 | 588.8301 | 1175.6456 | 1175.6451 | 2 | 0.0005 | 11.08 | NIIRGWIFGT |
| P0DTC2 | Spike glycoprotein | 1206 | 588.8305 | 1175.6464 | 1175.6451 | 2 | 0.0013 | 36.56 | NIIRGWIFGT |
| P0DTC2 | Spike glycoprotein | 1206 | 588.8309 | 1175.6473 | 1175.6451 | 2 | 0.0022 | 16.79 | NIIRGWIFGT |
| P0DTC2 | Spike glycoprotein | 1206 | 588.831 | 1175.6475 | 1175.6451 | 2 | 0.0024 | 17.67 | NIIRGWIFGT |
| P0DTC2 | Spike glycoprotein | 1206 | 588.8311 | 1175.6477 | 1175.6451 | 2 | 0.0026 | 32.47 | NIIRGWIFGT |
| P0DTC2 | Spike glycoprotein | 1206 | 588.8321 | 1175.6496 | 1175.6451 | 2 | 0.0045 | 33.2 | NIIRGWIFGT |
| P0DTC2 | Spike glycoprotein | 1206 | 588.8322 | 1175.6498 | 1175.6451 | 2 | 0.0046 | 11.62 | NIIRGWIFGT |
| P0DTC2 | Spike glycoprotein | 1206 | 599.3207 | 1196.6268 | 1196.6262 | 2 | 0.0006 | 2.57 | RGDEVRQIAPG |
| P0DTC2 | Spike glycoprotein | 1206 | 599.3213 | 1196.628 | 1196.6262 | 2 | 0.0019 | 3.07 | RGDEVRQIAPG |
| P0DTC2 | Spike glycoprotein | 1206 | 599.3215 | 1196.6285 | 1196.6262 | 2 | 0.0023 | 8.15 | RGDEVRQIAPG |
| P0DTC2 | Spike glycoprotein | 1206 | 599.3217 | 1196.6289 | 1196.6262 | 2 | 0.0028 | 30.54 | RGDEVRQIAPG |
| P0DTC2 | Spike glycoprotein | 1206 | 399.8837 | 1196.6293 | 1196.6262 | 3 | 0.0031 | 17.82 | RGDEVRQIAPG |
| P0DTC2 | Spike glycoprotein | 1206 | 599.3219 | 1196.6293 | 1196.6262 | 2 | 0.0031 | 21.19 | RGDEVRQIAPG |
| P0DTC2 | Spike glycoprotein | 1206 | 599.3224 | 1196.6303 | 1196.6262 | 2 | 0.0042 | 4.09 | RGDEVRQIAPG |
| P0DTC2 | Spike glycoprotein | 1206 | 599.3225 | 1196.6305 | 1196.6262 | 2 | 0.0043 | 17.74 | RGDEVRQIAPG |
| P0DTC2 | Spike glycoprotein | 1206 | 599.3226 | 1196.6307 | 1196.6262 | 2 | 0.0045 | 9.67 | RGDEVRQIAPG |

| | | | | | | | | | |
|---|---|---|---|---|---|---|---|---|---|
| P0DTC2 | Spike glycoprotein | 1206 | 399.8842 | 1196.6307 | 1196.6262 | 3 | 0.0045 | 19.36 | RGDEVRQIAPG |
| P0DTC2 | Spike glycoprotein | 1206 | 599.3226 | 1196.6307 | 1196.6262 | 2 | 0.0046 | 21.27 | RGDEVRQIAPG |
| P0DTC2 | Spike glycoprotein | 1206 | 599.3228 | 1196.631 | 1196.6262 | 2 | 0.0048 | 13.96 | RGDEVRQIAPG |
| P0DTC2 | Spike glycoprotein | 1206 | 599.3233 | 1196.632 | 1196.6262 | 2 | 0.0058 | 14.79 | RGDEVRQIAPG |
| P0DTC2 | Spike glycoprotein | 1206 | 601.8323 | 1201.65 | 1201.6455 | 2 | 0.0045 | 16.52 | LVRDLPQGFSA |
| P0DTC2 | Spike glycoprotein | 1206 | 601.8325 | 1201.6504 | 1201.6455 | 2 | 0.0048 | 16.91 | LVRDLPQGFSA |
| P0DTC2 | Spike glycoprotein | 1206 | 603.3318 | 1204.6491 | 1204.6452 | 2 | 0.0039 | 7.04 | LKPFERDIST |
| P0DTC2 | Spike glycoprotein | 1206 | 603.3325 | 1204.6505 | 1204.6452 | 2 | 0.0054 | 15.96 | LKPFERDIST |
| P0DTC2 | Spike glycoprotein | 1206 | 607.3146 | 1212.6146 | 1212.6099 | 2 | 0.0047 | 10.73 | DEVRQIAPGQT |
| P0DTC2 | Spike glycoprotein | 1206 | 609.8255 | 1217.6364 | 1217.6404 | 2 | -0.004 | 9.42 | NLKPFERDIS |
| P0DTC2 | Spike glycoprotein | 1206 | 609.8257 | 1217.6369 | 1217.6404 | 2 | -0.0035 | 11.45 | NLKPFERDIS |
| P0DTC2 | Spike glycoprotein | 1206 | 609.8265 | 1217.6384 | 1217.6404 | 2 | -0.002 | 9.06 | NLKPFERDIS |
| P0DTC2 | Spike glycoprotein | 1206 | 406.8874 | 1217.6403 | 1217.6404 | 3 | -0.0001 | 4.94 | NLKPFERDIS |
| P0DTC2 | Spike glycoprotein | 1206 | 609.8275 | 1217.6404 | 1217.6404 | 2 | 0 | 5 | NLKPFERDIS |
| P0DTC2 | Spike glycoprotein | 1206 | 406.8874 | 1217.6405 | 1217.6404 | 3 | 0.0001 | 6.4 | NLKPFERDIS |
| P0DTC2 | Spike glycoprotein | 1206 | 609.8277 | 1217.6408 | 1217.6404 | 2 | 0.0004 | 0.81 | NLKPFERDIS |
| P0DTC2 | Spike glycoprotein | 1206 | 406.8877 | 1217.6413 | 1217.6404 | 3 | 0.0008 | 10.48 | NLKPFERDIS |
| P0DTC2 | Spike glycoprotein | 1206 | 609.8279 | 1217.6413 | 1217.6404 | 2 | 0.0009 | 12.6 | NLKPFERDIS |
| P0DTC2 | Spike glycoprotein | 1206 | 406.8877 | 1217.6414 | 1217.6404 | 3 | 0.001 | 1.55 | NLKPFERDIS |
| P0DTC2 | Spike glycoprotein | 1206 | 406.8878 | 1217.6417 | 1217.6404 | 3 | 0.0013 | 4.95 | NLKPFERDIS |
| P0DTC2 | Spike glycoprotein | 1206 | 406.8879 | 1217.6418 | 1217.6404 | 3 | 0.0014 | 7.62 | NLKPFERDIS |

| | | | | | | | | | |
|---|---|---|---|---|---|---|---|---|---|
| P0DTC2 | Spike glycoprotein | 1206 | 406.888 | 1217.6421 | 1217.6404 | 3 | 0.0017 | 3.53 | NLKPFERDIS |
| P0DTC2 | Spike glycoprotein | 1206 | 406.888 | 1217.6421 | 1217.6404 | 3 | 0.0017 | 7.32 | NLKPFERDIS |
| P0DTC2 | Spike glycoprotein | 1206 | 406.888 | 1217.6421 | 1217.6404 | 3 | 0.0017 | 4.26 | NLKPFERDIS |
| P0DTC2 | Spike glycoprotein | 1206 | 609.8284 | 1217.6422 | 1217.6404 | 2 | 0.0018 | 0.2 | NLKPFERDIS |
| P0DTC2 | Spike glycoprotein | 1206 | 609.8285 | 1217.6424 | 1217.6404 | 2 | 0.0019 | 9.92 | NLKPFERDIS |
| P0DTC2 | Spike glycoprotein | 1206 | 406.8881 | 1217.6424 | 1217.6404 | 3 | 0.002 | 1.17 | NLKPFERDIS |
| P0DTC2 | Spike glycoprotein | 1206 | 406.8882 | 1217.6426 | 1217.6404 | 3 | 0.0022 | 1.64 | NLKPFERDIS |
| P0DTC2 | Spike glycoprotein | 1206 | 406.8883 | 1217.6431 | 1217.6404 | 3 | 0.0027 | 3.71 | NLKPFERDIS |
| P0DTC2 | Spike glycoprotein | 1206 | 406.8884 | 1217.6433 | 1217.6404 | 3 | 0.0029 | 5.97 | NLKPFERDIS |
| P0DTC2 | Spike glycoprotein | 1206 | 609.8289 | 1217.6433 | 1217.6404 | 2 | 0.0029 | 10.81 | NLKPFERDIS |
| P0DTC2 | Spike glycoprotein | 1206 | 609.8291 | 1217.6436 | 1217.6404 | 2 | 0.0032 | 9.31 | NLKPFERDIS |
| P0DTC2 | Spike glycoprotein | 1206 | 406.8885 | 1217.6436 | 1217.6404 | 3 | 0.0032 | 5.05 | NLKPFERDIS |
| P0DTC2 | Spike glycoprotein | 1206 | 609.8291 | 1217.6436 | 1217.6404 | 2 | 0.0032 | 9.2 | NLKPFERDIS |
| P0DTC2 | Spike glycoprotein | 1206 | 609.8291 | 1217.6437 | 1217.6404 | 2 | 0.0033 | 17.35 | NLKPFERDIS |
| P0DTC2 | Spike glycoprotein | 1206 | 406.8885 | 1217.6437 | 1217.6404 | 3 | 0.0033 | 6.52 | NLKPFERDIS |
| P0DTC2 | Spike glycoprotein | 1206 | 609.8292 | 1217.6439 | 1217.6404 | 2 | 0.0035 | 17.54 | NLKPFERDIS |
| P0DTC2 | Spike glycoprotein | 1206 | 609.8293 | 1217.644 | 1217.6404 | 2 | 0.0036 | 12.11 | NLKPFERDIS |
| P0DTC2 | Spike glycoprotein | 1206 | 609.8293 | 1217.644 | 1217.6404 | 2 | 0.0036 | 11.58 | NLKPFERDIS |
| P0DTC2 | Spike glycoprotein | 1206 | 406.8887 | 1217.6443 | 1217.6404 | 3 | 0.0039 | 2.13 | NLKPFERDIS |
| P0DTC2 | Spike glycoprotein | 1206 | 609.8295 | 1217.6444 | 1217.6404 | 2 | 0.004 | 15.75 | NLKPFERDIS |
| P0DTC2 | Spike glycoprotein | 1206 | 406.8888 | 1217.6445 | 1217.6404 | 3 | 0.0041 | 3.13 | NLKPFERDIS |

| | | | | | | | | | |
|---|---|---|---|---|---|---|---|---|---|
| P0DTC2 | Spike glycoprotein | 1206 | 406.8888 | 1217.6446 | 1217.6404 | 3 | 0.0042 | 12.52 | NLKPFERDIS |
| P0DTC2 | Spike glycoprotein | 1206 | 406.8889 | 1217.6448 | 1217.6404 | 3 | 0.0044 | 6.64 | NLKPFERDIS |
| P0DTC2 | Spike glycoprotein | 1206 | 609.8297 | 1217.6449 | 1217.6404 | 2 | 0.0044 | 12 | NLKPFERDIS |
| P0DTC2 | Spike glycoprotein | 1206 | 406.8889 | 1217.645 | 1217.6404 | 3 | 0.0046 | 11.74 | NLKPFERDIS |
| P0DTC2 | Spike glycoprotein | 1206 | 609.8299 | 1217.6453 | 1217.6404 | 2 | 0.0049 | 2.05 | NLKPFERDIS |
| P0DTC2 | Spike glycoprotein | 1206 | 406.8892 | 1217.6457 | 1217.6404 | 3 | 0.0053 | 7.28 | NLKPFERDIS |
| P0DTC2 | Spike glycoprotein | 1206 | 406.8894 | 1217.6462 | 1217.6404 | 3 | 0.0058 | 3.6 | NLKPFERDIS |
| P0DTC2 | Spike glycoprotein | 1206 | 609.8304 | 1217.6463 | 1217.6404 | 2 | 0.0059 | 11.09 | NLKPFERDIS |
| P0DTC2 | Spike glycoprotein | 1206 | 609.8306 | 1217.6466 | 1217.6404 | 2 | 0.0062 | 11.15 | NLKPFERDIS |
| P0DTC2 | Spike glycoprotein | 1206 | 609.8315 | 1217.6485 | 1217.6404 | 2 | 0.0081 | 4.97 | NLKPFERDIS |
| P0DTC2 | Spike glycoprotein | 1206 | 614.2688 | 1226.5231 | 1226.5213 | 2 | 0.0018 | 3.56 | DFCGKGYHLM |
| P0DTC2 | Spike glycoprotein | 1206 | 616.8078 | 1231.601 | 1231.5985 | 2 | 0.0025 | 24.34 | GGNYNYLYRL |
| P0DTC2 | Spike glycoprotein | 1206 | 617.298 | 1232.5815 | 1232.5786 | 2 | 0.003 | 4.8 | WNSNNLDSKVG |
| P0DTC2 | Spike glycoprotein | 1206 | 620.3646 | 1238.7146 | 1238.7095 | 2 | 0.0051 | 42.19 | RQIAPGQTGKIA |
| P0DTC2 | Spike glycoprotein | 1206 | 622.3283 | 1242.642 | 1242.6397 | 2 | 0.0023 | 10.54 | RGVYYPDKVF |
| P0DTC2 | Spike glycoprotein | 1206 | 622.3315 | 1242.6484 | 1242.6397 | 2 | 0.0087 | 9.57 | RGVYYPDKVF |
| P0DTC2 | Spike glycoprotein | 1206 | 623.3322 | 1244.6498 | 1244.6513 | 2 | -0.0016 | 16.25 | NLVRDLPQGFS |
| P0DTC2 | Spike glycoprotein | 1206 | 623.3326 | 1244.6507 | 1244.6513 | 2 | -0.0006 | 19.36 | NLVRDLPQGFS |
| P0DTC2 | Spike glycoprotein | 1206 | 623.3326 | 1244.6507 | 1244.6513 | 2 | -0.0006 | 34.33 | NLVRDLPQGFS |
| P0DTC2 | Spike glycoprotein | 1206 | 623.3327 | 1244.6509 | 1244.6513 | 2 | -0.0004 | 20.3 | NLVRDLPQGFS |
| P0DTC2 | Spike glycoprotein | 1206 | 623.3328 | 1244.651 | 1244.6513 | 2 | -0.0003 | 22.36 | NLVRDLPQGFS |

| | | | | | | | | | |
|---|---|---|---|---|---|---|---|---|---|
| P0DTC2 | Spike glycoprotein | 1206 | 623.3332 | 1244.6518 | 1244.6513 | 2 | 0.0005 | 19.56 | NLVRDLPQGFS |
| P0DTC2 | Spike glycoprotein | 1206 | 623.3337 | 1244.6529 | 1244.6513 | 2 | 0.0016 | 26.06 | NLVRDLPQGFS |
| P0DTC2 | Spike glycoprotein | 1206 | 623.334 | 1244.6535 | 1244.6513 | 2 | 0.0021 | 22.49 | NLVRDLPQGFS |
| P0DTC2 | Spike glycoprotein | 1206 | 623.334 | 1244.6535 | 1244.6513 | 2 | 0.0022 | 37.95 | NLVRDLPQGFS |
| P0DTC2 | Spike glycoprotein | 1206 | 623.3341 | 1244.6537 | 1244.6513 | 2 | 0.0023 | 41.58 | NLVRDLPQGFS |
| P0DTC2 | Spike glycoprotein | 1206 | 623.3341 | 1244.6537 | 1244.6513 | 2 | 0.0023 | 15.7 | NLVRDLPQGFS |
| P0DTC2 | Spike glycoprotein | 1206 | 623.3343 | 1244.6541 | 1244.6513 | 2 | 0.0028 | 48.2 | NLVRDLPQGFS |
| P0DTC2 | Spike glycoprotein | 1206 | 623.3345 | 1244.6545 | 1244.6513 | 2 | 0.0032 | 35.17 | NLVRDLPQGFS |
| P0DTC2 | Spike glycoprotein | 1206 | 623.3346 | 1244.6547 | 1244.6513 | 2 | 0.0034 | 44.48 | NLVRDLPQGFS |
| P0DTC2 | Spike glycoprotein | 1206 | 623.3346 | 1244.6547 | 1244.6513 | 2 | 0.0034 | 45.65 | NLVRDLPQGFS |
| P0DTC2 | Spike glycoprotein | 1206 | 623.3347 | 1244.6548 | 1244.6513 | 2 | 0.0034 | 44.37 | NLVRDLPQGFS |
| P0DTC2 | Spike glycoprotein | 1206 | 623.3347 | 1244.6548 | 1244.6513 | 2 | 0.0034 | 23.93 | NLVRDLPQGFS |
| P0DTC2 | Spike glycoprotein | 1206 | 623.3347 | 1244.6549 | 1244.6513 | 2 | 0.0035 | 46.68 | NLVRDLPQGFS |
| P0DTC2 | Spike glycoprotein | 1206 | 623.3349 | 1244.6553 | 1244.6513 | 2 | 0.004 | 39.4 | NLVRDLPQGFS |
| P0DTC2 | Spike glycoprotein | 1206 | 623.335 | 1244.6554 | 1244.6513 | 2 | 0.0041 | 47.34 | NLVRDLPQGFS |
| P0DTC2 | Spike glycoprotein | 1206 | 623.335 | 1244.6555 | 1244.6513 | 2 | 0.0042 | 27.58 | NLVRDLPQGFS |
| P0DTC2 | Spike glycoprotein | 1206 | 623.3352 | 1244.6559 | 1244.6513 | 2 | 0.0046 | 29.2 | NLVRDLPQGFS |
| P0DTC2 | Spike glycoprotein | 1206 | 623.3354 | 1244.6562 | 1244.6513 | 2 | 0.0049 | 26.76 | NLVRDLPQGFS |
| P0DTC2 | Spike glycoprotein | 1206 | 623.3354 | 1244.6563 | 1244.6513 | 2 | 0.005 | 46.7 | NLVRDLPQGFS |
| P0DTC2 | Spike glycoprotein | 1206 | 623.3361 | 1244.6576 | 1244.6513 | 2 | 0.0063 | 44.13 | NLVRDLPQGFS |
| P0DTC2 | Spike glycoprotein | 1206 | 623.3365 | 1244.6584 | 1244.6513 | 2 | 0.0071 | 29.06 | NLVRDLPQGFS |

| | | | | | | | | | |
|---|---|---|---|---|---|---|---|---|---|
| P0DTC2 | Spike glycoprotein | 1206 | 623.3371 | 1244.6596 | 1244.6513 | 2 | 0.0083 | 37.62 | NLVRDLPQGFS |
| P0DTC2 | Spike glycoprotein | 1206 | 623.3376 | 1244.6607 | 1244.6513 | 2 | 0.0094 | 12.3 | NLVRDLPQGFS |
| P0DTC2 | Spike glycoprotein | 1206 | 626.3244 | 1250.6342 | 1250.6408 | 2 | -0.0066 | 2.98 | NGVGYQPYRVV |
| P0DTC2 | Spike glycoprotein | 1206 | 626.3281 | 1250.6416 | 1250.6408 | 2 | 0.0008 | 13.92 | NGVGYQPYRVV |
| P0DTC2 | Spike glycoprotein | 1206 | 626.3286 | 1250.6426 | 1250.6408 | 2 | 0.0018 | 6.91 | NGVGYQPYRVV |
| P0DTC2 | Spike glycoprotein | 1206 | 626.3289 | 1250.6432 | 1250.6408 | 2 | 0.0024 | 25.44 | NGVGYQPYRVV |
| P0DTC2 | Spike glycoprotein | 1206 | 627.8577 | 1253.7008 | 1253.6979 | 2 | 0.0028 | 3.79 | AVRDPQTLEIL |
| P0DTC2 | Spike glycoprotein | 1206 | 629.3013 | 1256.5881 | 1256.586 | 2 | 0.0021 | 31.75 | GFIKQYGDCLG |
| P0DTC2 | Spike glycoprotein | 1206 | 631.3312 | 1260.6478 | 1260.6462 | 2 | 0.0016 | 21.18 | TRTQLPPAYTN |
| P0DTC2 | Spike glycoprotein | 1206 | 631.332 | 1260.6494 | 1260.6462 | 2 | 0.0032 | 34.4 | TRTQLPPAYTN |
| P0DTC2 | Spike glycoprotein | 1206 | 631.3321 | 1260.6497 | 1260.6462 | 2 | 0.0035 | 34.26 | TRTQLPPAYTN |
| P0DTC2 | Spike glycoprotein | 1206 | 631.3322 | 1260.6498 | 1260.6462 | 2 | 0.0035 | 30.55 | TRTQLPPAYTN |
| P0DTC2 | Spike glycoprotein | 1206 | 631.3322 | 1260.6498 | 1260.6462 | 2 | 0.0035 | 26.03 | TRTQLPPAYTN |
| P0DTC2 | Spike glycoprotein | 1206 | 631.3322 | 1260.6499 | 1260.6462 | 2 | 0.0036 | 26.06 | TRTQLPPAYTN |
| P0DTC2 | Spike glycoprotein | 1206 | 631.3324 | 1260.6502 | 1260.6462 | 2 | 0.004 | 12.26 | TRTQLPPAYTN |
| P0DTC2 | Spike glycoprotein | 1206 | 631.3327 | 1260.6508 | 1260.6462 | 2 | 0.0046 | 34.49 | TRTQLPPAYTN |
| P0DTC2 | Spike glycoprotein | 1206 | 631.3328 | 1260.6511 | 1260.6462 | 2 | 0.0049 | 25.93 | TRTQLPPAYTN |
| P0DTC2 | Spike glycoprotein | 1206 | 631.3329 | 1260.6513 | 1260.6462 | 2 | 0.0051 | 23.56 | TRTQLPPAYTN |
| P0DTC2 | Spike glycoprotein | 1206 | 631.333 | 1260.6514 | 1260.6462 | 2 | 0.0052 | 34.39 | TRTQLPPAYTN |
| P0DTC2 | Spike glycoprotein | 1206 | 631.3332 | 1260.6519 | 1260.6462 | 2 | 0.0056 | 14.66 | TRTQLPPAYTN |
| P0DTC2 | Spike glycoprotein | 1206 | 631.3333 | 1260.6521 | 1260.6462 | 2 | 0.0059 | 37.6 | TRTQLPPAYTN |

| | | | | | | | | | |
|---|---|---|---|---|---|---|---|---|---|
| P0DTC2 | Spike glycoprotein | 1206 | 631.3334 | 1260.6522 | 1260.6462 | 2 | 0.0059 | 25.9 | TRTQLPPAYTN |
| P0DTC2 | Spike glycoprotein | 1206 | 631.3334 | 1260.6522 | 1260.6462 | 2 | 0.006 | 2.23 | TRTQLPPAYTN |
| P0DTC2 | Spike glycoprotein | 1206 | 631.3336 | 1260.6526 | 1260.6462 | 2 | 0.0063 | 10.41 | TRTQLPPAYTN |
| P0DTC2 | Spike glycoprotein | 1206 | 631.3336 | 1260.6526 | 1260.6462 | 2 | 0.0063 | 30.01 | TRTQLPPAYTN |
| P0DTC2 | Spike glycoprotein | 1206 | 631.3339 | 1260.6533 | 1260.6462 | 2 | 0.0071 | 18.27 | TRTQLPPAYTN |
| P0DTC2 | Spike glycoprotein | 1206 | 631.334 | 1260.6535 | 1260.6462 | 2 | 0.0073 | 22.17 | TRTQLPPAYTN |
| P0DTC2 | Spike glycoprotein | 1206 | 635.8243 | 1269.634 | 1269.6313 | 2 | 0.0027 | 22.9 | DEVRQIAPGQTG |
| P0DTC2 | Spike glycoprotein | 1206 | 635.8262 | 1269.6378 | 1269.6313 | 2 | 0.0065 | 8.42 | DEVRQIAPGQTG |
| P0DTC2 | Spike glycoprotein | 1206 | 637.3327 | 1272.6508 | 1272.6503 | 2 | 0.0005 | 39.2 | VYYPDKVFRS |
| P0DTC2 | Spike glycoprotein | 1206 | 425.2245 | 1272.6516 | 1272.6503 | 3 | 0.0013 | 26.98 | VYYPDKVFRS |
| P0DTC2 | Spike glycoprotein | 1206 | 425.2248 | 1272.6527 | 1272.6503 | 3 | 0.0024 | 18.45 | VYYPDKVFRS |
| P0DTC2 | Spike glycoprotein | 1206 | 425.2255 | 1272.6548 | 1272.6503 | 3 | 0.0045 | 19.48 | VYYPDKVFRS |
| P0DTC2 | Spike glycoprotein | 1206 | 425.2273 | 1272.6601 | 1272.6503 | 3 | 0.0098 | 9.74 | VYYPDKVFRS |
| P0DTC2 | Spike glycoprotein | 1206 | 639.3523 | 1276.6901 | 1276.6928 | 2 | -0.0027 | 34.1 | NIIRGWIFGTT |
| P0DTC2 | Spike glycoprotein | 1206 | 639.3529 | 1276.6912 | 1276.6928 | 2 | -0.0017 | 6.11 | NIIRGWIFGTT |
| P0DTC2 | Spike glycoprotein | 1206 | 639.3535 | 1276.6924 | 1276.6928 | 2 | -0.0004 | 22.86 | NIIRGWIFGTT |
| P0DTC2 | Spike glycoprotein | 1206 | 639.3539 | 1276.6932 | 1276.6928 | 2 | 0.0004 | 34.91 | NIIRGWIFGTT |
| P0DTC2 | Spike glycoprotein | 1206 | 639.3544 | 1276.6943 | 1276.6928 | 2 | 0.0015 | 24.56 | NIIRGWIFGTT |
| P0DTC2 | Spike glycoprotein | 1206 | 639.3548 | 1276.6951 | 1276.6928 | 2 | 0.0023 | 46.32 | NIIRGWIFGTT |
| P0DTC2 | Spike glycoprotein | 1206 | 639.355 | 1276.6955 | 1276.6928 | 2 | 0.0027 | 47.44 | NIIRGWIFGTT |
| P0DTC2 | Spike glycoprotein | 1206 | 639.3552 | 1276.6959 | 1276.6928 | 2 | 0.0031 | 45.25 | NIIRGWIFGTT |

| | | | | | | | | | |
|---|---|---|---|---|---|---|---|---|---|
| P0DTC2 | Spike glycoprotein | 1206 | 639.3553 | 1276.696 | 1276.6928 | 2 | 0.0032 | 6.47 | NIIRGWIFGTT |
| P0DTC2 | Spike glycoprotein | 1206 | 639.3554 | 1276.6963 | 1276.6928 | 2 | 0.0034 | 48.7 | NIIRGWIFGTT |
| P0DTC2 | Spike glycoprotein | 1206 | 639.3556 | 1276.6966 | 1276.6928 | 2 | 0.0037 | 16.05 | NIIRGWIFGTT |
| P0DTC2 | Spike glycoprotein | 1206 | 639.3556 | 1276.6966 | 1276.6928 | 2 | 0.0038 | 56.49 | NIIRGWIFGTT |
| P0DTC2 | Spike glycoprotein | 1206 | 639.3556 | 1276.6966 | 1276.6928 | 2 | 0.0038 | 39.18 | NIIRGWIFGTT |
| P0DTC2 | Spike glycoprotein | 1206 | 639.3556 | 1276.6967 | 1276.6928 | 2 | 0.0038 | 43.84 | NIIRGWIFGTT |
| P0DTC2 | Spike glycoprotein | 1206 | 639.3556 | 1276.6967 | 1276.6928 | 2 | 0.0039 | 43.81 | NIIRGWIFGTT |
| P0DTC2 | Spike glycoprotein | 1206 | 639.3557 | 1276.6968 | 1276.6928 | 2 | 0.004 | 22.56 | NIIRGWIFGTT |
| P0DTC2 | Spike glycoprotein | 1206 | 639.3557 | 1276.6968 | 1276.6928 | 2 | 0.004 | 45.3 | NIIRGWIFGTT |
| P0DTC2 | Spike glycoprotein | 1206 | 639.3558 | 1276.6971 | 1276.6928 | 2 | 0.0043 | 44.88 | NIIRGWIFGTT |
| P0DTC2 | Spike glycoprotein | 1206 | 639.3558 | 1276.6971 | 1276.6928 | 2 | 0.0043 | 46.61 | NIIRGWIFGTT |
| P0DTC2 | Spike glycoprotein | 1206 | 639.356 | 1276.6974 | 1276.6928 | 2 | 0.0046 | 41.45 | NIIRGWIFGTT |
| P0DTC2 | Spike glycoprotein | 1206 | 645.3492 | 1288.6838 | 1288.6776 | 2 | 0.0063 | 7.69 | NFRVQPTESIV |
| P0DTC2 | Spike glycoprotein | 1206 | 646.8665 | 1291.7184 | 1291.7149 | 2 | 0.0035 | 5.88 | VYAWNRKRIS |
| P0DTC2 | Spike glycoprotein | 1206 | 646.8676 | 1291.7206 | 1291.7149 | 2 | 0.0057 | 9.94 | VYAWNRKRIS |
| P0DTC2 | Spike glycoprotein | 1206 | 655.862 | 1309.7095 | 1309.7102 | 2 | -0.0007 | 23.88 | IRGDEVRQIAPG |
| P0DTC2 | Spike glycoprotein | 1206 | 655.8636 | 1309.7126 | 1309.7102 | 2 | 0.0024 | 27.99 | IRGDEVRQIAPG |
| P0DTC2 | Spike glycoprotein | 1206 | 437.5783 | 1309.713 | 1309.7102 | 3 | 0.0028 | 11.3 | IRGDEVRQIAPG |
| P0DTC2 | Spike glycoprotein | 1206 | 655.8638 | 1309.7131 | 1309.7102 | 2 | 0.0029 | 25.88 | IRGDEVRQIAPG |
| P0DTC2 | Spike glycoprotein | 1206 | 655.864 | 1309.7134 | 1309.7102 | 2 | 0.0032 | 23.43 | IRGDEVRQIAPG |
| P0DTC2 | Spike glycoprotein | 1206 | 437.5785 | 1309.7136 | 1309.7102 | 3 | 0.0034 | 15.3 | IRGDEVRQIAPG |

| | | | | | | | | | |
|---|---|---|---|---|---|---|---|---|---|
| P0DTC2 | Spike glycoprotein | 1206 | 437.5785 | 1309.7137 | 1309.7102 | 3 | 0.0035 | 10.87 | IRGDEVRQIAPG |
| P0DTC2 | Spike glycoprotein | 1206 | 655.8642 | 1309.7139 | 1309.7102 | 2 | 0.0036 | 30.48 | IRGDEVRQIAPG |
| P0DTC2 | Spike glycoprotein | 1206 | 437.5787 | 1309.7142 | 1309.7102 | 3 | 0.0039 | 4.71 | IRGDEVRQIAPG |
| P0DTC2 | Spike glycoprotein | 1206 | 655.8644 | 1309.7142 | 1309.7102 | 2 | 0.0039 | 15.96 | IRGDEVRQIAPG |
| P0DTC2 | Spike glycoprotein | 1206 | 437.5789 | 1309.7148 | 1309.7102 | 3 | 0.0046 | 3.98 | IRGDEVRQIAPG |
| P0DTC2 | Spike glycoprotein | 1206 | 655.8647 | 1309.7148 | 1309.7102 | 2 | 0.0046 | 11.04 | IRGDEVRQIAPG |
| P0DTC2 | Spike glycoprotein | 1206 | 655.8647 | 1309.7149 | 1309.7102 | 2 | 0.0046 | 12.76 | IRGDEVRQIAPG |
| P0DTC2 | Spike glycoprotein | 1206 | 655.865 | 1309.7155 | 1309.7102 | 2 | 0.0052 | 4.82 | IRGDEVRQIAPG |
| P0DTC2 | Spike glycoprotein | 1206 | 655.8652 | 1309.7158 | 1309.7102 | 2 | 0.0056 | 12.34 | IRGDEVRQIAPG |
| P0DTC2 | Spike glycoprotein | 1206 | 437.5794 | 1309.7163 | 1309.7102 | 3 | 0.0061 | 3.02 | IRGDEVRQIAPG |
| P0DTC2 | Spike glycoprotein | 1206 | 655.8655 | 1309.7164 | 1309.7102 | 2 | 0.0061 | 16.32 | IRGDEVRQIAPG |
| P0DTC2 | Spike glycoprotein | 1206 | 658.8517 | 1315.6889 | 1315.6884 | 2 | 0.0004 | 23.49 | NLVRDLPQGFSA |
| P0DTC2 | Spike glycoprotein | 1206 | 658.8518 | 1315.689 | 1315.6884 | 2 | 0.0005 | 4.79 | NLVRDLPQGFSA |
| P0DTC2 | Spike glycoprotein | 1206 | 658.8526 | 1315.6907 | 1315.6884 | 2 | 0.0023 | 41.47 | NLVRDLPQGFSA |
| P0DTC2 | Spike glycoprotein | 1206 | 658.8528 | 1315.691 | 1315.6884 | 2 | 0.0026 | 66.23 | NLVRDLPQGFSA |
| P0DTC2 | Spike glycoprotein | 1206 | 658.8528 | 1315.691 | 1315.6884 | 2 | 0.0026 | 6.94 | NLVRDLPQGFSA |
| P0DTC2 | Spike glycoprotein | 1206 | 658.8528 | 1315.6911 | 1315.6884 | 2 | 0.0026 | 16.15 | NLVRDLPQGFSA |
| P0DTC2 | Spike glycoprotein | 1206 | 658.8529 | 1315.6912 | 1315.6884 | 2 | 0.0028 | 14.92 | NLVRDLPQGFSA |
| P0DTC2 | Spike glycoprotein | 1206 | 658.8529 | 1315.6912 | 1315.6884 | 2 | 0.0028 | 47.17 | NLVRDLPQGFSA |
| P0DTC2 | Spike glycoprotein | 1206 | 658.8529 | 1315.6913 | 1315.6884 | 2 | 0.0028 | 39.51 | NLVRDLPQGFSA |
| P0DTC2 | Spike glycoprotein | 1206 | 658.853 | 1315.6915 | 1315.6884 | 2 | 0.0031 | 24.07 | NLVRDLPQGFSA |

| | | | | | | | | | |
|---|---|---|---|---|---|---|---|---|---|
| P0DTC2 | Spike glycoprotein | 1206 | 658.8531 | 1315.6915 | 1315.6884 | 2 | 0.0031 | 32.5 | NLVRDLPQGFSA |
| P0DTC2 | Spike glycoprotein | 1206 | 658.8531 | 1315.6916 | 1315.6884 | 2 | 0.0032 | 66.21 | NLVRDLPQGFSA |
| P0DTC2 | Spike glycoprotein | 1206 | 658.8531 | 1315.6916 | 1315.6884 | 2 | 0.0032 | 45.55 | NLVRDLPQGFSA |
| P0DTC2 | Spike glycoprotein | 1206 | 658.8531 | 1315.6916 | 1315.6884 | 2 | 0.0032 | 50.93 | NLVRDLPQGFSA |
| P0DTC2 | Spike glycoprotein | 1206 | 658.8531 | 1315.6917 | 1315.6884 | 2 | 0.0032 | 66.25 | NLVRDLPQGFSA |
| P0DTC2 | Spike glycoprotein | 1206 | 658.8532 | 1315.6918 | 1315.6884 | 2 | 0.0034 | 25.73 | NLVRDLPQGFSA |
| P0DTC2 | Spike glycoprotein | 1206 | 658.8532 | 1315.6919 | 1315.6884 | 2 | 0.0034 | 22.43 | NLVRDLPQGFSA |
| P0DTC2 | Spike glycoprotein | 1206 | 658.8532 | 1315.6919 | 1315.6884 | 2 | 0.0035 | 26.27 | NLVRDLPQGFSA |
| P0DTC2 | Spike glycoprotein | 1206 | 658.8533 | 1315.692 | 1315.6884 | 2 | 0.0035 | 49.65 | NLVRDLPQGFSA |
| P0DTC2 | Spike glycoprotein | 1206 | 658.8533 | 1315.6921 | 1315.6884 | 2 | 0.0036 | 23.82 | NLVRDLPQGFSA |
| P0DTC2 | Spike glycoprotein | 1206 | 658.8535 | 1315.6924 | 1315.6884 | 2 | 0.0039 | 62.23 | NLVRDLPQGFSA |
| P0DTC2 | Spike glycoprotein | 1206 | 658.8535 | 1315.6924 | 1315.6884 | 2 | 0.004 | 25.05 | NLVRDLPQGFSA |
| P0DTC2 | Spike glycoprotein | 1206 | 658.8536 | 1315.6926 | 1315.6884 | 2 | 0.0042 | 50.89 | NLVRDLPQGFSA |
| P0DTC2 | Spike glycoprotein | 1206 | 658.8537 | 1315.6928 | 1315.6884 | 2 | 0.0044 | 27.5 | NLVRDLPQGFSA |
| P0DTC2 | Spike glycoprotein | 1206 | 658.8538 | 1315.693 | 1315.6884 | 2 | 0.0046 | 24.19 | NLVRDLPQGFSA |
| P0DTC2 | Spike glycoprotein | 1206 | 658.8538 | 1315.6931 | 1315.6884 | 2 | 0.0047 | 14.51 | NLVRDLPQGFSA |
| P0DTC2 | Spike glycoprotein | 1206 | 658.8539 | 1315.6932 | 1315.6884 | 2 | 0.0048 | 31.19 | NLVRDLPQGFSA |
| P0DTC2 | Spike glycoprotein | 1206 | 658.8543 | 1315.6941 | 1315.6884 | 2 | 0.0056 | 28.6 | NLVRDLPQGFSA |
| P0DTC2 | Spike glycoprotein | 1206 | 658.8544 | 1315.6943 | 1315.6884 | 2 | 0.0058 | 18.26 | NLVRDLPQGFSA |
| P0DTC2 | Spike glycoprotein | 1206 | 658.8546 | 1315.6947 | 1315.6884 | 2 | 0.0063 | 27.58 | NLVRDLPQGFSA |
| P0DTC2 | Spike glycoprotein | 1206 | 658.8561 | 1315.6977 | 1315.6884 | 2 | 0.0093 | 43.61 | NLVRDLPQGFSA |

| | | | | | | | | | |
|---|---|---|---|---|---|---|---|---|---|
| P0DTC2 | Spike glycoprotein | 1206 | 660.3511 | 1318.6876 | 1318.6881 | 2 | -0.0005 | 17.18 | NLKPFERDIST |
| P0DTC2 | Spike glycoprotein | 1206 | 660.3514 | 1318.6882 | 1318.6881 | 2 | 0.0001 | 13.78 | NLKPFERDIST |
| P0DTC2 | Spike glycoprotein | 1206 | 660.3514 | 1318.6883 | 1318.6881 | 2 | 0.0002 | 6.39 | NLKPFERDIST |
| P0DTC2 | Spike glycoprotein | 1206 | 660.3517 | 1318.6888 | 1318.6881 | 2 | 0.0007 | 13.7 | NLKPFERDIST |
| P0DTC2 | Spike glycoprotein | 1206 | 660.3518 | 1318.6891 | 1318.6881 | 2 | 0.001 | 15.4 | NLKPFERDIST |
| P0DTC2 | Spike glycoprotein | 1206 | 660.3519 | 1318.6892 | 1318.6881 | 2 | 0.0011 | 19.54 | NLKPFERDIST |
| P0DTC2 | Spike glycoprotein | 1206 | 660.3519 | 1318.6892 | 1318.6881 | 2 | 0.0011 | 15.58 | NLKPFERDIST |
| P0DTC2 | Spike glycoprotein | 1206 | 660.352 | 1318.6895 | 1318.6881 | 2 | 0.0014 | 21.15 | NLKPFERDIST |
| P0DTC2 | Spike glycoprotein | 1206 | 440.5707 | 1318.6901 | 1318.6881 | 3 | 0.002 | 1.85 | NLKPFERDIST |
| P0DTC2 | Spike glycoprotein | 1206 | 660.3524 | 1318.6903 | 1318.6881 | 2 | 0.0022 | 14.18 | NLKPFERDIST |
| P0DTC2 | Spike glycoprotein | 1206 | 660.3525 | 1318.6904 | 1318.6881 | 2 | 0.0023 | 10.56 | NLKPFERDIST |
| P0DTC2 | Spike glycoprotein | 1206 | 660.3525 | 1318.6904 | 1318.6881 | 2 | 0.0023 | 15.92 | NLKPFERDIST |
| P0DTC2 | Spike glycoprotein | 1206 | 660.3525 | 1318.6905 | 1318.6881 | 2 | 0.0024 | 9.09 | NLKPFERDIST |
| P0DTC2 | Spike glycoprotein | 1206 | 660.3526 | 1318.6907 | 1318.6881 | 2 | 0.0026 | 12.2 | NLKPFERDIST |
| P0DTC2 | Spike glycoprotein | 1206 | 660.3529 | 1318.6913 | 1318.6881 | 2 | 0.0032 | 19.43 | NLKPFERDIST |
| P0DTC2 | Spike glycoprotein | 1206 | 660.3531 | 1318.6916 | 1318.6881 | 2 | 0.0035 | 14.13 | NLKPFERDIST |
| P0DTC2 | Spike glycoprotein | 1206 | 440.5712 | 1318.6917 | 1318.6881 | 3 | 0.0036 | 6.07 | NLKPFERDIST |
| P0DTC2 | Spike glycoprotein | 1206 | 660.3532 | 1318.6918 | 1318.6881 | 2 | 0.0037 | 24.17 | NLKPFERDIST |
| P0DTC2 | Spike glycoprotein | 1206 | 660.3533 | 1318.692 | 1318.6881 | 2 | 0.0039 | 19.39 | NLKPFERDIST |
| P0DTC2 | Spike glycoprotein | 1206 | 660.3533 | 1318.692 | 1318.6881 | 2 | 0.0039 | 20.65 | NLKPFERDIST |
| P0DTC2 | Spike glycoprotein | 1206 | 660.3533 | 1318.692 | 1318.6881 | 2 | 0.0039 | 16.27 | NLKPFERDIST |

| | | | | | | | | | |
|---|---|---|---|---|---|---|---|---|---|
| P0DTC2 | Spike glycoprotein | 1206 | 660.3534 | 1318.6922 | 1318.6881 | 2 | 0.0041 | 10.01 | NLKPFERDIST |
| P0DTC2 | Spike glycoprotein | 1206 | 660.3536 | 1318.6926 | 1318.6881 | 2 | 0.0045 | 18.3 | NLKPFERDIST |
| P0DTC2 | Spike glycoprotein | 1206 | 660.3537 | 1318.6928 | 1318.6881 | 2 | 0.0047 | 19.43 | NLKPFERDIST |
| P0DTC2 | Spike glycoprotein | 1206 | 660.3537 | 1318.6929 | 1318.6881 | 2 | 0.0048 | 10.11 | NLKPFERDIST |
| P0DTC2 | Spike glycoprotein | 1206 | 440.5716 | 1318.693 | 1318.6881 | 3 | 0.0049 | 2.92 | NLKPFERDIST |
| P0DTC2 | Spike glycoprotein | 1206 | 660.3538 | 1318.6931 | 1318.6881 | 2 | 0.005 | 17.55 | NLKPFERDIST |
| P0DTC2 | Spike glycoprotein | 1206 | 660.3542 | 1318.6938 | 1318.6881 | 2 | 0.0057 | 16.88 | NLKPFERDIST |
| P0DTC2 | Spike glycoprotein | 1206 | 669.8416 | 1337.6687 | 1337.6656 | 2 | 0.0031 | 15.51 | PVLPFNDGVYFA |
| P0DTC2 | Spike glycoprotein | 1206 | 669.8422 | 1337.6698 | 1337.6656 | 2 | 0.0042 | 17.65 | PVLPFNDGVYFA |
| P0DTC2 | Spike glycoprotein | 1206 | 672.3635 | 1342.7124 | 1342.7092 | 2 | 0.0031 | 7.95 | EAEVQIDRLITG |
| P0DTC2 | Spike glycoprotein | 1206 | 448.9036 | 1343.689 | 1343.6874 | 3 | 0.0016 | 13.81 | FTRGVYYPDKV |
| P0DTC2 | Spike glycoprotein | 1206 | 451.9164 | 1352.7273 | 1352.7241 | 3 | 0.0032 | 10.53 | NKKFLPFQQFG |
| P0DTC2 | Spike glycoprotein | 1206 | 677.3718 | 1352.729 | 1352.7241 | 2 | 0.0049 | 10.31 | NKKFLPFQQFG |
| P0DTC2 | Spike glycoprotein | 1206 | 454.2355 | 1359.6845 | 1359.6823 | 3 | 0.0022 | 22.52 | VYYPDKVFRSS |
| P0DTC2 | Spike glycoprotein | 1206 | 681.858 | 1361.7015 | 1361.6939 | 2 | 0.0076 | 7.14 | DLEGKQGNFKNL |
| P0DTC2 | Spike glycoprotein | 1206 | 685.3693 | 1368.7241 | 1368.7249 | 2 | -0.0008 | 0.79 | DAVRDPQTLEIL |
| P0DTC2 | Spike glycoprotein | 1206 | 685.3702 | 1368.7259 | 1368.7249 | 2 | 0.001 | 9.08 | DAVRDPQTLEIL |
| P0DTC2 | Spike glycoprotein | 1206 | 685.3705 | 1368.7264 | 1368.7249 | 2 | 0.0016 | 44.42 | DAVRDPQTLEIL |
| P0DTC2 | Spike glycoprotein | 1206 | 685.3707 | 1368.7269 | 1368.7249 | 2 | 0.002 | 31.87 | DAVRDPQTLEIL |
| P0DTC2 | Spike glycoprotein | 1206 | 685.3708 | 1368.727 | 1368.7249 | 2 | 0.0021 | 26 | DAVRDPQTLEIL |
| P0DTC2 | Spike glycoprotein | 1206 | 685.3708 | 1368.7271 | 1368.7249 | 2 | 0.0022 | 28.77 | DAVRDPQTLEIL |

| | | | | | | | | | |
|---|---|---|---|---|---|---|---|---|---|
| P0DTC2 | Spike glycoprotein | 1206 | 685.3709 | 1368.7272 | 1368.7249 | 2 | 0.0023 | 35.22 | DAVRDPQTLEIL |
| P0DTC2 | Spike glycoprotein | 1206 | 685.3711 | 1368.7277 | 1368.7249 | 2 | 0.0028 | 43.13 | DAVRDPQTLEIL |
| P0DTC2 | Spike glycoprotein | 1206 | 685.3712 | 1368.7278 | 1368.7249 | 2 | 0.0029 | 21.94 | DAVRDPQTLEIL |
| P0DTC2 | Spike glycoprotein | 1206 | 685.3712 | 1368.7279 | 1368.7249 | 2 | 0.003 | 36.69 | DAVRDPQTLEIL |
| P0DTC2 | Spike glycoprotein | 1206 | 685.3712 | 1368.7279 | 1368.7249 | 2 | 0.003 | 26.93 | DAVRDPQTLEIL |
| P0DTC2 | Spike glycoprotein | 1206 | 685.3712 | 1368.7279 | 1368.7249 | 2 | 0.003 | 32.28 | DAVRDPQTLEIL |
| P0DTC2 | Spike glycoprotein | 1206 | 685.3714 | 1368.7282 | 1368.7249 | 2 | 0.0033 | 28.31 | DAVRDPQTLEIL |
| P0DTC2 | Spike glycoprotein | 1206 | 685.3715 | 1368.7284 | 1368.7249 | 2 | 0.0035 | 33.45 | DAVRDPQTLEIL |
| P0DTC2 | Spike glycoprotein | 1206 | 685.3716 | 1368.7286 | 1368.7249 | 2 | 0.0037 | 30.63 | DAVRDPQTLEIL |
| P0DTC2 | Spike glycoprotein | 1206 | 685.3716 | 1368.7287 | 1368.7249 | 2 | 0.0038 | 41.47 | DAVRDPQTLEIL |
| P0DTC2 | Spike glycoprotein | 1206 | 685.3716 | 1368.7287 | 1368.7249 | 2 | 0.0038 | 28.78 | DAVRDPQTLEIL |
| P0DTC2 | Spike glycoprotein | 1206 | 685.3717 | 1368.7289 | 1368.7249 | 2 | 0.004 | 17.94 | DAVRDPQTLEIL |
| P0DTC2 | Spike glycoprotein | 1206 | 685.3717 | 1368.7289 | 1368.7249 | 2 | 0.004 | 26.98 | DAVRDPQTLEIL |
| P0DTC2 | Spike glycoprotein | 1206 | 685.3718 | 1368.729 | 1368.7249 | 2 | 0.0041 | 18.35 | DAVRDPQTLEIL |
| P0DTC2 | Spike glycoprotein | 1206 | 685.3718 | 1368.729 | 1368.7249 | 2 | 0.0041 | 19.96 | DAVRDPQTLEIL |
| P0DTC2 | Spike glycoprotein | 1206 | 685.3718 | 1368.7291 | 1368.7249 | 2 | 0.0042 | 28.77 | DAVRDPQTLEIL |
| P0DTC2 | Spike glycoprotein | 1206 | 685.3722 | 1368.7298 | 1368.7249 | 2 | 0.0049 | 29.4 | DAVRDPQTLEIL |
| P0DTC2 | Spike glycoprotein | 1206 | 685.3723 | 1368.73 | 1368.7249 | 2 | 0.0051 | 27.14 | DAVRDPQTLEIL |
| P0DTC2 | Spike glycoprotein | 1206 | 685.3726 | 1368.7306 | 1368.7249 | 2 | 0.0058 | 17.68 | DAVRDPQTLEIL |
| P0DTC2 | Spike glycoprotein | 1206 | 685.3733 | 1368.732 | 1368.7249 | 2 | 0.0071 | 16.62 | DAVRDPQTLEIL |
| P0DTC2 | Spike glycoprotein | 1206 | 685.3747 | 1368.7348 | 1368.7249 | 2 | 0.0099 | 33.59 | DAVRDPQTLEIL |

| | | | | | | | | | |
|---|---|---|---|---|---|---|---|---|---|
| P0DTC2 | Spike glycoprotein | 1206 | 685.375 | 1368.7355 | 1368.7249 | 2 | 0.0106 | 45.68 | DAVRDPQTLEIL |
| P0DTC2 | Spike glycoprotein | 1206 | 685.8342 | 1369.6539 | 1369.6514 | 2 | 0.0025 | 31.28 | YADSFVIRGDEV |
| P0DTC2 | Spike glycoprotein | 1206 | 685.8355 | 1369.6565 | 1369.6514 | 2 | 0.0051 | 45.3 | YADSFVIRGDEV |
| P0DTC2 | Spike glycoprotein | 1206 | 685.8372 | 1369.6599 | 1369.6514 | 2 | 0.0085 | 7.15 | YADSFVIRGDEV |
| P0DTC2 | Spike glycoprotein | 1206 | 460.5908 | 1378.7507 | 1378.7469 | 3 | 0.0037 | 15.62 | SVYAWNRKRIS |
| P0DTC2 | Spike glycoprotein | 1206 | 460.5909 | 1378.7508 | 1378.7469 | 3 | 0.0039 | 8.83 | SVYAWNRKRIS |
| P0DTC2 | Spike glycoprotein | 1206 | 690.3832 | 1378.7518 | 1378.7469 | 2 | 0.0049 | 9.08 | SVYAWNRKRIS |
| P0DTC2 | Spike glycoprotein | 1206 | 691.3418 | 1380.669 | 1380.6674 | 2 | 0.0017 | 0.69 | ELDKYFKNHTS |
| P0DTC2 | Spike glycoprotein | 1206 | 695.813 | 1389.6114 | 1389.6089 | 2 | 0.0025 | 9.18 | DYNYKLPDDFT |
| P0DTC2 | Spike glycoprotein | 1206 | 695.8136 | 1389.6127 | 1389.6089 | 2 | 0.0038 | 16.71 | DYNYKLPDDFT |
| P0DTC2 | Spike glycoprotein | 1206 | 695.8137 | 1389.6128 | 1389.6089 | 2 | 0.0039 | 14.89 | DYNYKLPDDFT |
| P0DTC2 | Spike glycoprotein | 1206 | 695.8137 | 1389.6129 | 1389.6089 | 2 | 0.0041 | 13.51 | DYNYKLPDDFT |
| P0DTC2 | Spike glycoprotein | 1206 | 695.8138 | 1389.6131 | 1389.6089 | 2 | 0.0042 | 18.27 | DYNYKLPDDFT |
| P0DTC2 | Spike glycoprotein | 1206 | 697.3275 | 1392.6405 | 1392.6384 | 2 | 0.0021 | 2.64 | FSTFKCYGVSPT |
| P0DTC2 | Spike glycoprotein | 1206 | 697.3279 | 1392.6413 | 1392.6384 | 2 | 0.0029 | 2.19 | FSTFKCYGVSPT |
| P0DTC2 | Spike glycoprotein | 1206 | 697.3286 | 1392.6426 | 1392.6384 | 2 | 0.0042 | 5.64 | FSTFKCYGVSPT |
| P0DTC2 | Spike glycoprotein | 1206 | 467.2555 | 1398.7447 | 1398.7408 | 3 | 0.0038 | 18.42 | RGVYYPDKVFR |
| P0DTC2 | Spike glycoprotein | 1206 | 704.866 | 1407.7174 | 1407.7146 | 2 | 0.0027 | 8.67 | QRNFYEPQIIT |
| P0DTC2 | Spike glycoprotein | 1206 | 713.8753 | 1425.7361 | 1425.7324 | 2 | 0.0036 | 25.37 | RGDEVRQIAPGQT |
| P0DTC2 | Spike glycoprotein | 1206 | 713.8756 | 1425.7366 | 1425.7324 | 2 | 0.0042 | 13.44 | RGDEVRQIAPGQT |
| P0DTC2 | Spike glycoprotein | 1206 | 476.2529 | 1425.7367 | 1425.7324 | 3 | 0.0043 | 21.26 | RGDEVRQIAPGQT |

| | | | | | | | | | |
|---|---|---|---|---|---|---|---|---|---|
| P0DTC2 | Spike glycoprotein | 1206 | 713.8759 | 1425.7372 | 1425.7324 | 2 | 0.0048 | 4.06 | RGDEVRQIAPGQT |
| P0DTC2 | Spike glycoprotein | 1206 | 713.8761 | 1425.7377 | 1425.7324 | 2 | 0.0052 | 1.28 | RGDEVRQIAPGQT |
| P0DTC2 | Spike glycoprotein | 1206 | 713.8764 | 1425.7382 | 1425.7324 | 2 | 0.0058 | 12.3 | RGDEVRQIAPGQT |
| P0DTC2 | Spike glycoprotein | 1206 | 719.8788 | 1437.7431 | 1437.7405 | 2 | 0.0026 | 41.54 | FLPFQQFGRDIA |
| P0DTC2 | Spike glycoprotein | 1206 | 719.8788 | 1437.7431 | 1437.7405 | 2 | 0.0026 | 41.38 | FLPFQQFGRDIA |
| P0DTC2 | Spike glycoprotein | 1206 | 719.8789 | 1437.7433 | 1437.7405 | 2 | 0.0028 | 41.48 | FLPFQQFGRDIA |
| P0DTC2 | Spike glycoprotein | 1206 | 719.8795 | 1437.7445 | 1437.7405 | 2 | 0.004 | 41.31 | FLPFQQFGRDIA |
| P0DTC2 | Spike glycoprotein | 1206 | 719.8807 | 1437.7468 | 1437.7405 | 2 | 0.0063 | 44.66 | FLPFQQFGRDIA |
| P0DTC2 | Spike glycoprotein | 1206 | 721.8863 | 1441.7581 | 1441.7565 | 2 | 0.0016 | 27.68 | RDLPQGFSALEPL |
| P0DTC2 | Spike glycoprotein | 1206 | 721.8866 | 1441.7585 | 1441.7565 | 2 | 0.002 | 45.49 | RDLPQGFSALEPL |
| P0DTC2 | Spike glycoprotein | 1206 | 722.3341 | 1442.6536 | 1442.65 | 2 | 0.0036 | 38.89 | DAGFIKQYGDCLG |
| P0DTC2 | Spike glycoprotein | 1206 | 722.3343 | 1442.654 | 1442.65 | 2 | 0.004 | 40.63 | DAGFIKQYGDCLG |
| P0DTC2 | Spike glycoprotein | 1206 | 724.3222 | 1446.6299 | 1446.6303 | 2 | -0.0004 | 1.43 | DYNYKLPDDFTG |
| P0DTC2 | Spike glycoprotein | 1206 | 724.3227 | 1446.6309 | 1446.6303 | 2 | 0.0006 | 13.39 | DYNYKLPDDFTG |
| P0DTC2 | Spike glycoprotein | 1206 | 724.3229 | 1446.6313 | 1446.6303 | 2 | 0.001 | 20.85 | DYNYKLPDDFTG |
| P0DTC2 | Spike glycoprotein | 1206 | 724.3232 | 1446.6319 | 1446.6303 | 2 | 0.0016 | 14.56 | DYNYKLPDDFTG |
| P0DTC2 | Spike glycoprotein | 1206 | 724.3235 | 1446.6324 | 1446.6303 | 2 | 0.002 | 26.37 | DYNYKLPDDFTG |
| P0DTC2 | Spike glycoprotein | 1206 | 724.3235 | 1446.6324 | 1446.6303 | 2 | 0.0021 | 19.6 | DYNYKLPDDFTG |
| P0DTC2 | Spike glycoprotein | 1206 | 724.3236 | 1446.6326 | 1446.6303 | 2 | 0.0023 | 10.12 | DYNYKLPDDFTG |
| P0DTC2 | Spike glycoprotein | 1206 | 724.3237 | 1446.6328 | 1446.6303 | 2 | 0.0025 | 9.34 | DYNYKLPDDFTG |
| P0DTC2 | Spike glycoprotein | 1206 | 724.3238 | 1446.633 | 1446.6303 | 2 | 0.0026 | 19.39 | DYNYKLPDDFTG |

| | | | | | | | | | |
|---|---|---|---|---|---|---|---|---|---|
| P0DTC2 | Spike glycoprotein | 1206 | 724.3238 | 1446.633 | 1446.6303 | 2 | 0.0027 | 20.5 | DYNYKLPDDFTG |
| P0DTC2 | Spike glycoprotein | 1206 | 724.324 | 1446.6334 | 1446.6303 | 2 | 0.0031 | 8.53 | DYNYKLPDDFTG |
| P0DTC2 | Spike glycoprotein | 1206 | 724.3241 | 1446.6335 | 1446.6303 | 2 | 0.0032 | 8.15 | DYNYKLPDDFTG |
| P0DTC2 | Spike glycoprotein | 1206 | 724.3241 | 1446.6336 | 1446.6303 | 2 | 0.0033 | 6.63 | DYNYKLPDDFTG |
| P0DTC2 | Spike glycoprotein | 1206 | 724.3241 | 1446.6337 | 1446.6303 | 2 | 0.0033 | 8.59 | DYNYKLPDDFTG |
| P0DTC2 | Spike glycoprotein | 1206 | 724.3241 | 1446.6337 | 1446.6303 | 2 | 0.0033 | 19.89 | DYNYKLPDDFTG |
| P0DTC2 | Spike glycoprotein | 1206 | 724.3241 | 1446.6337 | 1446.6303 | 2 | 0.0034 | 23.42 | DYNYKLPDDFTG |
| P0DTC2 | Spike glycoprotein | 1206 | 724.3243 | 1446.634 | 1446.6303 | 2 | 0.0037 | 24.61 | DYNYKLPDDFTG |
| P0DTC2 | Spike glycoprotein | 1206 | 724.3244 | 1446.6343 | 1446.6303 | 2 | 0.004 | 10.23 | DYNYKLPDDFTG |
| P0DTC2 | Spike glycoprotein | 1206 | 724.3245 | 1446.6345 | 1446.6303 | 2 | 0.0041 | 11.31 | DYNYKLPDDFTG |
| P0DTC2 | Spike glycoprotein | 1206 | 724.3248 | 1446.6351 | 1446.6303 | 2 | 0.0048 | 7.13 | DYNYKLPDDFTG |
| P0DTC2 | Spike glycoprotein | 1206 | 724.325 | 1446.6354 | 1446.6303 | 2 | 0.0051 | 22.38 | DYNYKLPDDFTG |
| P0DTC2 | Spike glycoprotein | 1206 | 724.3252 | 1446.6358 | 1446.6303 | 2 | 0.0054 | 15.35 | DYNYKLPDDFTG |
| P0DTC2 | Spike glycoprotein | 1206 | 724.3253 | 1446.6361 | 1446.6303 | 2 | 0.0058 | 11.5 | DYNYKLPDDFTG |
| P0DTC2 | Spike glycoprotein | 1206 | 724.3257 | 1446.6368 | 1446.6303 | 2 | 0.0064 | 5.9 | DYNYKLPDDFTG |
| P0DTC2 | Spike glycoprotein | 1206 | 724.3257 | 1446.6369 | 1446.6303 | 2 | 0.0065 | 20.8 | DYNYKLPDDFTG |
| P0DTC2 | Spike glycoprotein | 1206 | 724.3258 | 1446.637 | 1446.6303 | 2 | 0.0066 | 8.3 | DYNYKLPDDFTG |
| P0DTC2 | Spike glycoprotein | 1206 | 490.9362 | 1469.7868 | 1469.7838 | 3 | 0.003 | 6.69 | NIQKEIDRLNEV |
| P0DTC2 | Spike glycoprotein | 1206 | 495.2598 | 1482.7577 | 1482.7539 | 3 | 0.0038 | 3.51 | RGDEVRQIAPGQTG |
| P0DTC2 | Spike glycoprotein | 1206 | 742.3862 | 1482.7578 | 1482.7539 | 2 | 0.0039 | 22.93 | RGDEVRQIAPGQTG |
| P0DTC2 | Spike glycoprotein | 1206 | 495.2599 | 1482.7578 | 1482.7539 | 3 | 0.0039 | 1.36 | RGDEVRQIAPGQTG |

| | | | | | | | | | |
|---|---|---|---|---|---|---|---|---|---|
| P0DTC2 | Spike glycoprotein | 1206 | 742.3863 | 1482.758 | 1482.7539 | 2 | 0.0041 | 12.76 | RGDEVRQIAPGQTG |
| P0DTC2 | Spike glycoprotein | 1206 | 495.26 | 1482.7582 | 1482.7539 | 3 | 0.0043 | 0.58 | RGDEVRQIAPGQTG |
| P0DTC2 | Spike glycoprotein | 1206 | 742.3869 | 1482.7592 | 1482.7539 | 2 | 0.0052 | 10.77 | RGDEVRQIAPGQTG |
| P0DTC2 | Spike glycoprotein | 1206 | 742.3873 | 1482.7601 | 1482.7539 | 2 | 0.0062 | 10.37 | RGDEVRQIAPGQTG |
| P0DTC2 | Spike glycoprotein | 1206 | 495.2607 | 1482.7603 | 1482.7539 | 3 | 0.0064 | 9.4 | RGDEVRQIAPGQTG |
| P0DTC2 | Spike glycoprotein | 1206 | 495.2609 | 1482.7608 | 1482.7539 | 3 | 0.0069 | 3.55 | RGDEVRQIAPGQTG |
| P0DTC2 | Spike glycoprotein | 1206 | 496.2644 | 1485.7714 | 1485.7728 | 3 | -0.0015 | 26.68 | RGVYYPDKVFRS |
| P0DTC2 | Spike glycoprotein | 1206 | 743.8939 | 1485.7732 | 1485.7728 | 2 | 0.0003 | 38.02 | RGVYYPDKVFRS |
| P0DTC2 | Spike glycoprotein | 1206 | 743.8939 | 1485.7733 | 1485.7728 | 2 | 0.0005 | 36.67 | RGVYYPDKVFRS |
| P0DTC2 | Spike glycoprotein | 1206 | 496.2653 | 1485.7742 | 1485.7728 | 3 | 0.0014 | 25.36 | RGVYYPDKVFRS |
| P0DTC2 | Spike glycoprotein | 1206 | 743.8955 | 1485.7765 | 1485.7728 | 2 | 0.0036 | 5.93 | RGVYYPDKVFRS |
| P0DTC2 | Spike glycoprotein | 1206 | 743.8957 | 1485.7769 | 1485.7728 | 2 | 0.004 | 31.42 | RGVYYPDKVFRS |
| P0DTC2 | Spike glycoprotein | 1206 | 496.2663 | 1485.777 | 1485.7728 | 3 | 0.0042 | 24.82 | RGVYYPDKVFRS |
| P0DTC2 | Spike glycoprotein | 1206 | 496.2663 | 1485.7771 | 1485.7728 | 3 | 0.0043 | 22.86 | RGVYYPDKVFRS |
| P0DTC2 | Spike glycoprotein | 1206 | 496.2666 | 1485.7779 | 1485.7728 | 3 | 0.0051 | 16.86 | RGVYYPDKVFRS |
| P0DTC2 | Spike glycoprotein | 1206 | 496.2666 | 1485.7779 | 1485.7728 | 3 | 0.0051 | 17.76 | RGVYYPDKVFRS |
| P0DTC2 | Spike glycoprotein | 1206 | 496.2666 | 1485.7781 | 1485.7728 | 3 | 0.0052 | 11.66 | RGVYYPDKVFRS |
| P0DTC2 | Spike glycoprotein | 1206 | 496.2667 | 1485.7782 | 1485.7728 | 3 | 0.0054 | 18.44 | RGVYYPDKVFRS |
| P0DTC2 | Spike glycoprotein | 1206 | 496.2668 | 1485.7785 | 1485.7728 | 3 | 0.0056 | 17.98 | RGVYYPDKVFRS |
| P0DTC2 | Spike glycoprotein | 1206 | 496.2669 | 1485.7789 | 1485.7728 | 3 | 0.0061 | 14.93 | RGVYYPDKVFRS |
| P0DTC2 | Spike glycoprotein | 1206 | 496.2669 | 1485.779 | 1485.7728 | 3 | 0.0061 | 18.13 | RGVYYPDKVFRS |

| | | | | | | | | | |
|---|---|---|---|---|---|---|---|---|---|
| P0DTC2 | Spike glycoprotein | 1206 | 496.2669 | 1485.779 | 1485.7728 | 3 | 0.0062 | 19.65 | RGVYYPDKVFRS |
| P0DTC2 | Spike glycoprotein | 1206 | 496.267 | 1485.7791 | 1485.7728 | 3 | 0.0062 | 17.38 | RGVYYPDKVFRS |
| P0DTC2 | Spike glycoprotein | 1206 | 496.267 | 1485.7791 | 1485.7728 | 3 | 0.0062 | 10.94 | RGVYYPDKVFRS |
| P0DTC2 | Spike glycoprotein | 1206 | 496.267 | 1485.7791 | 1485.7728 | 3 | 0.0063 | 18.14 | RGVYYPDKVFRS |
| P0DTC2 | Spike glycoprotein | 1206 | 496.2671 | 1485.7794 | 1485.7728 | 3 | 0.0066 | 19.43 | RGVYYPDKVFRS |
| P0DTC2 | Spike glycoprotein | 1206 | 496.2673 | 1485.78 | 1485.7728 | 3 | 0.0071 | 17.7 | RGVYYPDKVFRS |
| P0DTC2 | Spike glycoprotein | 1206 | 496.2673 | 1485.7802 | 1485.7728 | 3 | 0.0074 | 20.27 | RGVYYPDKVFRS |
| P0DTC2 | Spike glycoprotein | 1206 | 496.2674 | 1485.7804 | 1485.7728 | 3 | 0.0076 | 14.18 | RGVYYPDKVFRS |
| P0DTC2 | Spike glycoprotein | 1206 | 743.8981 | 1485.7816 | 1485.7728 | 2 | 0.0087 | 17.6 | RGVYYPDKVFRS |
| P0DTC2 | Spike glycoprotein | 1206 | 496.2678 | 1485.7816 | 1485.7728 | 3 | 0.0087 | 19.71 | RGVYYPDKVFRS |
| P0DTC2 | Spike glycoprotein | 1206 | 496.2686 | 1485.784 | 1485.7728 | 3 | 0.0111 | 19.98 | RGVYYPDKVFRS |
| P0DTC2 | Spike glycoprotein | 1206 | 747.8861 | 1493.7576 | 1493.7554 | 2 | 0.0021 | 14.89 | FKNIDGYFKIYS |
| P0DTC2 | Spike glycoprotein | 1206 | 498.9266 | 1493.7579 | 1493.7554 | 3 | 0.0024 | 4.16 | FKNIDGYFKIYS |
| P0DTC2 | Spike glycoprotein | 1206 | 498.9268 | 1493.7586 | 1493.7554 | 3 | 0.0032 | 15.28 | FKNIDGYFKIYS |
| P0DTC2 | Spike glycoprotein | 1206 | 498.9269 | 1493.7589 | 1493.7554 | 3 | 0.0035 | 24.78 | FKNIDGYFKIYS |
| P0DTC2 | Spike glycoprotein | 1206 | 498.9271 | 1493.7594 | 1493.7554 | 3 | 0.004 | 6.22 | FKNIDGYFKIYS |
| P0DTC2 | Spike glycoprotein | 1206 | 767.3063 | 1532.598 | 1532.6065 | 2 | -0.0084 | 5.61 | CEFQFCNDPFLG |
| P0DTC2 | Spike glycoprotein | 1206 | 767.3099 | 1532.6052 | 1532.6065 | 2 | -0.0012 | 6.43 | CEFQFCNDPFLG |
| P0DTC2 | Spike glycoprotein | 1206 | 767.3106 | 1532.6067 | 1532.6065 | 2 | 0.0003 | 11.9 | CEFQFCNDPFLG |
| P0DTC2 | Spike glycoprotein | 1206 | 767.3108 | 1532.607 | 1532.6065 | 2 | 0.0005 | 8.41 | CEFQFCNDPFLG |
| P0DTC2 | Spike glycoprotein | 1206 | 767.3108 | 1532.607 | 1532.6065 | 2 | 0.0005 | 5.9 | CEFQFCNDPFLG |

| | | | | | | | | | |
|---|---|---|---|---|---|---|---|---|---|
| P0DTC2 | Spike glycoprotein | 1206 | 767.3115 | 1532.6084 | 1532.6065 | 2 | 0.0019 | 25.18 | CEFQFCNDPFLG |
| P0DTC2 | Spike glycoprotein | 1206 | 767.3122 | 1532.6098 | 1532.6065 | 2 | 0.0033 | 16.24 | CEFQFCNDPFLG |
| P0DTC2 | Spike glycoprotein | 1206 | 767.3123 | 1532.6101 | 1532.6065 | 2 | 0.0036 | 25.17 | CEFQFCNDPFLG |
| P0DTC2 | Spike glycoprotein | 1206 | 767.3124 | 1532.6102 | 1532.6065 | 2 | 0.0037 | 4.66 | CEFQFCNDPFLG |
| P0DTC2 | Spike glycoprotein | 1206 | 767.3124 | 1532.6103 | 1532.6065 | 2 | 0.0038 | 0.47 | CEFQFCNDPFLG |
| P0DTC2 | Spike glycoprotein | 1206 | 767.3124 | 1532.6103 | 1532.6065 | 2 | 0.0039 | 9.78 | CEFQFCNDPFLG |
| P0DTC2 | Spike glycoprotein | 1206 | 767.3125 | 1532.6105 | 1532.6065 | 2 | 0.004 | 14.55 | CEFQFCNDPFLG |
| P0DTC2 | Spike glycoprotein | 1206 | 767.3129 | 1532.6113 | 1532.6065 | 2 | 0.0048 | 15.17 | CEFQFCNDPFLG |
| P0DTC2 | Spike glycoprotein | 1206 | 767.3133 | 1532.612 | 1532.6065 | 2 | 0.0056 | 5.16 | CEFQFCNDPFLG |
| P0DTC2 | Spike glycoprotein | 1206 | 767.3134 | 1532.6123 | 1532.6065 | 2 | 0.0059 | 11.03 | CEFQFCNDPFLG |
| P0DTC2 | Spike glycoprotein | 1206 | 767.3145 | 1532.6144 | 1532.6065 | 2 | 0.0079 | 43.41 | CEFQFCNDPFLG |
| P0DTC2 | Spike glycoprotein | 1206 | 771.4188 | 1540.823 | 1540.8249 | 2 | -0.002 | 10.27 | RDLPQGFSALEPLV |
| P0DTC2 | Spike glycoprotein | 1206 | 777.3876 | 1552.7606 | 1552.7674 | 2 | -0.0068 | 14 | FLPFQQFGRDIAD |
| P0DTC2 | Spike glycoprotein | 1206 | 525.2757 | 1572.8052 | 1572.8049 | 3 | 0.0004 | 30.86 | RGVYYPDKVFRSS |
| P0DTC2 | Spike glycoprotein | 1206 | 394.209 | 1572.8067 | 1572.8049 | 4 | 0.0018 | 4.15 | RGVYYPDKVFRSS |
| P0DTC2 | Spike glycoprotein | 1206 | 525.2765 | 1572.8076 | 1572.8049 | 3 | 0.0028 | 31.44 | RGVYYPDKVFRSS |
| P0DTC2 | Spike glycoprotein | 1206 | 394.2102 | 1572.8119 | 1572.8049 | 4 | 0.007 | 13.52 | RGVYYPDKVFRSS |
| P0DTC2 | Spike glycoprotein | 1206 | 792.3939 | 1582.7732 | 1582.7627 | 2 | 0.0105 | 37.33 | NVYADSFVIRGDEV |
| P0DTC2 | Spike glycoprotein | 1206 | 792.3954 | 1582.7762 | 1582.7627 | 2 | 0.0135 | 49.88 | NVYADSFVIRGDEV |
| P0DTC2 | Spike glycoprotein | 1206 | 814.3946 | 1626.7746 | 1626.7712 | 2 | 0.0034 | 25.17 | LADAGFIKQYGDCLG |
| P0DTC2 | Spike glycoprotein | 1206 | 814.3953 | 1626.7761 | 1626.7712 | 2 | 0.0049 | 11.16 | LADAGFIKQYGDCLG |

| | | | | | | | | | |
|---|---|---|---|---|---|---|---|---|---|
| P0DTC2 | Spike glycoprotein | 1206 | 814.3974 | 1626.7802 | 1626.7712 | 2 | 0.009 | 6.5 | LADAGFIKQYGDCLG |
| P0DTC2 | Spike glycoprotein | 1206 | 825.9329 | 1649.8512 | 1649.8454 | 2 | 0.0058 | 8.72 | VLHSTQDLFLPFFS |
| P0DTC2 | Spike glycoprotein | 1206 | 827.914 | 1653.8135 | 1653.8151 | 2 | -0.0016 | 11.55 | FLPFQQFGRDIADT |
| P0DTC2 | Spike glycoprotein | 1206 | 827.9172 | 1653.8198 | 1653.8151 | 2 | 0.0047 | 16.66 | FLPFQQFGRDIADT |
| P0DTC2 | Spike glycoprotein | 1206 | 849.9515 | 1697.8885 | 1697.8836 | 2 | 0.0049 | 14.5 | DAVRDPQTLEILDIT |
| P0DTC2 | Spike glycoprotein | 1206 | 849.952 | 1697.8894 | 1697.8836 | 2 | 0.0058 | 33.12 | DAVRDPQTLEILDIT |
| P0DTC2 | Spike glycoprotein | 1206 | 570.6585 | 1708.9537 | 1708.9512 | 3 | 0.0026 | 23.12 | KRSFIEDLLFNKVT |
| P0DTC2 | Spike glycoprotein | 1206 | 855.4842 | 1708.9538 | 1708.9512 | 2 | 0.0027 | 18.6 | KRSFIEDLLFNKVT |
| P0DTC2 | Spike glycoprotein | 1206 | 855.4845 | 1708.9544 | 1708.9512 | 2 | 0.0033 | 19.26 | KRSFIEDLLFNKVT |
| P0DTC2 | Spike glycoprotein | 1206 | 570.6592 | 1708.9558 | 1708.9512 | 3 | 0.0046 | 29.45 | KRSFIEDLLFNKVT |
| P0DTC2 | Spike glycoprotein | 1206 | 570.6594 | 1708.9562 | 1708.9512 | 3 | 0.005 | 39.73 | KRSFIEDLLFNKVT |
| P0DTC2 | Spike glycoprotein | 1206 | 855.4854 | 1708.9563 | 1708.9512 | 2 | 0.0051 | 10.41 | KRSFIEDLLFNKVT |
| P0DTC2 | Spike glycoprotein | 1206 | 570.6597 | 1708.9572 | 1708.9512 | 3 | 0.006 | 14.47 | KRSFIEDLLFNKVT |
| P0DTC2 | Spike glycoprotein | 1206 | 570.6597 | 1708.9574 | 1708.9512 | 3 | 0.0062 | 18.76 | KRSFIEDLLFNKVT |
| P0DTC2 | Spike glycoprotein | 1206 | 570.6599 | 1708.9579 | 1708.9512 | 3 | 0.0067 | 30.58 | KRSFIEDLLFNKVT |
| P0DTC2 | Spike glycoprotein | 1206 | 570.6601 | 1708.9584 | 1708.9512 | 3 | 0.0073 | 8.56 | KRSFIEDLLFNKVT |
| P0DTC2 | Spike glycoprotein | 1206 | 584.3068 | 1749.8984 | 1749.8951 | 3 | 0.0034 | 13.49 | GGNYNYLYRLFRKS |
| P0DTC2 | Spike glycoprotein | 1206 | 438.482 | 1749.8991 | 1749.8951 | 4 | 0.004 | 9.34 | GGNYNYLYRLFRKS |
| P0DTC2 | Spike glycoprotein | 1206 | 438.4821 | 1749.8993 | 1749.8951 | 4 | 0.0043 | 5.24 | GGNYNYLYRLFRKS |
| P0DTC2 | Spike glycoprotein | 1206 | 584.3071 | 1749.8996 | 1749.8951 | 3 | 0.0045 | 6.11 | GGNYNYLYRLFRKS |
| P0DTC2 | Spike glycoprotein | 1206 | 438.4822 | 1749.8996 | 1749.8951 | 4 | 0.0045 | 9.33 | GGNYNYLYRLFRKS |

| | | | | | | | | | |
|---|---|---|---|---|---|---|---|---|---|
| P0DTC2 | Spike glycoprotein | 1206 | 584.3073 | 1749.9002 | 1749.8951 | 3 | 0.0051 | 17.08 | GGNYNYLYRLFRKS |
| P0DTC2 | Spike glycoprotein | 1206 | 438.4823 | 1749.9003 | 1749.8951 | 4 | 0.0052 | 5.31 | GGNYNYLYRLFRKS |
| P0DTC2 | Spike glycoprotein | 1206 | 438.4825 | 1749.9007 | 1749.8951 | 4 | 0.0056 | 3.11 | GGNYNYLYRLFRKS |
| P0DTC2 | Spike glycoprotein | 1206 | 438.4826 | 1749.9014 | 1749.8951 | 4 | 0.0064 | 4.01 | GGNYNYLYRLFRKS |
| P0DTC2 | Spike glycoprotein | 1206 | 878.4343 | 1754.8541 | 1754.8628 | 2 | -0.0087 | 5.02 | FLPFQQFGRDIADTT |
| P0DTC2 | Spike glycoprotein | 1206 | 878.438 | 1754.8615 | 1754.8628 | 2 | -0.0013 | 13.45 | FLPFQQFGRDIADTT |
| P0DTC2 | Spike glycoprotein | 1206 | 878.4381 | 1754.8616 | 1754.8628 | 2 | -0.0012 | 9.9 | FLPFQQFGRDIADTT |
| P0DTC2 | Spike glycoprotein | 1206 | 878.4385 | 1754.8625 | 1754.8628 | 2 | -0.0003 | 24.61 | FLPFQQFGRDIADTT |
| P0DTC2 | Spike glycoprotein | 1206 | 878.4385 | 1754.8625 | 1754.8628 | 2 | -0.0003 | 18.26 | FLPFQQFGRDIADTT |
| P0DTC2 | Spike glycoprotein | 1206 | 878.4387 | 1754.8628 | 1754.8628 | 2 | 0 | 18.82 | FLPFQQFGRDIADTT |
| P0DTC2 | Spike glycoprotein | 1206 | 878.4387 | 1754.8628 | 1754.8628 | 2 | 0 | 22.8 | FLPFQQFGRDIADTT |
| P0DTC2 | Spike glycoprotein | 1206 | 878.4389 | 1754.8632 | 1754.8628 | 2 | 0.0004 | 20.51 | FLPFQQFGRDIADTT |
| P0DTC2 | Spike glycoprotein | 1206 | 878.4394 | 1754.8643 | 1754.8628 | 2 | 0.0014 | 13.63 | FLPFQQFGRDIADTT |
| P0DTC2 | Spike glycoprotein | 1206 | 878.4395 | 1754.8644 | 1754.8628 | 2 | 0.0016 | 20.68 | FLPFQQFGRDIADTT |
| P0DTC2 | Spike glycoprotein | 1206 | 878.4402 | 1754.8659 | 1754.8628 | 2 | 0.0031 | 16.34 | FLPFQQFGRDIADTT |
| P0DTC2 | Spike glycoprotein | 1206 | 878.4407 | 1754.8669 | 1754.8628 | 2 | 0.0041 | 16.32 | FLPFQQFGRDIADTT |
| P0DTC2 | Spike glycoprotein | 1206 | 878.4408 | 1754.8671 | 1754.8628 | 2 | 0.0043 | 14.97 | FLPFQQFGRDIADTT |
| P0DTC2 | Spike glycoprotein | 1206 | 878.4411 | 1754.8676 | 1754.8628 | 2 | 0.0048 | 17.92 | FLPFQQFGRDIADTT |
| P0DTC2 | Spike glycoprotein | 1206 | 878.4412 | 1754.8679 | 1754.8628 | 2 | 0.0051 | 19.96 | FLPFQQFGRDIADTT |
| P0DTC2 | Spike glycoprotein | 1206 | 878.4415 | 1754.8684 | 1754.8628 | 2 | 0.0056 | 20.59 | FLPFQQFGRDIADTT |
| P0DTC2 | Spike glycoprotein | 1206 | 878.4415 | 1754.8685 | 1754.8628 | 2 | 0.0057 | 16.67 | FLPFQQFGRDIADTT |

| | | | | | | | | | |
|---|---|---|---|---|---|---|---|---|---|
| P0DTC2 | Spike glycoprotein | 1206 | 878.4417 | 1754.8689 | 1754.8628 | 2 | 0.0061 | 20.43 | FLPFQQFGRDIADTT |
| P0DTC2 | Spike glycoprotein | 1206 | 878.4424 | 1754.8703 | 1754.8628 | 2 | 0.0075 | 16.6 | FLPFQQFGRDIADTT |
| P0DTC2 | Spike glycoprotein | 1206 | 878.4424 | 1754.8703 | 1754.8628 | 2 | 0.0075 | 22.73 | FLPFQQFGRDIADTT |
| P0DTC2 | Spike glycoprotein | 1206 | 878.443 | 1754.8715 | 1754.8628 | 2 | 0.0087 | 15.26 | FLPFQQFGRDIADTT |
| P0DTC2 | Spike glycoprotein | 1206 | 884.985 | 1767.9554 | 1767.9519 | 2 | 0.0035 | 31.1 | NLVRDLPQGFSALEPL |
| P0DTC2 | Spike glycoprotein | 1206 | 884.9866 | 1767.9587 | 1767.9519 | 2 | 0.0068 | 9.81 | NLVRDLPQGFSALEPL |
| P0DTC2 | Spike glycoprotein | 1206 | 899.9325 | 1797.8505 | 1797.8396 | 2 | 0.0109 | 31.55 | NYKLPDDFTGCVIAW |
| P0DTC2 | Spike glycoprotein | 1206 | 453.0009 | 1807.9745 | 1807.9733 | 4 | 0.0012 | 1.26 | NKKFLPFQQFGRDIA |
| P0DTC2 | Spike glycoprotein | 1206 | 453.0014 | 1807.9763 | 1807.9733 | 4 | 0.003 | 12.5 | NKKFLPFQQFGRDIA |
| P0DTC2 | Spike glycoprotein | 1206 | 603.6663 | 1807.9771 | 1807.9733 | 3 | 0.0038 | 33.49 | NKKFLPFQQFGRDIA |
| P0DTC2 | Spike glycoprotein | 1206 | 453.0016 | 1807.9772 | 1807.9733 | 4 | 0.0039 | 9.06 | NKKFLPFQQFGRDIA |
| P0DTC2 | Spike glycoprotein | 1206 | 603.6664 | 1807.9773 | 1807.9733 | 3 | 0.004 | 30.59 | NKKFLPFQQFGRDIA |
| P0DTC2 | Spike glycoprotein | 1206 | 453.0016 | 1807.9774 | 1807.9733 | 4 | 0.0041 | 10.26 | NKKFLPFQQFGRDIA |
| P0DTC2 | Spike glycoprotein | 1206 | 603.6665 | 1807.9777 | 1807.9733 | 3 | 0.0043 | 19.27 | NKKFLPFQQFGRDIA |
| P0DTC2 | Spike glycoprotein | 1206 | 453.0018 | 1807.9781 | 1807.9733 | 4 | 0.0047 | 9.02 | NKKFLPFQQFGRDIA |
| P0DTC2 | Spike glycoprotein | 1206 | 603.6667 | 1807.9782 | 1807.9733 | 3 | 0.0048 | 30.45 | NKKFLPFQQFGRDIA |
| P0DTC2 | Spike glycoprotein | 1206 | 603.6668 | 1807.9785 | 1807.9733 | 3 | 0.0052 | 28.43 | NKKFLPFQQFGRDIA |
| P0DTC2 | Spike glycoprotein | 1206 | 603.6669 | 1807.9787 | 1807.9733 | 3 | 0.0054 | 7.81 | NKKFLPFQQFGRDIA |
| P0DTC2 | Spike glycoprotein | 1206 | 603.6669 | 1807.979 | 1807.9733 | 3 | 0.0056 | 21.69 | NKKFLPFQQFGRDIA |
| P0DTC2 | Spike glycoprotein | 1206 | 453.0021 | 1807.9793 | 1807.9733 | 4 | 0.006 | 12.04 | NKKFLPFQQFGRDIA |
| P0DTC2 | Spike glycoprotein | 1206 | 603.6672 | 1807.9799 | 1807.9733 | 3 | 0.0066 | 21.26 | NKKFLPFQQFGRDIA |

| | | | | | | | | | |
|---|---|---|---|---|---|---|---|---|---|
| P0DTC2 | Spike glycoprotein | 1206 | 904.9976 | 1807.9806 | 1807.9733 | 2 | 0.0073 | 3.03 | NKKFLPFQQFGRDIA |
| P0DTC2 | Spike glycoprotein | 1206 | 453.0037 | 1807.9857 | 1807.9733 | 4 | 0.0123 | 8.63 | NKKFLPFQQFGRDIA |
| P0DTC2 | Spike glycoprotein | 1206 | 937.4348 | 1872.855 | 1872.8539 | 2 | 0.0011 | 12.2 | IKVCEFQFCNDPFLG |
| P0DTC2 | Spike glycoprotein | 1206 | 937.4355 | 1872.8564 | 1872.8539 | 2 | 0.0025 | 40.47 | IKVCEFQFCNDPFLG |
| P0DTC2 | Spike glycoprotein | 1206 | 631.9982 | 1892.9728 | 1892.9744 | 3 | -0.0016 | 33.51 | DLEGKQGNFKNLREFV |
| P0DTC2 | Spike glycoprotein | 1206 | 631.9985 | 1892.9737 | 1892.9744 | 3 | -0.0008 | 29.1 | DLEGKQGNFKNLREFV |
| P0DTC2 | Spike glycoprotein | 1206 | 631.999 | 1892.975 | 1892.9744 | 3 | 0.0006 | 13.19 | DLEGKQGNFKNLREFV |
| P0DTC2 | Spike glycoprotein | 1206 | 631.9991 | 1892.9754 | 1892.9744 | 3 | 0.001 | 29.35 | DLEGKQGNFKNLREFV |
| P0DTC2 | Spike glycoprotein | 1206 | 947.4951 | 1892.9757 | 1892.9744 | 2 | 0.0013 | 31.89 | DLEGKQGNFKNLREFV |
| P0DTC2 | Spike glycoprotein | 1206 | 947.4959 | 1892.9773 | 1892.9744 | 2 | 0.0029 | 15.15 | DLEGKQGNFKNLREFV |
| P0DTC2 | Spike glycoprotein | 1206 | 632 | 1892.9782 | 1892.9744 | 3 | 0.0037 | 7.97 | DLEGKQGNFKNLREFV |
| P0DTC2 | Spike glycoprotein | 1206 | 632.0002 | 1892.9788 | 1892.9744 | 3 | 0.0043 | 62.43 | DLEGKQGNFKNLREFV |
| P0DTC2 | Spike glycoprotein | 1206 | 632.0006 | 1892.9799 | 1892.9744 | 3 | 0.0054 | 47.99 | DLEGKQGNFKNLREFV |
| P0DTC2 | Spike glycoprotein | 1206 | 947.4972 | 1892.9799 | 1892.9744 | 2 | 0.0055 | 0.22 | DLEGKQGNFKNLREFV |
| P0DTC2 | Spike glycoprotein | 1206 | 632.0007 | 1892.9802 | 1892.9744 | 3 | 0.0057 | 44.29 | DLEGKQGNFKNLREFV |
| P0DTC2 | Spike glycoprotein | 1206 | 632.0008 | 1892.9806 | 1892.9744 | 3 | 0.0061 | 33.77 | DLEGKQGNFKNLREFV |
| P0DTC2 | Spike glycoprotein | 1206 | 632.0008 | 1892.9807 | 1892.9744 | 3 | 0.0062 | 22.42 | DLEGKQGNFKNLREFV |
| P0DTC2 | Spike glycoprotein | 1206 | 632.0024 | 1892.9855 | 1892.9744 | 3 | 0.011 | 23.19 | DLEGKQGNFKNLREFV |
| P0DTC2 | Spike glycoprotein | 1206 | 675.6915 | 2024.0527 | 2024.0479 | 3 | 0.0047 | 35.31 | ESNKKFLPFQQFGRDIA |
| P0DTC2 | Spike glycoprotein | 1206 | 675.6915 | 2024.0528 | 2024.0479 | 3 | 0.0049 | 7.25 | ESNKKFLPFQQFGRDIA |
| P0DTC2 | Spike glycoprotein | 1206 | 507.0206 | 2024.0531 | 2024.0479 | 4 | 0.0052 | 1.81 | ESNKKFLPFQQFGRDIA |

| | | | | | | | | | |
|---|---|---|---|---|---|---|---|---|---|
| P0DTC2 | Spike glycoprotein | 1206 | 507.0208 | 2024.0541 | 2024.0479 | 4 | 0.0062 | 5.6 | ESNKKFLPFQQFGRDIA |
| P0DTC2 | Spike glycoprotein | 1206 | 1019.0628 | 2036.1111 | 2036.1129 | 2 | -0.0018 | 3.46 | VVVLSFELLHAPATVCGPK |
| P0DTC2 | Spike glycoprotein | 1206 | 679.7118 | 2036.1137 | 2036.1129 | 3 | 0.0008 | 16.88 | VVVLSFELLHAPATVCGPK |
| P0DTC2 | Spike glycoprotein | 1206 | 679.7124 | 2036.1154 | 2036.1129 | 3 | 0.0025 | 13 | VVVLSFELLHAPATVCGPK |
| P0DTC2 | Spike glycoprotein | 1206 | 679.7128 | 2036.1167 | 2036.1129 | 3 | 0.0038 | 28.11 | VVVLSFELLHAPATVCGPK |
| P0DTC2 | Spike glycoprotein | 1206 | 679.7134 | 2036.1184 | 2036.1129 | 3 | 0.0055 | 14.81 | VVVLSFELLHAPATVCGPK |
| P0DTC2 | Spike glycoprotein | 1206 | 679.7137 | 2036.1194 | 2036.1129 | 3 | 0.0065 | 10.09 | VVVLSFELLHAPATVCGPK |
| P0DTC2 | Spike glycoprotein | 1206 | 679.7142 | 2036.1207 | 2036.1129 | 3 | 0.0078 | 28.65 | VVVLSFELLHAPATVCGPK |
| P0DTC2 | Spike glycoprotein | 1206 | 1019.0689 | 2036.1233 | 2036.1129 | 2 | 0.0104 | 34.15 | VVVLSFELLHAPATVCGPK |
| P0DTC2 | Spike glycoprotein | 1206 | 679.7177 | 2036.1314 | 2036.1129 | 3 | 0.0185 | 8 | VVVLSFELLHAPATVCGPK |
| P0DTC2 | Spike glycoprotein | 1206 | 1023.9986 | 2045.9827 | 2045.9881 | 2 | -0.0053 | 27.63 | LNDLCFTNVYADSFVIR |
| P0DTC2 | Spike glycoprotein | 1206 | 1023.9999 | 2045.9852 | 2045.9881 | 2 | -0.0028 | 14.23 | LNDLCFTNVYADSFVIR |
| P0DTC2 | Spike glycoprotein | 1206 | 1024.0018 | 2045.9891 | 2045.9881 | 2 | 0.001 | 12.26 | LNDLCFTNVYADSFVIR |
| P0DTC2 | Spike glycoprotein | 1206 | 1024.0022 | 2045.9898 | 2045.9881 | 2 | 0.0017 | 13.09 | LNDLCFTNVYADSFVIR |
| P0DTC2 | Spike glycoprotein | 1206 | 1024.0026 | 2045.9907 | 2045.9881 | 2 | 0.0027 | 30.41 | LNDLCFTNVYADSFVIR |
| P0DTC2 | Spike glycoprotein | 1206 | 683.0043 | 2045.991 | 2045.9881 | 3 | 0.0029 | 20.56 | LNDLCFTNVYADSFVIR |
| P0DTC2 | Spike glycoprotein | 1206 | 1024.0035 | 2045.9924 | 2045.9881 | 2 | 0.0044 | 72.45 | LNDLCFTNVYADSFVIR |
| P0DTC2 | Spike glycoprotein | 1206 | 683.0051 | 2045.9935 | 2045.9881 | 3 | 0.0055 | 27.62 | LNDLCFTNVYADSFVIR |
| P0DTC2 | Spike glycoprotein | 1206 | 1024.0041 | 2045.9936 | 2045.9881 | 2 | 0.0055 | 63.05 | LNDLCFTNVYADSFVIR |
| P0DTC2 | Spike glycoprotein | 1206 | 683.0052 | 2045.9937 | 2045.9881 | 3 | 0.0056 | 46.7 | LNDLCFTNVYADSFVIR |
| P0DTC2 | Spike glycoprotein | 1206 | 1024.0045 | 2045.9944 | 2045.9881 | 2 | 0.0063 | 51.81 | LNDLCFTNVYADSFVIR |

| | | | | | | | | | |
|---|---|---|---|---|---|---|---|---|---|
| P0DTC2 | Spike glycoprotein | 1206 | 1024.0047 | 2045.9947 | 2045.9881 | 2 | 0.0067 | 85.76 | LNDLCFTNVYADSFVIR |
| P0DTC2 | Spike glycoprotein | 1206 | 1024.005 | 2045.9954 | 2045.9881 | 2 | 0.0073 | 60.03 | LNDLCFTNVYADSFVIR |
| P0DTC2 | Spike glycoprotein | 1206 | 1024.0051 | 2045.9957 | 2045.9881 | 2 | 0.0076 | 16.47 | LNDLCFTNVYADSFVIR |
| P0DTC2 | Spike glycoprotein | 1206 | 1024.0054 | 2045.9963 | 2045.9881 | 2 | 0.0082 | 82.05 | LNDLCFTNVYADSFVIR |
| P0DTC2 | Spike glycoprotein | 1206 | 1024.0065 | 2045.9985 | 2045.9881 | 2 | 0.0105 | 49.53 | LNDLCFTNVYADSFVIR |
| P0DTC2 | Spike glycoprotein | 1206 | 709.3721 | 2125.0943 | 2125.0956 | 3 | -0.0013 | 5.64 | NKKFLPFQQFGRDIADTT |
| P0DTC2 | Spike glycoprotein | 1206 | 709.3727 | 2125.0963 | 2125.0956 | 3 | 0.0006 | 5.54 | NKKFLPFQQFGRDIADTT |
| P0DTC2 | Spike glycoprotein | 1206 | 709.3734 | 2125.0982 | 2125.0956 | 3 | 0.0026 | 9.49 | NKKFLPFQQFGRDIADTT |
| P0DTC2 | Spike glycoprotein | 1206 | 709.3734 | 2125.0984 | 2125.0956 | 3 | 0.0028 | 12.19 | NKKFLPFQQFGRDIADTT |
| P0DTC2 | Spike glycoprotein | 1206 | 709.3735 | 2125.0987 | 2125.0956 | 3 | 0.0031 | 9.95 | NKKFLPFQQFGRDIADTT |
| P0DTC2 | Spike glycoprotein | 1206 | 1063.557 | 2125.0995 | 2125.0956 | 2 | 0.0038 | 9.39 | NKKFLPFQQFGRDIADTT |
| P0DTC2 | Spike glycoprotein | 1206 | 709.3738 | 2125.0995 | 2125.0956 | 3 | 0.0039 | 14.28 | NKKFLPFQQFGRDIADTT |
| P0DTC2 | Spike glycoprotein | 1206 | 709.3738 | 2125.0997 | 2125.0956 | 3 | 0.0041 | 10.1 | NKKFLPFQQFGRDIADTT |
| P0DTC2 | Spike glycoprotein | 1206 | 709.3741 | 2125.1003 | 2125.0956 | 3 | 0.0047 | 3.87 | NKKFLPFQQFGRDIADTT |
| P0DTC2 | Spike glycoprotein | 1206 | 709.3742 | 2125.1007 | 2125.0956 | 3 | 0.0051 | 12.25 | NKKFLPFQQFGRDIADTT |
| P0DTC2 | Spike glycoprotein | 1206 | 709.3744 | 2125.1013 | 2125.0956 | 3 | 0.0056 | 11.23 | NKKFLPFQQFGRDIADTT |
| P0DTC2 | Spike glycoprotein | 1206 | 709.3744 | 2125.1013 | 2125.0956 | 3 | 0.0056 | 8.9 | NKKFLPFQQFGRDIADTT |
| P0DTC2 | Spike glycoprotein | 1206 | 1063.5584 | 2125.1023 | 2125.0956 | 2 | 0.0066 | 10.88 | NKKFLPFQQFGRDIADTT |
| P0DTC2 | Spike glycoprotein | 1206 | 1063.5584 | 2125.1023 | 2125.0956 | 2 | 0.0067 | 5.4 | NKKFLPFQQFGRDIADTT |
| P0DTC2 | Spike glycoprotein | 1206 | 709.3747 | 2125.1024 | 2125.0956 | 3 | 0.0067 | 9.82 | NKKFLPFQQFGRDIADTT |
| P0DTC2 | Spike glycoprotein | 1206 | 709.3748 | 2125.1027 | 2125.0956 | 3 | 0.0071 | 4.97 | NKKFLPFQQFGRDIADTT |

| | | | | | | | | | |
|---|---|---|---|---|---|---|---|---|---|
| P0DTC2 | Spike glycoprotein | 1206 | 709.375 | 2125.1031 | 2125.0956 | 3 | 0.0075 | 5.26 | NKKFLPFQQFGRDIADTT |
| P0DTC2 | Spike glycoprotein | 1206 | 1063.5591 | 2125.1036 | 2125.0956 | 2 | 0.008 | 6.14 | NKKFLPFQQFGRDIADTT |
| P0DTC2 | Spike glycoprotein | 1206 | 1063.5618 | 2125.109 | 2125.0956 | 2 | 0.0133 | 7.04 | NKKFLPFQQFGRDIADTT |
| P0DTC2 | Spike glycoprotein | 1206 | 759.6968 | 2276.0687 | 2276.061 | 3 | 0.0077 | 8.75 | WNSNNLDSKVGGNYNYLYR |
| P0DTC2 | Spike glycoprotein | 1206 | 772.7168 | 2315.1284 | 2315.1223 | 3 | 0.0062 | 38.86 | RFDNPVLPFNDGVYFASTEK |
| P0DTC2 | Spike glycoprotein | 1206 | 772.7172 | 2315.1299 | 2315.1223 | 3 | 0.0076 | 29.78 | RFDNPVLPFNDGVYFASTEK |
| P0DTC2 | Spike glycoprotein | 1206 | 775.0158 | 2322.0255 | 2322.0238 | 3 | 0.0016 | 23.1 | VCEFQFCNDPFLGVYYHK |
| P0DTC2 | Spike glycoprotein | 1206 | 781.3992 | 2341.1757 | 2341.1703 | 3 | 0.0054 | 13.22 | ESNKKFLPFQQFGRDIADTT |
| P0DTC2 | Spike glycoprotein | 1206 | 781.3994 | 2341.1764 | 2341.1703 | 3 | 0.0062 | 9.12 | ESNKKFLPFQQFGRDIADTT |
| P0DTC2 | Spike glycoprotein | 1206 | 781.4003 | 2341.1791 | 2341.1703 | 3 | 0.0088 | 9.7 | ESNKKFLPFQQFGRDIADTT |
| P0DTC2 | Spike glycoprotein | 1206 | 791.7073 | 2372.1002 | 2372.0995 | 3 | 0.0008 | 51.14 | ISNCVADYSVLYNSASFSTFK |
| P0DTC2 | Spike glycoprotein | 1206 | 1187.058 | 2372.1015 | 2372.0995 | 2 | 0.002 | 18.04 | ISNCVADYSVLYNSASFSTFK |
| P0DTC2 | Spike glycoprotein | 1206 | 791.7092 | 2372.1057 | 2372.0995 | 3 | 0.0062 | 62.65 | ISNCVADYSVLYNSASFSTFK |
| P0DTC2 | Spike glycoprotein | 1206 | 1187.0606 | 2372.1066 | 2372.0995 | 2 | 0.0072 | 28.01 | ISNCVADYSVLYNSASFSTFK |
| P0DTC2 | Spike glycoprotein | 1206 | 1187.0654 | 2372.1162 | 2372.0995 | 2 | 0.0167 | 19.25 | ISNCVADYSVLYNSASFSTFK |
| P0DTC2 | Spike glycoprotein | 1206 | 946.7849 | 2837.333 | 2837.3297 | 3 | 0.0033 | 11.52 | SYLTPGDSSSGWTAGAAAYYVGYLQPR |
| P0DTC2 | Spike glycoprotein | 1206 | 727.8684 | 2907.4445 | 2907.4416 | 4 | 0.0029 | 9.26 | WNSNNLDSKVGGNYNYLYRLFRKS |
| P0DTC2 | Spike glycoprotein | 1206 | 727.8691 | 2907.4472 | 2907.4416 | 4 | 0.0057 | 7.77 | WNSNNLDSKVGGNYNYLYRLFRKS |
| P0DTC2 | Spike glycoprotein | 1206 | 1141.5609 | 3421.6608 | 3421.646 | 3 | 0.0148 | 2.71 | QQFGRDIADTTDAVRDPQTLEILDITPCSF |
| P0DTC9 | Nucleoprotein | 105 | 398.715 | 795.4155 | 795.4127 | 2 | 0.0029 | 30.23 | TKAYNVT |
| P0DTC9 | Nucleoprotein | 105 | 420.7568 | 839.499 | 839.4977 | 2 | 0.0013 | 9.55 | ALPQRQK |
| P0DTC9 | Nucleoprotein | 105 | 429.2159 | 856.4173 | 856.4151 | 2 | 0.0022 | 8.35 | TRNPANNA |

| | | | | | | | | | |
|---|---|---|---|---|---|---|---|---|---|
| P0DTC9 | Nucleoprotein | 105 | 485.309 | 968.6035 | 968.6018 | 2 | 0.0016 | 14.33 | LLLLDRLN |
| P0DTC9 | Nucleoprotein | 105 | 485.3093 | 968.6041 | 968.6018 | 2 | 0.0022 | 34.54 | LLLLDRLN |
| P0DTC9 | Nucleoprotein | 105 | 485.3094 | 968.6043 | 968.6018 | 2 | 0.0025 | 30.64 | LLLLDRLN |
| P0DTC9 | Nucleoprotein | 105 | 485.3096 | 968.6047 | 968.6018 | 2 | 0.0029 | 34.65 | LLLLDRLN |
| P0DTC9 | Nucleoprotein | 105 | 485.3098 | 968.6051 | 968.6018 | 2 | 0.0033 | 51.7 | LLLLDRLN |
| P0DTC9 | Nucleoprotein | 105 | 485.31 | 968.6055 | 968.6018 | 2 | 0.0036 | 18.08 | LLLLDRLN |
| P0DTC9 | Nucleoprotein | 105 | 485.31 | 968.6055 | 968.6018 | 2 | 0.0036 | 33.14 | LLLLDRLN |
| P0DTC9 | Nucleoprotein | 105 | 485.3118 | 968.609 | 968.6018 | 2 | 0.0071 | 34.39 | LLLLDRLN |
| P0DTC9 | Nucleoprotein | 105 | 548.8332 | 1095.6518 | 1095.6512 | 2 | 0.0006 | 12.43 | ALPQRQKKQ |
| P0DTC9 | Nucleoprotein | 105 | 579.7976 | 1157.5807 | 1157.5789 | 2 | 0.0018 | 28.02 | GKGQQQQGQTV |
| P0DTC9 | Nucleoprotein | 105 | 581.7812 | 1161.5479 | 1161.5486 | 2 | -0.0007 | 6.3 | SRNSSRNSTPG |
| P0DTC9 | Nucleoprotein | 105 | 400.5829 | 1198.727 | 1198.7258 | 3 | 0.0012 | 3.69 | SKKPRQKRTA |
| P0DTC9 | Nucleoprotein | 105 | 623.3136 | 1244.6127 | 1244.6109 | 2 | 0.0018 | 11.25 | SGKGQQQQGQTV |
| P0DTC9 | Nucleoprotein | 105 | 626.3211 | 1250.6277 | 1250.6255 | 2 | 0.0022 | 1.73 | EGALNTPKDHIG |
| P0DTC9 | Nucleoprotein | 105 | 417.884 | 1250.6301 | 1250.6255 | 3 | 0.0046 | 12.09 | EGALNTPKDHIG |
| P0DTC9 | Nucleoprotein | 105 | 626.3225 | 1250.6304 | 1250.6255 | 2 | 0.0049 | 15.61 | EGALNTPKDHIG |
| P0DTC9 | Nucleoprotein | 105 | 630.3231 | 1258.6316 | 1258.6266 | 2 | 0.005 | 22.06 | GKGQQQQGQTVT |
| P0DTC9 | Nucleoprotein | 105 | 663.3875 | 1324.7605 | 1324.7575 | 2 | 0.003 | 6.01 | ALPQRQKKQQT |
| P0DTC9 | Nucleoprotein | 105 | 673.8383 | 1345.6621 | 1345.6586 | 2 | 0.0035 | 20.39 | SGKGQQQQGQTVT |
| P0DTC9 | Nucleoprotein | 105 | 676.8458 | 1351.677 | 1351.6732 | 2 | 0.0038 | 2.4 | TEGALNTPKDHIG |
| P0DTC9 | Nucleoprotein | 105 | 752.8825 | 1503.7504 | 1503.7464 | 2 | 0.004 | 31.46 | KMSGKGQQQQGQTV |
| P0DTC9 | Nucleoprotein | 105 | 502.2576 | 1503.751 | 1503.7464 | 3 | 0.0047 | 15.79 | KMSGKGQQQQGQTV |
| P0DTC9 | Nucleoprotein | 105 | 781.8805 | 1561.7465 | 1561.7485 | 2 | -0.002 | 65.29 | QGNFGDQELIRQGT |
| P0DTC9 | Nucleoprotein | 105 | 781.884 | 1561.7535 | 1561.7485 | 2 | 0.005 | 65.61 | QGNFGDQELIRQGT |
| P0DTC9 | Nucleoprotein | 105 | 781.8857 | 1561.7569 | 1561.7485 | 2 | 0.0084 | 25.53 | QGNFGDQELIRQGT |
| P0DTC9 | Nucleoprotein | 105 | 535.9401 | 1604.7984 | 1604.7941 | 3 | 0.0043 | 9.58 | KMSGKGQQQQGQTVT |
| P0DTC9 | Nucleoprotein | 105 | 803.4067 | 1604.7988 | 1604.7941 | 2 | 0.0047 | 22.41 | KMSGKGQQQQGQTVT |
| P0DTC9 | Nucleoprotein | 105 | 600.3373 | 1797.9901 | 1797.9737 | 3 | 0.0164 | 4.66 | KEDLKFPRGQGVPINT |
| P0DTC9 | Nucleoprotein | 105 | 465.5048 | 1857.99 | 1857.9836 | 4 | 0.0064 | 6.93 | AIKLDDKDPNFKDQVI |
| P0DTC9 | Nucleoprotein | 105 | 465.5057 | 1857.9939 | 1857.9836 | 4 | 0.0103 | 0.53 | AIKLDDKDPNFKDQVI |

| | | | | | | | | | |
|---|---|---|---|---|---|---|---|---|---|
| P0DTC9 | Nucleoprotein | 105 | 779.0876 | 2334.2411 | 2334.2444 | 3 | -0.0033 | 0.92 | LTQHGKEDLKFPRGQGVPINT |
| P0DTC5 | Membrane protein | 48 | 391.2639 | 780.5132 | 780.5109 | 2 | 0.0023 | 3.59 | ILLNVPL |
| P0DTC5 | Membrane protein | 48 | 411.7576 | 821.5007 | 821.4984 | 2 | 0.0023 | 5.17 | RGHLRIA |
| P0DTC5 | Membrane protein | 48 | 457.7475 | 913.4803 | 913.477 | 2 | 0.0034 | 8.85 | YSRYRIG |
| P0DTC5 | Membrane protein | 48 | 546.2608 | 1090.507 | 1090.5043 | 2 | 0.0027 | 36.09 | NYKLNTDHS |
| P0DTC5 | Membrane protein | 48 | 546.2625 | 1090.5105 | 1090.5043 | 2 | 0.0062 | 1.73 | NYKLNTDHS |
| P0DTC5 | Membrane protein | 48 | 589.7765 | 1177.5385 | 1177.5363 | 2 | 0.0022 | 10.33 | NYKLNTDHSS |
| P0DTC5 | Membrane protein | 48 | 633.293 | 1264.5714 | 1264.5684 | 2 | 0.0031 | 18.69 | NYKLNTDHSSS |
| P0DTC5 | Membrane protein | 48 | 691.9166 | 1381.8187 | 1381.818 | 2 | 0.0006 | 33.82 | ILTRPLLESELV |
| P0DTC5 | Membrane protein | 48 | 691.9182 | 1381.8218 | 1381.818 | 2 | 0.0038 | 43.82 | ILTRPLLESELV |
| P0DTC5 | Membrane protein | 48 | 691.9183 | 1381.822 | 1381.818 | 2 | 0.004 | 2.81 | ILTRPLLESELV |
| P0DTC5 | Membrane protein | 48 | 744.3979 | 1486.7812 | 1486.7813 | 2 | -0.0001 | 15.77 | GRCDIKDLPKEIT |
| P0DTC5 | Membrane protein | 48 | 496.6018 | 1486.7834 | 1486.7813 | 3 | 0.0021 | 1.83 | GRCDIKDLPKEIT |
| P0DTC5 | Membrane protein | 48 | 496.6018 | 1486.7837 | 1486.7813 | 3 | 0.0023 | 12.64 | GRCDIKDLPKEIT |
| P0DTC5 | Membrane protein | 48 | 744.4 | 1486.7854 | 1486.7813 | 2 | 0.0041 | 33.53 | RCDIKDLPKEIT |
| P0DTC5 | Membrane protein | 48 | 496.6027 | 1486.7863 | 1486.7813 | 3 | 0.0049 | 16.48 | GRCDIKDLPKEIT |
| P0DTC5 | Membrane protein | 48 | 744.4022 | 1486.7899 | 1486.7813 | 2 | 0.0086 | 8.64 | GRCDIKDLPKEIT |
| P0DTC5 | Membrane protein | 48 | 793.9334 | 1585.8522 | 1585.8498 | 2 | 0.0024 | 5.74 | GRCDIKDLPKEITV |
| P0DTC5 | Membrane protein | 48 | 793.9337 | 1585.8529 | 1585.8498 | 2 | 0.0032 | 6.78 | GRCDIKDLPKEITV |
| P0DTC5 | Membrane protein | 48 | 397.4708 | 1585.854 | 1585.8498 | 4 | 0.0042 | 23.42 | GRCDIKDLPKEITV |

**Supplementary Table 2:** Known antibodies recognising the SARS-CoV 2 spike.

| Antibody | Binding Region | Neutralising | Reference |
|---|---|---|---|
| S2E12, S2M11 | RBD (S1) | Yes | [3] |
| 31B5, 32D4 | RBD (S1) | Yes | [4] |
| 31B9 | RBD (S1) | No | [4] |
| ADI-56046, ADI-55993, ADI-56010, ADI-55688, ADI-55689, ADI-55690, ADI-56000, ADI-55951 | RBD (S1) | Yes | [5] |
| ADI-56058, ADI-56051, ADI-55692, ADI-56040, ADI-55694, ADI-55958, ADI-56020, ADI-55754, ADI-56035 | RBD (S1) | No | [5] |
| ADI-55699, ADI-55749, ADI-56071 | RBD (S1) | Weak | [5] |
| ADI-56006 | S2 | poor | [5] |
| ADI-56032 | NTD (S1) | poor | [5] |
| CC12.1, CC12.2, CC12.3, CC12.10, CC6.29, CC6.31 | RBD (S1) | Yes | [6] |
| CC12.15, CC12.16, CC12.17, CC12.18, CC12.19 | RBD (S1) | Poor | [6] |
| P2C-1F11, P2C-1C10, P2B-2F6, P2C-1A3, P2A-1A8, P2B-2G4, P2A-1A10 | RBD (S1) | Yes | [7] |
| ACE2-Ig, mACE2-Ig | RBD (S1) | Yes | [8] |
| 4A8 | NTD (S1) | Yes | [9] |
| 0304-3H3 | S2/S-ECD (1-1208) | Yes | [9] |
| 1M-1D2 | S1 | Yes | [9] |
| 2M-7E9, 2M-8E7,0304-4A10, 2M-8H10, 2M-9H1, 2M-12D7, 2M-13A3, 2M-13D11, 2M-14E4, 1M-14E5, 9A1 | S2 | No | [9] |
| 2M-10B11 | RBD (S1) | No | [9] |
| 0304-4A2, 0317-A7 | S1 | No | [9] |
| 10C10 | S-ECD (1-1208) | No | [9] |
| CA1, CB6 | RBD (S1) | Yes | [10] |
| S309 | RBD (S1) | Yes | [11] |
| S304, S315 | RBD (S1) | Weak | [11] |
| S110 | Not clear | No | [11] |
| S124, S303 | RBD | No | [11] |
| S306, S310 | non-RBD | No | [11] |
| B38, B5, H2, H4 | RBD(S1) | Yes | [12] |
| C121, C144, C135, C002, C101, C009 | RBD (S1) | Yes | [13] |
| C105 | RBD (S1) | Yes | [14] |
| 47D11 | RBD (S1) | Yes | [15] |
| Ty1, Ty1-Fc | RBD (S1) | Yes | [16] |
| H014 | RBD (S1) | Yes | [17] |
| VHH-72 | RBD (S1) | Yes | [18] |
| CV1, CV35 | non-RBD (S2P-Ecto) | Weak | [19] |
| CV30 | RBD (S1) | Yes | [19]s |
| CV5, CV43 | RBD (S1) | No | [19] |
| CV1, CV2, CV3, CV4, CV6, CV7, CV8, CV9, CV10, CV11, CV12, CV13, CV15, CV16, CV17, CV18, CV19, CV21, CV22, CV23, CV24, CV25, CV26, CV27, CV31, CV32, CV33, CV34, CV35, CV36, CV37, CV38, CV39, CV40, CV41, CV42, CV44, CV45, CV46, CV47, CV48, CV50 | S2P | No | [19] |
| S2H14, S2H13, S2A4, S2X35 | RBD (S1) | Yes | [20] |
| REGN10989, REGN10987, REGN10933,REGN10934, REGN10964, REGN10954, REGN10984, REGN10986 | RBD (S1) | Yes | [21] |

| | | | |
|---|---|---|---|
| BD-368-2, BD-23, BD-217, BD-218, BD-236, BD-361, BD-368, BD-395, BD-503, BD-508, BD-515 | RBD (S1) | Yes | [22] |
| COV2-2196, COV2-2072, COV2-2499, COV2-3025, COV2-2381, COV2-2479, COV2-2096, COV2-2832, COV2-2050, COV2-2130, COV2-2819, COV2-2955 | RBD (S1) | Yes | [23] |
| COV2-2676, COV2-2489 | S2P-ecto | poor | [23] |
| 414-1, 505-3, 553-63, 515-1, 505-5, 553-15, 553-49, 553-60 | RBD (S1) | Yes | [24] |
| 505-8, 413-2, 515-5 | non-RBD | Yes | [24] |
| 2-15, 2-7, 1-57, 1-20, 2-38, 2-36, 4-20, 2-30, 2-4 | RBD | Yes | [25] |
| 2-17, 5-24, 4-8, 5-7,1-87, 4-19, 4-18, 1-68 | NTD (S1) | Yes | [25] |
| 2-43, | S-trimer, non-RBD, non-NTD | Yes | [25] |
| 2-51, | involves NTD (S1), not clear | No | [25] |
| Nb6, Nb11 (Nanobodies), Nb6-tri, mNb6 | RBD | Yes | [26] |
| Nb3, Nb3-tri | non-RBD (S1) | Yes | [26] |
| 2G12 | S2 | No | [27] |
| CR3022 | RBD (S1) | No | [28] |
| SR31 (sybody) | RBD (S1) | No | [29] |
| SR4, MR17, MR3, MR4 (sybody) | RBD (S1) | Yes | [30] |
| COVA1-22 | NTD (S1) | Yes | [31] |
| COVA1-18, COVA2-04, COVA2-07, COVA2-15, COVA2-17, COVA2-39, COVA2-15 | RBD (S1) | Yes | [31] |
| COVA1-21, COVA1-25 | non-RBD | Yes | [31] |
| COVA2-16 | RBD | No | [31] |
| COVA1-19,COVA2-33, COVA2-42, COVA2-10, COVA2-28, COVA2-12, COVA2-41, COVA1-19, COVA1-01, COVA1-02, COVA2-34, COVA2-47, COVA2-43, COVA2-38, COVA1-26, COVA2-25, COVA2-03, COVA2-30, COVA1-06, COVA3-07, COVA1-20, COVA2-26, COVA1-09 | S | No | [31] |
| CnC2t1p1_B4, FnC1t2p1_D4, FnC1t2p1_G5, HbnC3t1p1_C6, HbnC3t1p1_F4, HbnC3t1p1_G4, HbnC3t1p2_B10, HbnC3t1p2_C6 | RBD (S1) | Yes | [32] |
| CnC2t1p1_A2, FnC1t1p1_B6, FnC1t1p1_C11, FnC2t1p1_B1, HbnC4t1p1_A1, HbnC4t1p1_C11 | RBD (S1) | No | [32] |
| CnC2t1p1_E7, CnC2t1p2_G4, HbnC1t1p3_H8 | non-RBD (S1) | No | [32] |
| 5A6, 1F4, 2H4, 6F8, 3D11 | RBD (S1) | Yes | [33] |
| EY6A | RBD (S1) | Yes | [34] |
| CV05-163, CV07-209, CV07-222, CV07-250, CV07-262, CV38-183, CV07-283, CV07-287, CV07-315, CV38-113, CV38-139 , CV38-142 , CV38-221, CV-X1-126 | RBD (S1) | Yes | [35] |

**Supplementary videos**

Supplementary Movie 1: Mass photometry of $S1_{vaccine\ antigen}$ showing particle landing events.
Supplementary Movie 2: Mass photometry of $S_{recombinant\ trimer}$ showing particle landing events.
